# Inferences on a Multidimensional Social Hierarchy Use a Grid-like Code

**DOI:** 10.1101/2020.05.29.124651

**Authors:** Seongmin A. Park, Douglas S. Miller, Erie D. Boorman

## Abstract

Generalizing experiences to guide decision making in novel situations is a hallmark of flexible behavior. It has been hypothesized such flexibility depends on a cognitive map of an environment or task, but directly linking the two has proven elusive. Here, we find that discretely sampled abstract relationships between entities in an unseen two-dimensional (2-D) social hierarchy are reconstructed into a unitary 2-D cognitive map in the hippocampus and entorhinal cortex. We further show that humans utilize a grid-like code in several brain regions, including entorhinal cortex and medial prefrontal cortex, for *inferred* direct trajectories between entities in the reconstructed abstract space during discrete decisions. Moreover, these neural grid-like codes in the entorhinal cortex are associated with neural decision value computations in the medial prefrontal cortex and temporoparietal junction area during choice. Collectively, these findings show that grid-like codes are used by the human brain to infer novel solutions, even in abstract and discrete problems, and suggest a general mechanism underpinning flexible decision making and generalization.

---

Humans and other animals have a remarkable capacity for inferring novel solutions in situations never encountered before. This extreme flexibility sets humans in particular apart from artificial intelligence, where inferring new solutions from sparse data and flexibly generalizing knowledge across tasks remains a significant challenge (Sutton and Barto, 1998; Behrens *et al*., 2018). As early as the 1940s, it was originally proposed that animals form and use some type of a “cognitive map” of the environment that allows them to flexibly infer novel routes and shortcuts through a spatial environment to reach a known reward’s location (Tolman, 1948).

In the context of spatial navigation, so-called “place cells” in the hippocampus (HC) and “grid cells” in medial entorhinal cortex (EC) have been proposed to represent the animal’s current or predicted position in a spatiotemporal environment (O’Keefe and Nadel, 1978; Hafting *et al*., 2005; Stachenfeld, Botvinick and Gershman, 2017; Ekstrom and Ranganath, 2018) and to provide a spatial metric of that environment (Bush *et al*., 2015), respectively. Together with related cell types (e.g. border cells, band cells, etc.), these cells may provide a neuronal basis for a cognitive map of space. Critically, recent biologically inspired computational models have further proposed how these grid codes in EC may reflect a basis for statistical transitions in 2-D topologies that can in principle enable animals and artificial agents to compose new routes and find shortcuts to reach goals in spatial environments (Bush *et al*., 2015; Banino *et al*., 2018; Behrens *et al*., 2018; Whittington *et al.*, 2020). These findings suggest a novel framework for understanding how humans might effectively “shortcut” solutions in new situations.

Representing non-spatial relationships between entities or task states in the world as a cognitive map would likewise be incredibly powerful because it could theoretically be leveraged to make inferences from sparse experiences that can dramatically accelerate learning and even guide decisions never faced before (Kriete *et al*., 2013; Behrens *et al*., 2018; Wang *et al*., 2018). This is because, in theory, a cognitive map of abstract relationships, such as a conceptual space (Constantinescu, O’Reilly and Behrens, 2016), or task space (Schuck *et al*., 2016; Zhou *et al*., 2019), would allow for “novel routes” and “shortcuts” to be inferred, as in physical space (Behrens *et al*., 2018; Whittington *et al*., 2020) (**Fig.1a**). Notably, such direct inferences go beyond chaining together directly experienced associations (**Fig.1a)**, and are seen as the defining characteristic of a genuine cognitive map of spatial environments (Tolman, 1948; O’Keefe and Nadel, 1978; Bennett, 1996). Despite these theoretical insights, whether such direct inferences (as opposed to chaining together previously experienced associations) are in fact used for non-spatial decision making, and whether a grid code underpins these inferences about non-spatial relationships, is unknown.

**Figure 1.**
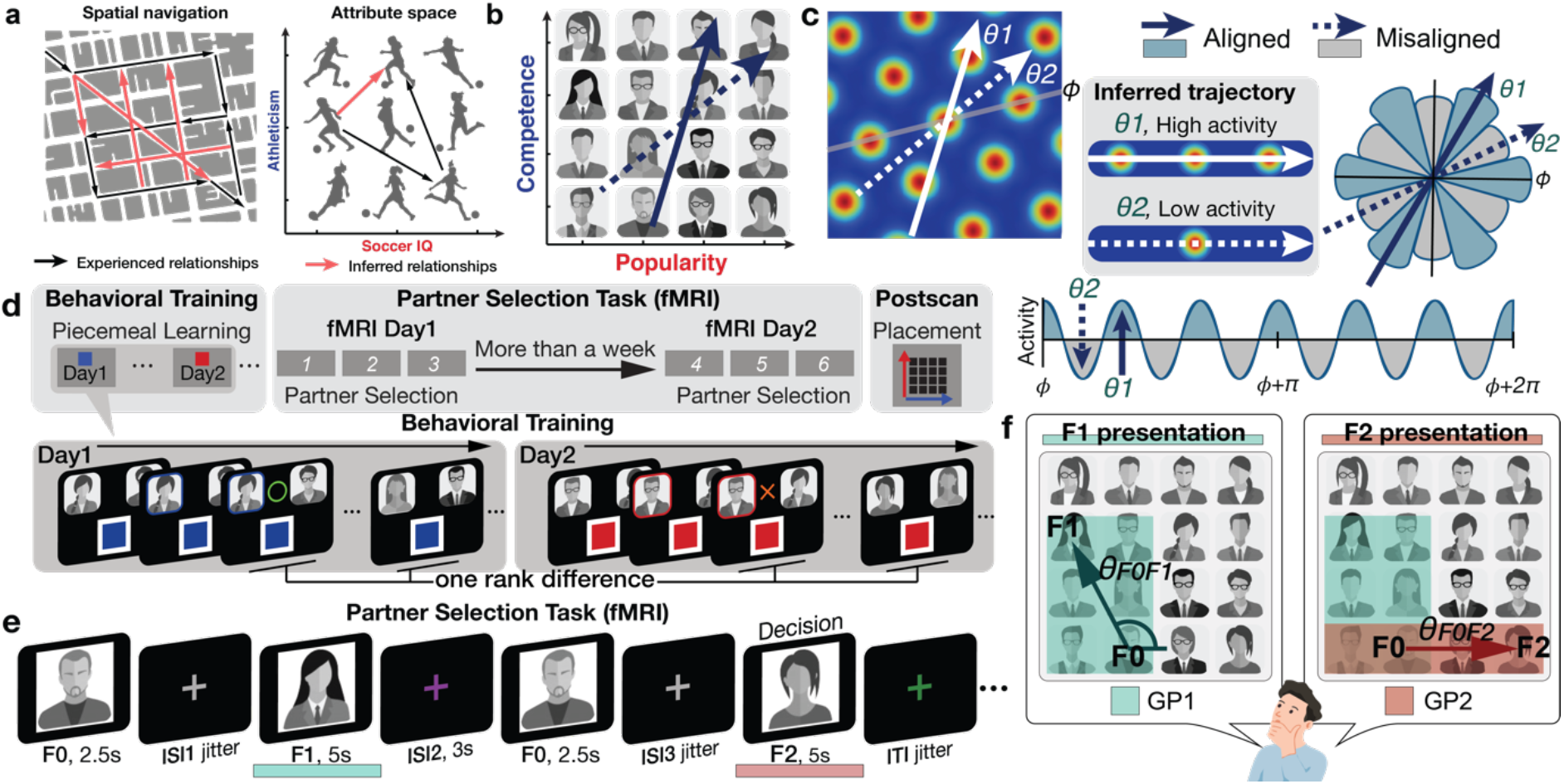
Behavioral training and experimental task. **a.** A cognitive map of spatial (left) and non-spatial (right) relational structures allows for new direct routes or relationships to be inferred from directly experienced relationships, dramatically accelerating learning and decision capabilities through generalization. **b.** Participants learned the rank of 16 individuals organized into a 2-D social hierarchy defined by competence and popularity. Participants never saw neither 1-D or 2-D social hierarchies, but they could learn it piecemeal from dyadic comparisons in one dimension at a time during behavioral training. We hypothesized neural activity would be modulated hexadirectionally by the inferred trajectories over the 2-D social space, as predicted by a hexagonal grid organization. **c.** The inferred trajectories can be categorized as aligned and misaligned with the mean grid orientation, *ϕ*, which is different for each participant. *θ*1 and *θ*2 show examples of aligned and misaligned trajectories, respectively. Greater activity is predicted when the inferred trajectory is aligned compared to misaligned because it passes over more grid fields, which generates hexadirectional grid-like modulation. **d.** Behavioral training procedure. On day 1 and day 2 of behavioral training, participants learned ranks of 16 individuals (face stimuli) in each of two dimensions (competence or popularity) through binary decisions about the higher rank individual in a pair who differed by only one rank level in a given dimension, with each dimension learned on a different day. Within a day, the order of pairs compared was further randomized. After behavioral training, participants performed 3 blocks of the “partner selection task” twice during fMRI scanning, with a gap between sessions of at least one week. After the second session, participants were asked to place individuals according to their believed combined rank in a 2-D space for the first time (placement task). **e.** Illustration of a trial of the partner selection task during fMRI. Participants were asked to make a binary decision by choosing a better business partner for a given individual (F0) between two (F1 and F2). The better partner is determined by the “growth potential (GP)” that each pair could expect from their collaboration. Participants could compute the GP of the F0 and F1 pair when F1 is shown and GP of the F0 and F2 pair when F2 is shown. Participants were subsequently asked to make a decision during F2 presentation. No feedback was given. **f.** To compute the GP of a pair, participants were instructed the GP corresponds to the higher “rank of the pair” in each dimension. Participants were further asked to weigh the ‘rank of the pair’ in both dimensions equally. Therefore, the GP corresponds to the area drawn by the higher rank of the two people in each dimension in the 2-D hierarchy (GP_F0F1_ (green rectangle) > GP_F0F2_ (red rectangle); F1 is the better partner for F0 in this example). We hypothesized people would infer direct trajectories over the mentally reconstructed 2-D space between the positions of F0 and each potential partner, F1 and F2, to compute the GP for each collaboration. We searched for neural evidence for hexadirectional modulation of inferred trajectories though the reconstructed cognitive map (*θ*_F0F1_ at the time of F1 presentation and *θ*_F0F2_ at the time of F2 presentation).

Recent pioneering studies have identified hippocampal place-like and entorhinal grid-like neural modulation for behaviorally relevant, continuous, non-spatial stimulus dimensions, including sound frequency (Aronov, Nevers and Tank, 2017), metric length of bird legs and necks in a conceptual “bird space”(Constantinescu, O’Reilly and Behrens, 2016), and odor concentration (Bao *et al*., 2019). These findings support theories proposing that place and grid cells revealed in physical space may reflect a general mechanism for organizing relationships between any behaviorally relevant dimensions (Eichenbaum and Cohen, 2014; Behrens *et al*., 2018). In the real world, a wide range of everyday decisions involve discrete entities that vary along multiple abstract dimensions whose feature values must be learned discretely from piecemeal experiences, and in the absence of explicit sensory feedback. Here, we test the hypotheses that, first, such piecemeal experiences are integrated into an abstract cognitive map in the HC–EC system, and, second, a grid code is used to make direct inferences to guide decision making based on that cognitive map (see **Fig.1b,c**).

## Results

We utilized social hierarchies to test the above hypotheses, because they represent an ecologically important example of abstracted multidimensional relationships, and build on recent studies suggesting that the brain constructs and maintains structural knowledge of social relationships (Tavares *et al*., 2015; Parkinson, Kleinbaum and Wheatley, 2017; Stolier, Hehman and Freeman, 2018; Tamir and Thornton, 2018). Specifically, we leveraged established fMRI approaches (Kriegeskorte, 2008; Doeller, Barry and Burgess, 2010; Constantinescu, O’Reilly and Behrens, 2016) to test for a reconstructed cognitive map of an abstract two-dimensional (2-D) social hierarchy, and a grid-like code for inferred direct trajectories between entities during decision making (**Fig.1b**). Importantly, the social hierarchy was never seen but could be reconstructed from the outcomes of binary comparisons about the higher rank individual between pairs on one dimension at a time, with each dimension learned on a separate day. To test our hypotheses, we designed a “partner selection” task that required participants (n=21) to infer a better collaborator for a given “entrepreneur” by comparing their relative ordinal ranks in two ecologically important social dimensions: their competence and popularity (Fiske, Cuddy and Glick, 2007). During behavioral training, participants learned the relative rank between individuals in the two independent dimensions through feedback-based binary comparisons. Participants only learned about individuals who differed by one rank level and on one dimension at a time (**Fig.1d**), with each dimension learned on a separate day (on average 2 days apart). Importantly, participants were never shown the true hierarchy, nor were they asked to construct it spatially, but they could theoretically reconstruct it through transitive inferences within a dimension on a given day (Kumaran, Melo and Duzel, 2012) and integration across dimensions learned on different days (**Fig.S1**). During fMRI participants were asked to choose a better partner for a given “entrepreneur” (F0) between two potential partners (F1 or F2) (**Fig.1e**). Critically, these individuals’ 2-D rank in the combined social space was never learned during training, meaning that each decision during the fMRI task required *inferences* to determine the better partner. That is, the two separately learned dimensions had to be combined to compute the potential of a collaboration which was never learned from feedback during training.

In our task the better collaboration was determined by the growth potential (GP) of each pair. To compute the growth potential, participants were instructed they should weight the ranks of the pair in both dimensions equally, with the rank of a pair in each dimension determined by the higher rank individual between the two. Therefore, the GP corresponds to the area drawn by the higher rank of the two entrepreneurs in each dimension in the 2-D hierarchy (**Fig.1f**). We hypothesized people would infer direct trajectories (which we refer to as *direct inferences* because they correspond to direct vectors over the 2-D space that were never learned during training) over the reconstructed 2-D space between the positions of the entrepreneur, F0, and each potential partner, F1 and F2, to compute the growth potential for each collaboration (**Fig.1b,e,f; Fig.S3d**). After training on the partner selection task, performed with an entirely different set of choice comparisons and in the absence of any choice feedback, participants who passed our performance criterion (>95% accuracy, see Methods) were highly accurate in choosing the better collaboration during the fMRI session (mean ± SE accuracy = 98.68 ± 0.27%). We found significant effects of GP on the reaction times (RT) of decisions: the greater the GP of each pair, and the absolute difference between them, the faster the RT, even after partialling out Euclidean distances of the inferred trajectories (t_20_>4.6, p<0.005; **Fig.S4**).

We first tested whether the HC-EC system would reconstruct the true unseen social hierarchy as a 2-D cognitive map. Specifically, we reasoned that more proximal people in the true 2-D social space (measured by their Euclidian distance) would be represented progressively and linearly more similarly (**Fig. 2a**). Using representational similarity analysis (RSA) applied to BOLD activity patterns evoked by each face presentation (at F1 and F2, following down-sampling to match the number of observations per face and excluding faces 1 and 16, which were sampled less frequently [see Methods and **Fig.S2**]), we found that the level of pattern dissimilarity across voxels (measured by the Mahalanobis distance) in *a priori* anatomically defined regions of interests (ROIs) in the HC (Yushkevich *et al*., 2015) and EC (Amunts *et al*., 2005; Zilles and Amunts, 2010) (**Fig.2b)** was robustly related to the pairwise Euclidian distance between individuals in the true 2D social space (p<0.001 FWE-corrected over ROIs) (**Fig.2c**), such that closer people in the true 2-D space were represented increasingly more similarly. Results were similar when instead using all samples of all 16 faces (**Fig. S5e**). To ensure these pattern similarity effects in the HC and EC were better accounted for by the 2-D Euclidian distance than either 1-D distance alone, we decomposed the Euclidian distance into separate popularity and competence dimensions. We found significant effects of the 1-D rank distance in both popularity and competence dimensions on pattern similarity in HC (popularity: τ_A_ = 0.07; competence: τ_A_ = 0.07) and EC (popularity: τ_A_ = 0.05; competence: τ_A_ = 0.04) (all p<0.001 FWE-corrected; see **Fig.2d,e** for alternative definitions of X and Y dimensions), consistent with a 2-D cognitive map. Effects of the Euclidian distance were significantly stronger than either single dimension alone (**Fig. 2c**; see **Table S6** for formal comparisons). To further probe the representational architecture in HC and EC in a 2-D space, we applied multidimensional scaling (MDS) on their activity patterns, while matching the number of observations per face, which revealed a consistent relationship between the true Euclidian distances and the estimated distances between individuals’ locations in a 2-D space (**Fig.2h; Extended Data Figure 2a).** To complement these results based on the down-sampled events, we additionally performed the RSA and MDS of the same 14 faces while including all trials’ neural responses. Consistent with the results of the sample size-matched analysis, the results of the analysis including all responses (**Extended Data Figure 1** and **2b**) and additional control analyses (**Fig.S5c,d**) confirm that the HC-EC system integrates piecemeal, discrete observations about abstract relationships, learned on separate days from the outcomes of choices, into a unitary 2-D cognitive map.

**Figure 2.**
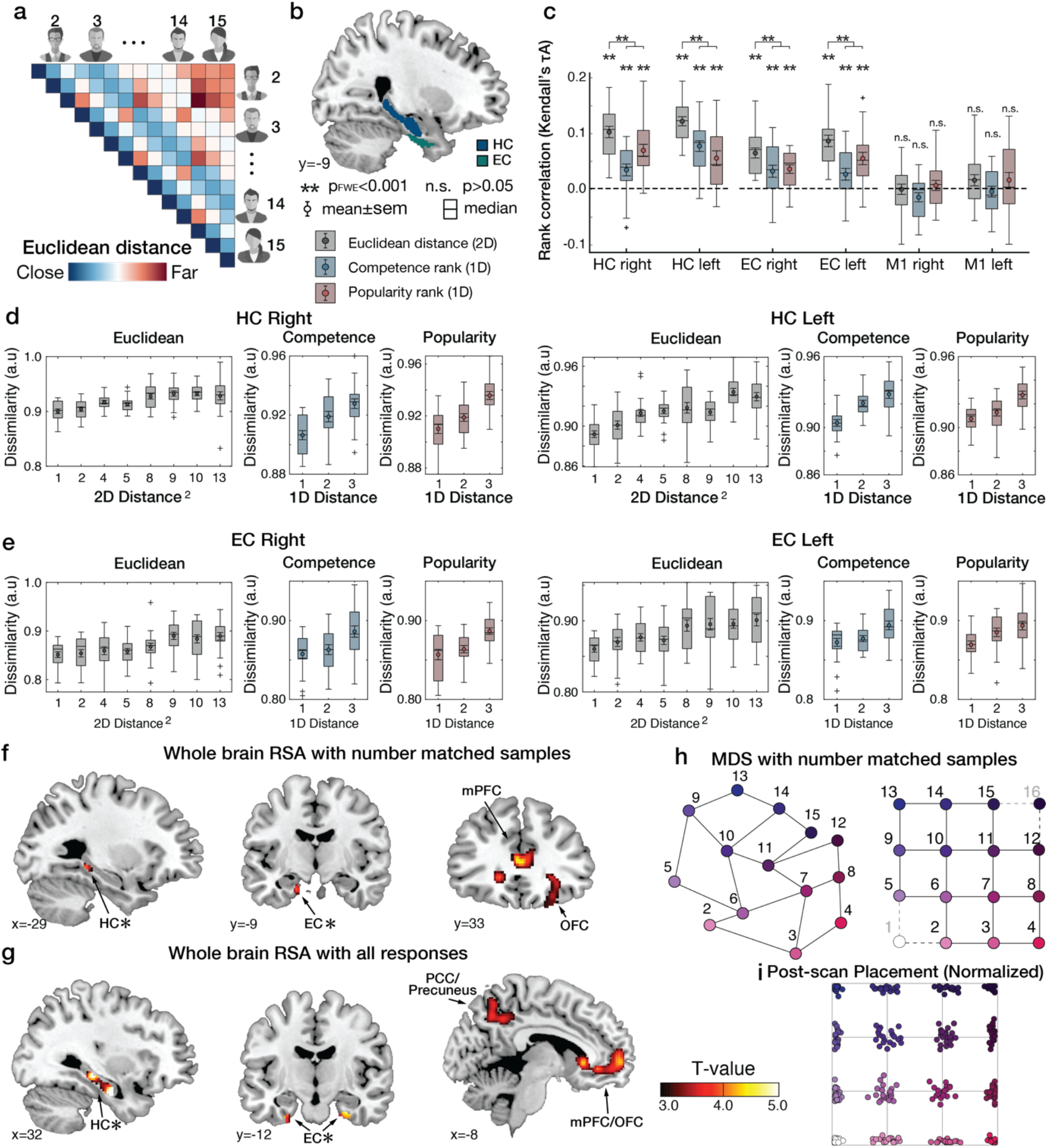
Building a 2-D representation of a social hierarchy. **a.** Model representational dissimilarity matrix (RDM) that models distances computed from the pairwise Euclidean distances between individuals on the true 2-D social hierarchy. **b**. We examine to what extent the model RDM explained the pattern dissimilarity between voxels in anatomically defined *a priori* regions of interest (ROIs) in bilateral hippocampus (HC) and entorhinal cortex (EC). The neural RDM was computed in the ROIs based on the pairwise Mahalanobis distance in the multi-voxel activity patterns evoked when face stimuli were presented. **c.** Representational similarity analysis (RSA) based equal sampling of 14 faces in the social hierarchy. The rank correlation (Kendall’s *τ_A_*) shows robust effects of Euclidean distance on the pattern dissimilarity (gray) estimated in the HC and EC compared to the permuted baseline (1000 iterations; dashed line), but not in a control region, the primary motor cortex (M1). The *τ_A_* of the one-dimensional rank distances are also shown for competence (blue) and popularity (red) dimensions. The *τ_A_* of the Euclidean distance was significantly greater than the one-dimensional rank distance in the competence and popularity (red) dimensions in each ROI (**Table S6**) (**, p_FWE_<0.001 after correction for the number of bilateral ROIs (n=4) with the Bonferroni-Holm method; n.s., p>0.05, uncorrected). **d and e.** The dissimilarity between activity patterns estimated in bilateral HC (**d**) and EC (**e**) increases in proportion to the true pairwise Euclidean distance between individuals in the 2-D abstract social space (gray). The dissimilarity between activity patterns increases not only with the 1-D rank distance in the competence dimension (blue) but also the 1-D rank distance in the popularity dimension (red). For display purposes, the dissimilarity level of each participant was normalized in the range of 0 to 1 and averaged to account for individual differences in scales (this normalization was not used for statistical inference). Box, lower and upper quartiles; line, median; whiskers, range of the data excluding outliers; +, the whiskers’ range of outliers. **f.** Whole-brain searchlight RSA based on equal sampling of 14 individuals (by down-sampling) at all events (F0, F1, and F2 presentations). **g.** Whole-brain searchlight RSA including all observations acquired from all events (F0, F1, and F2 presentations). The activity patterns in the HC, EC, mPFC/mOFC, and PCC/precuneus are explained by the model RDM for pairwise Euclidean distance (whole-brain TFCE correction, p_FWE_<0.05, except for HC and EC which was corrected using an anatomically defined *a priori* ROI [denoted by * next to HC and EC]). **h.** Visualization of the group representation of the social hierarchy in a 2-D space using multidimensional scaling (MDS) on the neural activity extracted from the HC ROIs while equating observations of each face. The MDS (left) captures the true 4×4 social hierarchy structure (right) better than random configurations (p<0.01 compared to 1000 random permutations; **Extended Data Figure 2**). **i.** Post-scan placement task. All participants successfully placed individual face stimuli according to their ranks in two dimensions in a 2-D social hierarchy space. For visualization purposes, the social hierarchy map of faces placed by each participant was rescaled according to their longest pairwise distance on the X and Y coordinates. Each dot indicates the position of a face placed by a single participant. The dots were colored according to their ranks in the true social hierarchy.

To examine the neural representation of the 2-D social hierarchy outside of our *a priori* ROIs, we performed a whole-brain searchlight RSA, both after down-sampling (**Fig.2f; Table S7a**) and based all observations (**Fig.2g; Table S7b)**. In both analyses, we found significant effects of Euclidean distances in the HC and EC (p_FWE_<0.05 threshold-free cluster enhancement (TFCE) (Smith and Nichols, 2009) correction in anatomically defined ROIs). In addition, we found significant effects in the medial prefrontal cortex (mPFC), orbitofrontal cortex, and posteromedial cortical areas (posterior cingulate cortex (PCC)/precuneus and retrosplenial cortex (RSC)), among other areas, in both analyses, and the inferior parietal lobule in the analysis incorporating all observations (p_FWE_<0.05 whole-brain TFCE correction) (**Fig.2g**).

Having found evidence that the 2-D social hierarchy is represented as a 2-D cognitive map in HC and EC, and interconnected cortical regions, we next searched for established fMRI markers of a grid-like code, but for inferred, rather than previously experienced, trajectories during decision making (Constantinescu, O’Reilly and Behrens, 2016). Specifically, we searched for neural evidence for hexadirectional modulation for inferred trajectories though the reconstructed cognitive map (**Fig.1 b,c**) that we hypothesized were used to compute the growth potential to guide decisions: 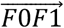 at the time of F1 presentation and 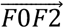 at the time of F2 presentation (**Fig.1e,f**). To test this hypothesis, we employed a two-step approach: first, we functionally identified ROIs showing hexagonal modulation for inferred trajectories independently of a participant’s EC grid orientation for future independent tests, and, second, we search across the whole brain for regions showing hexagonal modulation that is aligned to a participant’s EC grid orientation.

For the first step, we tested for regions where the BOLD signal was hexagonally modulated (i.e. any linear combination of sin(6θ) and cos(6θ) using a Z-transformed F-statistic (as in (Constantinescu, O’Reilly and Behrens, 2016)), where *θ* is the inferred trajectory angle between two faces in the social space). We found significant non-uniform clustering of putative grid orientations (*ϕ*) for all participants across voxels in the EC ROIs (p<0.01; mean z±SEM=50.98±4.14; **Fig.S7**). Note that this analysis was performed to functionally identify independent EC ROIs from which to compute the grid orientation angle *ϕ* for a given subject for future tests. We found evidence of hexagonal modulation, independently of grid orientation, in the EC (at Z>2.3, p<0.01) and additionally in mPFC, posterior cingulate cortex (PCC), and posterior parietal cortex (PPC) (p<0.05 whole-brain TFCE correction) (**Extended Data Figure 3a,b**; **Table S1**).

**Figure 3.**
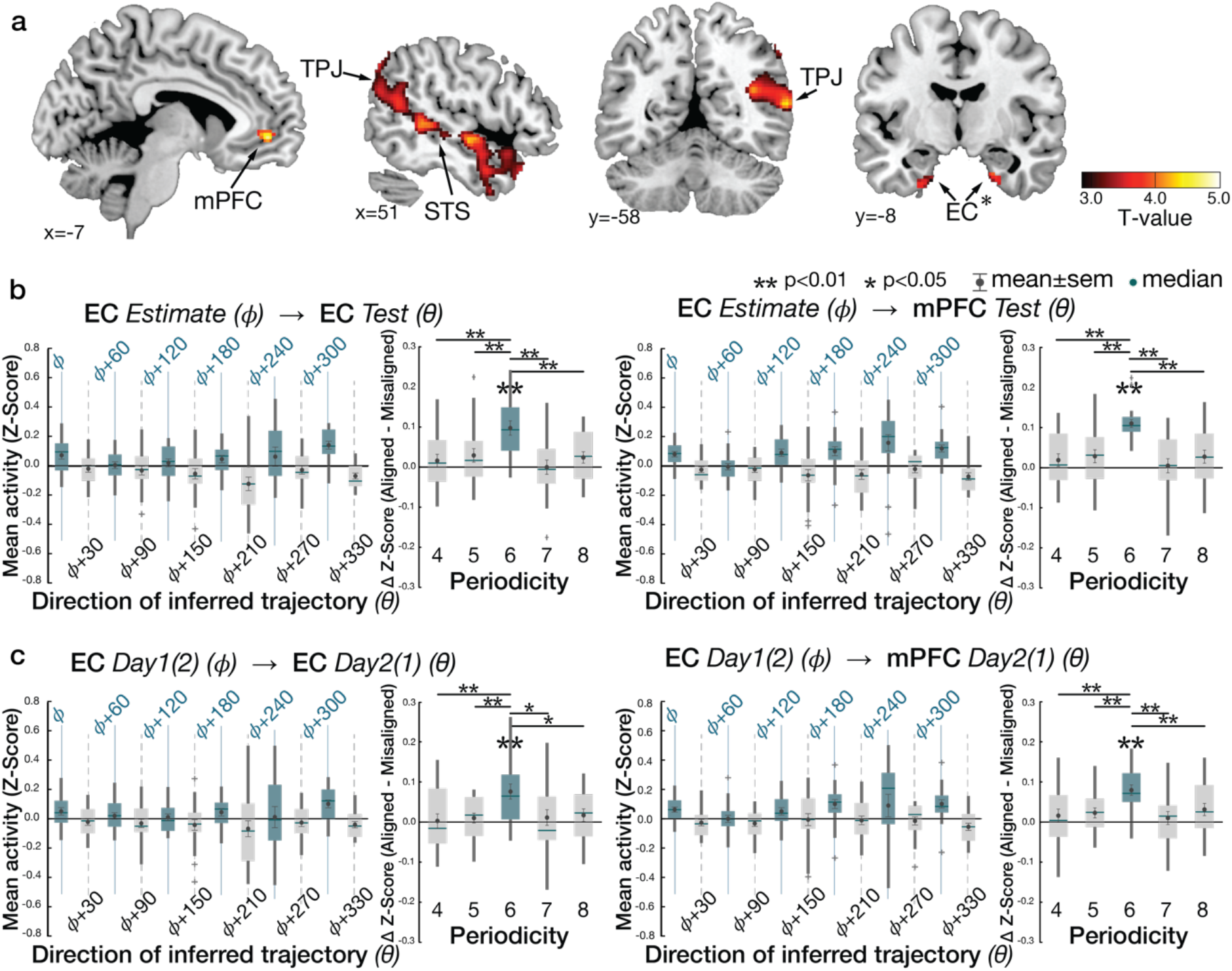
Hexagonal modulation for inferred trajectories. **a.** Whole-brain parametric analysis showing hexagonal grid-like representation of inferred trajectories in alignment with the mean EC grid orientation at the time of F1 and F2 presentations. Significant effects are shown in EC (peak MNI coordinates, [x,y,z]=[22,-10,-28], t=4.11), mPFC ([-6,48,-4]; t=4.72), STS ([50,-40,4], t=4.05 for right; [-60,-24,-6], t=4.29 for left), TPJ ([46,-58,20],t=3.67 for right; [-56,-68,24], t=5.71 for left) (all p_TFCE_<0.05, whole-brain cluster corrected using threshold-free cluster enhancement (TFCE) (Smith and Nichols, 2009)), and EC (peak MNI coordinates, [x,y,z]=[22,-10,-28], t=4.11) (p_FWE_<0.05 TFCE correction within a priori anatomical ROI). The maps are displayed at a cluster-corrected threshold p_TFCE_ < 0.05 over the whole brain for all brain regions, except for the EC, where we used a threshold of p_TFCE_ < 0.05 corrected within anatomically defined ROI (denoted by * next to EC), due to our strong a priori hypothesis of grid coding in EC (Hafting *et al*., 2005; Doeller, Barry and Burgess, 2010). **b.** Six-fold modulation signals in the EC (left panel) and mPFC (right panel) ROIs aligned to the grid orientation in EC. The grid orientation was estimated from separate fMRI sessions acquired from the same day. The mean (±SE) z-scored activity is plotted separately for aligned (teal) and misaligned (gray) trajectories categorized into 12 equal bins of 30° according to the direction of inferred trajectories. The mean activity difference between aligned and misaligned trajectories was larger than zero for six-fold (p<0.01) but not for the other control periodicities (four-, five-, seven- and eight-fold; all p>0.05), and the activity difference is greater for six-fold compared to the other periodicities (p<0.05). **c.** Cross-day consistency of the grid orientation in EC (See **Extended Data Figure 3d** for whole brain analysis). The activity in EC and mPFC shows hexadirectional modulations for the inferred trajectories in alignment with the grid orientation in EC estimated from separate sessions acquired from a different day more than a week apart. This effect is also specific to the six-fold (p<0.01) periodicity (all p>0.05), suggesting that the mean grid angle in EC is consistent between sessions more than a week apart. Box, lower and upper quartiles; line, median; whiskers, range of the data excluding outliers; + sign, the whiskers’ range of outliers. **, p<0.01, *, p<0.05.

We then tested for hexagonal modulation aligned to the EC grid orientation for each participant using an unbiased cross-validation (CV) procedure from fMRI sessions acquired in the same day. To obviate concerns of selection bias, we separated a ‘training dataset’ for estimating the putative grid angle *(ϕ)* from the EC ROI functionally identified by step 1 above from a ‘testing dataset’ on which the effects of hexagonal symmetry were tested (using the regressor *cos*(6[*θ_t_-ϕ*])) (see Methods and **Extended Data Figure 3c**). For this critical test, we observed hexagonal modulation for inferred trajectories aligned to the estimated grid orientation in bilateral EC and mPFC (*p*<0.05 TFCE-corrected; **Fig.3a,b**; **Table S2a**; see **Fig.S8a** for confirmatory analyses on independent anatomically defined EC (Amunts *et al*., 2005; Zilles and Amunts, 2010) and mPFC (Neubert *et al*., 2015) ROIs). Importantly, control analyses indicated that the modulation in EC and mPFC was specific to a six-fold periodicity, as significant modulation was not seen for four-, five-, seven-, or eightfold periodicities (all p>0.05). Furthermore, the effect at six-fold was significantly stronger than the control periodicities (all p<0.05) (**Fig.3b**).

Outside of these regions predicted from prior studies (Doeller, Barry and Burgess, 2010; Constantinescu, O’Reilly and Behrens, 2016), we also observed within-day hexagonal modulation aligned to the EC grid orientation in bilateral temporo-parietal junction (TPJ) areas and bilaterally along the superior temporal sulci (STS) (**Fig.3a**; see **Fig.S8b** for additional analyses of fusiform face area (FFA) ROIs). Notably, the TPJ, STS, and dorsal aspect of mPFC, in which we observed hexagonal modulation, are commonly recruited during tasks engaging theory of mind in social cognition (Hampton, Bossaerts and O’Doherty, 2008; Behrens, Hunt and Rushworth, 2009; Frith and Frith, 2012; Platt, Seyfarth and Cheney, 2016; Wittmann, Lockwood and Rushworth, 2018), including the modeling or simulation of different perspectives (Nicolle *et al*., 2012). Since participants were asked to take the perspective of the entrepreneur (F0) and decide with whom they should partner, perspective-taking on the cognitive map was a likely requirement to perform the partner selection task successfully. This requirement may therefore explain their recruitment, and alignment with the grid code, in our partner selection task.

Follow-up analyses tested plausible alternative decision-related vectors and subjectively defined cognitive maps. We did not find any evidence of hexadirectional grid-like coding of the alternative decision vector between F1 and F2 in the EC or mPFC (**Extended Data Figure 3e; Fig.S3d**). In addition, we tested for alternative cognitive map geometries. First, we confirmed that the activity patterns in these brain areas were not better captured by the inferred trajectories over a neurally-defined space, by using angles computed from the MDS of each participant’s HC (*θ*_MDS_) compared to those from the true social hierarchy structure (θ). Notably, it was not possible to clearly distinguish between distances defined on a space defined by strictly linear dimensions and one defined by certain nonlinear, monotonic dimensions, such as certain sigmoidal functions, in our task (**Fig.S6**).

To test whether grid alignment was consistent for the same behavioral task over time, we acquired another fMRI dataset on the same subjects (n=21) on the same task more than a week later (average duration = 8.95 ± 0.81 days). We found hexagonal modulation aligned to the EC grid calculated from the other (left-out) day in a similar set of brain regions including the EC and mPFC, and also in PCC (*p*<0.05 TFCE-corrected) (**Fig.3c**; **Extended data Figure 3**; **Table S2b**). Once again, the effect at the six-fold periodicity was significantly stronger than all other control periodicities (all p<0.05). This set of brain regions – EC, mPFC, and PCC – that showed a consistent grid-like code over time for novel inferences through a reconstructed social space during decision making align with previous findings for trajectories actually traveled through physical space to remembered locations in virtual reality (Doeller, Barry and Burgess, 2010).

To make optimal decisions in the partner selection task, participants need to compute and compare the growth potentials (GP) of each pair. The value of each decision option therefore corresponds to each pair’s GP. A whole-brain analysis of parametric effects of the GP at the time of face 1 and face 2 presentations revealed significant effects of the GP in bilateral TPJ, mPFC, and EC (p<0.05 TFCE-corrected; **Fig.4a**; **Table S3**), indicating that BOLD activity in these regions quantitatively reflected the GP of a pair under consideration. Importantly, the GP and hexagonal modulation (*cos*(6[*θ_t_-ϕ*])) regressors showed little correlation (mean r±SE =0.04±0.03), and both effects remained after including the other term as a regressor in the same GLM, indicating both terms explained independent variance in these regions (**Fig.S4a**). We further confirmed that the mPFC and right TPJ activity not only encoded the decision value (GP) but also showed a hexadirectional grid-like code for inferred trajectories for novel decisions (**Fig.S10**), even when controlling for effects of the Euclidean distances of the trajectories (p_FWE_<0.05 whole-brain TFCE correction, **Fig.S11**).

**Figure 4.**
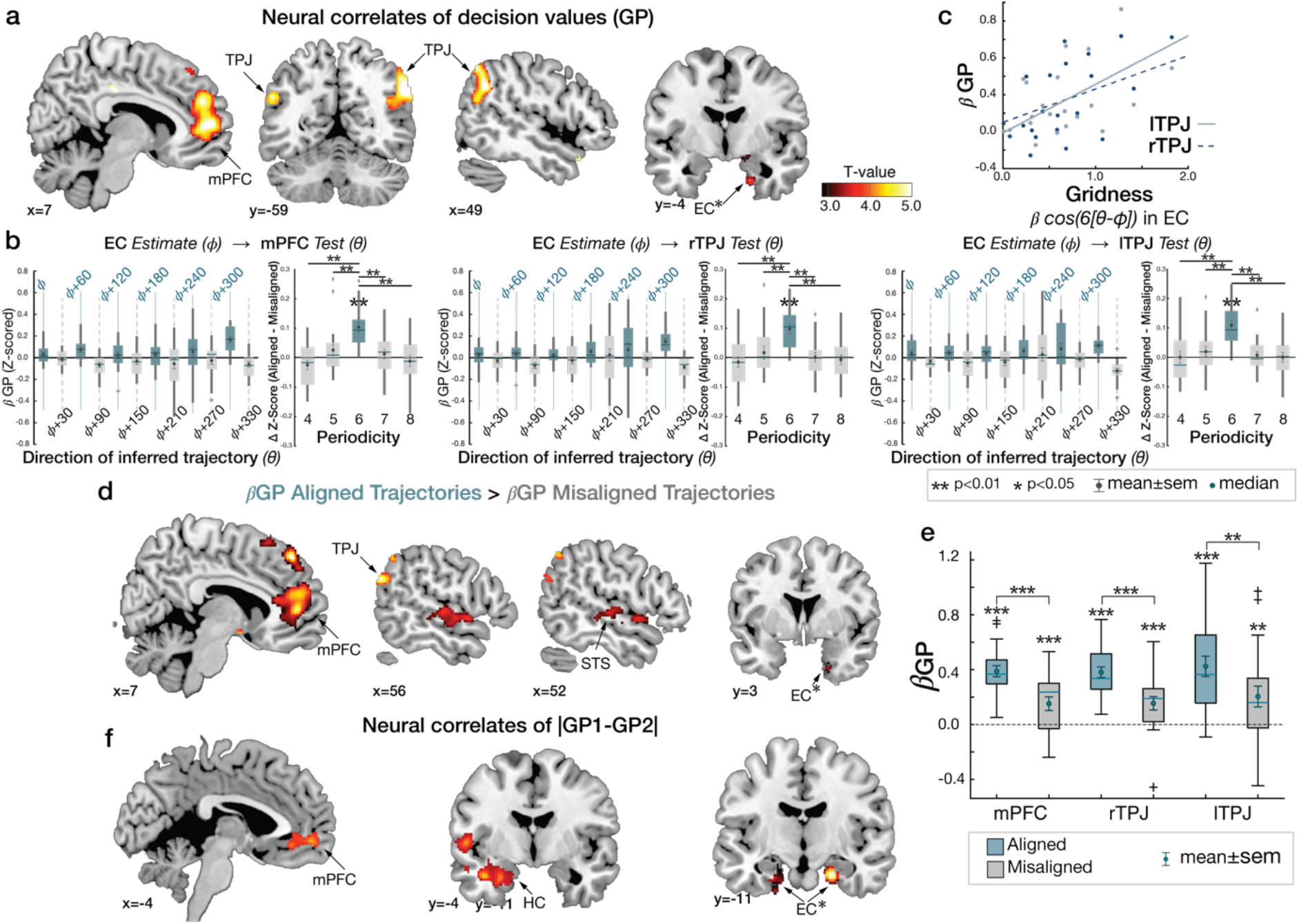
Growth potential and value comparison. **a.** Whole-brain map showing effects of the “growth potential” (GP) at the time of F1 and F2 presentation in a network of brain regions including the mPFC (peak MNI coordinates, [x,y,z]=[10,52,6], t=5.55) and bilateral TPJ ([54,-56,34], t=5.45) for right; [-54,-60,28], t=5.18) for left) (p_TFCE_<0.05 whole brain TFCE correction). The EC ([x,y,z]=[20,-4,-32], t=3.88) also showed GP effects. **b.** Mean (±SE) effects of GP (z-scored β) in mPFC and bilateral TPJ are modulated by the grid alignment of the inferred trajectories, *θ,* with six-fold periodicity aligned with the EC grid orientation, *ϕ.* Control analyses confirmed specificity for this six-fold periodicity (p<0.01) over control periodicities (effect of six-fold compared to other control periodicities, all p<0.05). **c.** Those participants with greater hexagonal modulation in EC show greater encoding of GP in independently defined bilateral TPJ ROIs (all p<0.05; **Extended Data Figure 5**). **d.** Whole-brain map contrasting the effects of GP (β GP) for trajectories aligned with the EC grid orientation, *ϕ,* compared to the misaligned trajectories (β GP aligned > β GP misaligned: p_TFCE_<0.05 whole brain TFCE correction). Contrasts are shown in mPFC, EC, STS, and TPJ. **e.** Mean (±SE) GP effects in anatomically defined mPFC, and bilateral TPJ ROIs (β GP). In all three areas, effects were greater for aligned compared to misaligned trajectories (all p<0.01), although the GP effects of both trajectories were significant (all p<0.01). **f.** Neural correlates of the relative decision value, IGP1-GP2I, during decision making, in vmPFC ([-8,54,-6]; t=5.07), HC ([28,-4,-26]; t=4.53) (p_TFCE_<0.05), and bilateral EC ([22,-12,-26], t=5.10 for right; [-26,-16,-32], t=4.00 for left; p_TFCE_<0.05 TFCE corrected in *a priori* ROIs). ***, p<0.005; **, p<0.01; *, p<0.05. All the maps are displayed at a cluster-corrected threshold p_TFCE_ < 0.05 over the whole brain for all brain regions, except for the EC, where we used a threshold of p_TFCE_ < 0.05 corrected within our anatomically defined ROI (denoted by * next to EC).

We hypothesized that the grid code for inferred vectors may be used to estimate the relative rank distances between F0 and F1(F2) in each of the two dimensions, which may further underlie the computation of the growth potential of each pair based on a cognitive map in our task (**Fig.S3d**). If true, we predicted that the GP effects in TPJ and mPFC regions would be modulated by trajectories that were aligned compared to misaligned to the EC grid orientation. Examining the TPJ and mPFC ROIs functionally defined using the statistically independent GP regressor (at p<0.001 uncorrected) showed greater GP effects for trajectories that were aligned compared to misaligned *with the EC grid orientation* in these regions (**Fig.4b**). Notably, while there was a significant GP effect in TPJ and mPFC for both trajectories that were aligned and misaligned to the grid orientation, the effect was modulated such that it was significantly stronger for aligned trajectories (**Fig.4d,e**; **Table S3c**), consistent with a dependence of GP coding on EC grid coding. Control analyses confirmed specificity for this 6-fold periodicity in TPJ and mPFC over control periodicities (all p<0.05) (**Fig.4b**). This dependence of GP coding on EC grid alignment may be hypothesized to influence behavior. To test this possibility, we examined behavioral measures for inferred trajectories that were aligned or misaligned to the EC grid orientation, but did not find a significant effect in RT (t_20_>1.5, p>0.05; **Fig.S4**) or accuracy (though there was limited behavioral variability in our subject sample).

We performed several confirmatory analyses and robustness checks, both for the GP effects aligned to the EC grid orientation and the grid-like coding reported previously. First, we confirmed that these effects were not driven by particular trajectories with specific angles sampled in sub-regions of the 2-D space (**Table S4**). Second, neither effects depended on the position of F0 in the pair, which was preferentially sampled to discourage the use of choice heuristics (p>0.05; **Fig.S9a**). Last, we confirmed these effects could not be explained by a memory of inferences made on previous trials in the absence of choice feedback but were utilized for novel inferences performed for the very first time (i.e. novel pairs of F0 and F1 [F2]). Specifically, we confirmed the hexadirectional effects in EC and mPFC were robust when testing only for the pairs that were presented for the first time during the first day of the partner selection task (n=88 pairs presented for the first time on Day 1 for 17 participants [83 pairs were novel for 4 participants]; 11 [16] pairs had been presented previously during practice, though they were compared against a different pair during decisions and in the absence of any choice feedback; **Extended Data Figure 4**; p<0.05). This analysis indicates that the grid-like code and GP dependence on EC grid coding are both present for novel inferred trajectories composed on the fly.

We further reasoned that if the grid code for inferred trajectories is related to the process of constructing the values of decision options (here, the pair’s GP), then those subjects with greater hexagonal modulation in EC would also be predicted to show greater neural GP effects. Consistent with this prediction, we found a positive correlation between the EC grid modulation effect and the GP effect in TPJ (**Fig.4c**; r=0.58, p=0.006 for left TPJ and r=0.44, p=0.05 for right TPJ), but not in mPFC (p>0.05), across participants, after controlling for choice accuracy. In post-hoc analyses, when we instead used the mPFC grid modulation effect, we found a marginal positive correlation with the GP effect in mPFC (r=0.42, p=0.057) and right TPJ (r=0.42, p=0.06) (**Extended Data Figure 5**). While there was limited inter-subject variability in behavioral measures, we additionally tested for behavioral correlates of grid effects. We did not find a significant effect of the grid-like coding in EC on trial-by-trial RT (**Fig.S4**; **Table S5**), nor a significant correlation between the extent of a subject’s grid modulation and the degree to which the difference in GPs explained RT across participants (r=-0.19 for EC, r=0.04 for mPFC; p>0.1).

At the time the second face (F2) is presented, subjects have all the information needed to make a decision based on the growth potential of F0 collaborating with one partner compared to the other (a comparison of the two potential collaborations’ GPs). Considerable evidence has supported a role for the ventromedial prefrontal cortex (vmPFC) in representing and comparing options’ reward values for goal-directed decisions, a signal that also reflects decision confidence (Daw *et al.*, 2006; Behrens *et al.*, 2008; Boorman *et al.*, 2009; FitzGerald, Seymour and Dolan, 2009; Lim, O’Doherty and Rangel, 2011; Noonan, Mars and Rushworth, 2011; Boorman, Behrens and Rushworth, 2011; Hunt *et al*., 2012; Kolling *et al*., 2012; Clithero and Rangel, 2013; De Martino *et al*., 2013; Lebreton *et al*., 2015; Yamada *et al*., 2018; Park *et al*., 2019). Searching for the equivalent value comparison term in our task – a comparison between the two options’ values, or |GP1 – GP2| – replicated this effect in vmPFC, and also revealed a significant effect in the EC extending into the HC (p<0.05 TFCE-corrected) (**Fig.4f**; **Table S3b**). Furthermore, participants who showed a greater hexagonal modulation in EC tended to show greater effects of the GP difference in vmPFC (r=0.43, p=0.05; **Extended Data Figure 5b**). These effects in EC/HC are consistent with the interpretation that the comparison between options’ values during decision making additionally involves the interconnected HC–EC system (Witter *et al*., 2000) when it is based on a cognitive map (Bakkour et al., 2019).

## Discussion

We have shown that abstract and discrete relational structures, never seen but inferred from discrete samples through a series of binary comparisons on each dimension made on different days, are reconstructed into a 2-D cognitive map in the HC-EC system. Specifically, we found that the closer faces were ranked in the true 2-D social hierarchy, the increasingly more similar their neural activity patterns were in the HC and EC, and also the interconnected mPFC, OFC, and PCC/RSC (Barbas and Blatt, 1995; Ferreira-Fernandes *et al*., 2019). This finding suggests that each entity is located according to its feature values along relevant dimensions in a 2-D relational space (Eichenbaum and Cohen, 2014; Behrens *et al*., 2018). Furthermore, we found evidence for hexagonal modulation for inferred direct trajectories over the abstract social space during discrete decision making, further supporting the interpretation that people infer over a 2-D cognitive map of abstract relationships. Our findings thus suggest that having an explicit representation of structural relationships is beneficial for generalizing experiences that can be used to guide inferences during novel decision making.

We found evidence for a grid-like code in EC, mPFC, PCC, TPJ, and STS, for inferred direct trajectories through the reconstructed 2-D social space during decision making. This finding provides novel evidence that humans can use direct inferred trajectories in non-spatial decision making, a defining feature of a cognitive map of space (O’Keefe and Nadel, 1978; Bennett, 1996). Recent studies suggest that our brains have a representation of the structure of social relationships during social interactions (Tavares *et al*., 2015; Parkinson, Kleinbaum and Wheatley, 2017; Stolier, Hehman and Freeman, 2018; Tamir and Thornton, 2018). One study showed that the HC activity correlates with the vector angle defined by power and affiliation dimensions (Tavares *et al*., 2015), suggesting that the HC and other regions may use a polar coordinate system. If the brain uses a polar coordinate system to locate individuals along abstract social dimensions during social interactions, one plausible hypothesis in the current task is that the brain would compute the vector angles to estimate the rank distance and relative position in the social hierarchy (**Fig.S3d**). Our results indeed show that the brain encodes the vector angle to relate individuals in a multidimensional space which allowed participants to integrate separately learned relationships in each dimension to make accurate inferences for novel decisions that they have not made before. Taken together with recent findings in non-social tasks (Doeller, Barry and Burgess, 2010; Jacobs *et al*., 2013; Constantinescu, O’Reilly and Behrens, 2016; Nau *et al*., 2018; Bao *et al*., 2019), these findings suggest that grid-like codes are present in a markedly common set of core cortical brain regions including the EC and interconnected mPFC and PCC (Witter *et al.*, 2000; Preston and Eichenbaum, 2013; Eichenbaum, 2017) for both spatial and non-spatial relational tasks. Furthermore, the grid-like code’s alignment between EC and mPFC was consistent when performing the same task over a week later, consistent with a previous study (Constantinescu, O’Reilly and Behrens, 2016), indicating a relatively stable representation over time, even in the absence of practice. Here, we extend these previous reports by showing for the first time that first, a grid-like code can be used for discretely sampled abstract spaces and second, to compose novel inferences during non-spatial decision making.

On the other hand, our findings also suggest that additional brain regions – TPJ, dorsal mPFC, and STS (and also to some extent FFA [see **Fig.S8b**]) – may come online in shared alignment with the EC grid code, depending on the specific processes required by a particular task. Notably, these areas are thought to play an important role in theory of mind (Hampton, Bossaerts and O’Doherty, 2008; Behrens, Hunt and Rushworth, 2009; Frith and Frith, 2012; Platt, Seyfarth and Cheney, 2016; Wittmann, Lockwood and Rushworth, 2018), which frequently involves simulating another person’s mental state (Nicolle *et al*., 2012). Since participants were explicitly asked to select a partner on behalf of another entrepreneur, a manipulation that served to anchor trajectories to different locations in the social space, perspective-taking on the cognitive map was engaged in our task. Whether the grid-like modulation of the BOLD signal in these regions reflects grid cells locally outside of EC (Jacobs *et al*., 2013; Long and Zhang, 2021), or synchronized input between EC and other interconnected cortical areas (Witter *et al*., 2000; Preston and Eichenbaum, 2013; Eichenbaum, 2017), remains an intriguing open question for future studies.

Finally, our study provides novel evidence linking a grid-like code to the construction of decision values in mPFC and TPJ during decision making. Specifically, we found that the growth potential of a pair, equivalent to the decision option’s value in our task, was encoded in the mPFC and TPJ, after controlling for the hexagonal modulation effect and Euclidian distance between individuals. This finding is consistent with past studies showing these areas encode another person’s decision value when predicting their decision (Nicolle *et al*., 2012; Piva *et al*., 2019). Importantly, this effect was modulated by the inferred trajectory’s alignment to the EC grid orientation such that the value was encoded more strongly for aligned than misaligned trajectories. Notably, hexagonal modulation of theta band local field data from pre-surgical human recordings has been identified for traveled trajectories in virtual reality (Maidenbaum *et al*., 2018; Staudigl *et al*., 2018). Moreover, recent evidence indicates a critical role for theta-phase synchrony between the hippocampal formation and orbitofrontal cortex that drives stimulus value coding in orbitofrontal cortex during value-based decision making (Knudsen and Wallis, 2020). Thus, one interpretation of the dependence of value coding in mPFC and TPJ on EC grid alignment in our study is that the grid code is theta-phase locked to the coding of the growth potential in mPFC and TPJ, thus providing an information channel between EC and these regions. Moreover, we found that those individuals who showed a stronger hexagonal modulation in EC also showed stronger GP effects in TPJ, even after controlling for choice accuracy. Collectively, these findings suggest that the grid code plays a central role in value construction when it is based on inferences over a cognitive map.

We reasoned that in order to have any chance to identify the grid-like coding we hypothesized, we needed to ensure it would be measurable. For this reason, the behavioral training protocols were designed to encourage participants to treat ranks as equidistant intervals and to use the equal rank intervals per dimension, thereby facilitating measurement of the hypothesized direct vectors. We acknowledge that we cannot rule out the possibility that the requirement of high accuracy in behavioral training may have excluded some participants who constructed different space geometries. However, during the partner selection task, participants were asked to make decisions that they had not made before. Decision values experienced in behavioral training thus cannot be generalized to those for the partner selection decisions during fMRI. Notably, during fMRI the growth potential of a pair cannot be consistently predicted by the rank of a single individual, but only by the combination of multidimensional information between two individuals (**Fig.S2**). Thus, while the GP cannot be mapped on the 4×4 social hierarchy structure, the partner selection decisions did not appear to impact the representation of the structure of the social hierarchy, since this was consistent across sessions within a day and even between days over a week apart. Moreover, it would be counterproductive to maintain a separate representation of the GP in a 16×16 structure, as well as the 4×4 social hierarchy structure. Rather, using the 4×4 social hierarchy structure theoretically enables subjects to flexibility compute the GP, even of new pairs, on the fly. This conclusion is supported by our RSA findings of Euclidian distance, grid-like coding for direct trajectories, and grid orientation consistency across sessions acquired more than a week apart.

Importantly, subjects do not need to represent the social hierarchy in a 2-D space to solve our tasks. Both for the behavioral training task, and for the novel decisions of the partner selection task, there is no *a priori* reason that these tasks need to be solved with a 2-D map-like representation. Alternatively, subjects could use two separate 1-D number lines (Piazza *et al*., 2004) or update the rank values with neither 1-D nor 2-D maps (von Fersen *et al*., 1991; Doucet, Godsill and Andrieu, 2000; Frank, Rudy and O’Reilly, 2003). If participants’ brains used either strategy above, they still could theoretically pass our performance criteria but should show no evidence of a 2-D neural representation and grid-like coding. Our results show that human participants integrate two separately learned 1-D social hierarchies into a 2-D representation even when there is no explicit task demand to do so and use grid-like codes to make novel decisions whose outcome they had never learned about before.

Taken together, our findings suggest the cognitive map in the HC-EC system, and interconnected “default-mode” regions, provides a systematic framework which the brain can utilize for vector navigation through spaces defined by abstract relationships, allowing discrete entities or items to be related to each other in new situations, and thus generalization and the discovery of new solutions for choice. We suggest the alignment of the “default mode” network, and potentially other regions, with the entorhinal grid system may reflect a general mechanism for knowledge-guided inference during learning and decision making.

## Methods

### Participants

A total of 25 participants (13 females, age range: 18–25, normal or corrected to normal vision) were recruited for this study via the University of California, Davis online recruitment system. Using six motion parameters to assess motion in each participant, we excluded the data of four participants who had movement in the x, y, and/or z directions greater than the voxel size (3 mm) relative to the middle time point across each scanning block and who had total displacement of greater than 3 mm from the beginning to the end of the scanning session. In total, 21 participants’ data were included in the analysis (10 female, mean age: 21.14±0.52, standard error mean (±SE)). The study was approved by the local ethics committee, all relevant ethical regulations were followed, and participants gave written consent before the experiment.

Participants received course credit as compensation for participating in day 1 and day 2 training. Participants received monetary compensation for their participation from day 3 training. In addition to the compensation, participants have been told that extra monetary rewards will be earned if they pass the training (see the criteria below in behavioral training) and these extra rewards would be paid if they completed the fMRI task.

### Social hierarchies

During behavioral training, participants were asked to learn social hierarchies between 16 entrepreneurs presented as face stimuli which were constructed in two independent ability dimensions – popularity and competence. Each dimension had four levels of rank. Four individuals were allocated at the same rank at each level. The rank of an individual in one dimension was not related to his/her rank in the other dimension and each had a unique combination of popularity and competence ranks. This allowed us to create a 4×4 social hierarchy structure comprising two social hierarchy dimensions (**Fig.1b**).

It is important to note that the true structure of social hierarchy (not only the twodimensional (2-D) social hierarchy but also each one-dimensional (1-D) social hierarchy) was never shown to participants. Moreover, participants were never given any information implying the structure of the social hierarchies, including the total number of ranks in each dimension and the number of individuals allocated to the same rank.

### Stimuli

The stimuli consisted of 16 grayscale photographic images of faces (Strohminger *et al*., 2016). All of the images were adjusted to have the same mean grayscale value. To ensure that visual features of a face (including gender, race, and age) were not associated with the rank of the entrepreneur, we prepared eight sets of 16 face stimuli selected from a pool of 42 faces. Each of the stimuli sets comprised 16 faces. A set of stimuli among eight was randomly assigned across participants. Stimuli were presented to participants through a mirror mounted on the head coil. The face stimuli presented in this paper are license-free images for display purposes.

### Behavioral training

The 16 face stimuli were introduced as entrepreneurs. Each day of the behavioral training comprised ‘Learn’ phases and ‘Test’ phases. During a Learn phase of behavioral training, participants learned the relative rank between entrepreneurs (face stimuli) in one of the two independent dimensions through feedback-based binary comparisons. Specifically, participants were asked to learn which individuals were more capable of attracting crowdfunds (labelled popularity) and which individuals had higher technical proficiency (labelled competence). Participants only learned about individuals who differed by one rank level and on one dimension at a time. Participants learned the relative status of all possible one rank difference pairs with feedback in random order. During the first two days of training, participants completed two learning-blocks. Each learning-block is followed by a test-block. During a learning-block, 48 possible pairs of individuals whose ranks have one rank difference in the given dimension were presented twice (96 trials). During test-blocks, participants make inferences on the relative status of a pair of individuals who differed by one or more levels on one dimension while no feedback is given. During a test block, all possible 96 pairs were presented. The pairs presented for the learning and test blocks are depicted in **Fig.S1**. It is important to note that the relative ranks in each dimension were learned on a separate day (e.g. popularity on day 1 and competence on day 2 or vice versa). The day 2 training occurred 2 days after day 1 training. During day 2 training, participants learned the hierarchies on the other, different dimension (**Fig. 1d and Fig. S1**).

Participants were never shown the 1 -D or 2-D structure of the social hierarchy, never asked to combine the two social hierarchy dimensions during training, and never asked to solve the task spatially. However, participants could theoretically reconstruct the social hierarchy through transitive inferences not only on two one-dimensional structures but also on a two-dimensional structure by integrating the two dimensions learned on different days (as shown in the right panels in **Fig.S1**). During training, participants were not been informed about the fMRI task (Partner selection task). Therefore, participants were not able to anticipate having to build a representation of the combined social hierarchy for future use for novel inferences. During behavioral training, each of the 16 face stimuli had been exposed to participants the same number of times.

#### Training day 1

Participants learned the hierarchy of 16 individuals in either the competence or popularity dimension. On each trial during the Learn phase, two face stimuli of entrepreneurs were presented with a color cue (red or blue square) which indicated the task-relevant dimension. Participants were asked to choose the one who is superior to the other (the higher rank individual) in the given dimension (**Fig.S1e**). They received feedback at the end of every trial. Before the training, participants were informed that two entrepreneurs presented in a given trial during the Learn phase differed by only one rank on the given dimension (**Fig.S1a**). The first learned hierarchy and the color associated with each dimension were counterbalanced across participants.

During the Test phase, which was followed by the Learn phase, participants’ knowledge about social hierarchy in the learned dimension was tested. On each trial during the Test phase, similar to the Learn phase, participants chose one of two face stimuli who is superior to the other in the given social hierarchy. Participants did not receive any feedback for their choices during the Test phase. Before the training, participants were informed that two entrepreneurs presented in the Test phase would have one or more rank differences (dotted lines in **Fig.S1e**). To make a correct decision, participants needed to use transitive inferences to infer the relative status of two individuals who have never been compared during learning (**Fig.S1a**).

#### Training day 2

Using the same method through which the participants learned the social hierarchy in one dimension during day 1 training, participants learned the hierarchy of the same 16 entrepreneurs in the other unlearned dimension (**Fig.S1e**). For example, participants who learned about the hierarchy in competence dimension on day 1 learned about the hierarchy in popularity dimension on day 2 (see **Fig.S1b**).

At the end of day 2 training, participants were asked to perform an additional test which we called the “Test 2” phase. During the Test 2 phase, we tested participants’ ability to make flexible inferences while the task-relevant dimension was interleaved across trials. As with the previous Test phase, on each trial of the Test 2 phase, participants were asked to choose the superior face between two entrepreneurs who have one or more rank differences in the given dimension (**Fig.S1f**), and no feedback was given for their decisions. However, in the Test 2 phase, the relevant dimension for decision making was randomized across trials. Only the participants who were able to reach more than 80% accuracy in the Test 2 phase were invited to continue to the day 3 training (**Fig.S1b**).

#### Day 3 and after training

##### A. Flexible inferences in intermixed behavioral contexts (Test 2)

The day 3 training started with another Test 2 phase in which participants’ knowledge about social hierarchy was tested with the task-relevant dimension randomized across trials. As before, participants were asked to infer the relative rank between two individuals with one or more rank differences without feedback. From the day 3 training, on each trial of Test 2 phase, stimuli were presented sequentially in the order of the conditional cue (red or blue square) indicating the task-relevant dimension, a face stimulus (F1), and another face stimulus (F2) (**Fig.S1d**). A fixation cross was shown for the inter-stimulus-intervals and inter-trial intervals. During F2 presentation (2s), participants were asked to indicate the higher rank individual between F1 and F2 in the given dimension. During the Test 2 phase, participants were presented all possible pairs of entrepreneurs (**Fig.S1f**) twice by altering the position of F1 and F2 on interleaved dimensions, and they did not get feedback to their decisions. Only the participants who could make accurate inferences at 95% or more in both dimensions finished the training and were given the introduction about the partner selection task.

##### B. Conditional Learn phase

There were additional Learn phases for the participants whose accuracy was below 95% in one of two dimensions for the Test 2 phase on day 3 training. This additional Lean phase comprised the two Learn phases given on the day 1 and day 2 training. That is, participants were asked to choose the superior face between two entrepreneurs who differed by one rank on the given dimension while getting feedback to their decisions. Participants learned the hierarchy in one dimension at a time during each of two blocks in the Learn phase (**Fig.S1d**).

After the Learn phase, participants were asked to repeat the Test 2 phase of day 3 training. It could potentially take multiple days of training until the threshold was reached (>95% in both dimensions) **(Fig.S1d**). This high-performance threshold was selected because we reasoned it would be necessary to accurately measure hypothesized grid angles for planned fMRI analyses.

#### Behavioral training of the “partner selection” task

Participants who successfully learned the social hierarchy in both dimensions independently subsequently learned the partner selection task for the first time. In each trial of partner selection task, participants were asked to choose the better business partner for a given entrepreneur between two candidate partners while considering their relative hierarchy in both social hierarchy dimensions. Therefore, each decision during the fMRI task required novel inferences to be made by combining the two social hierarchy dimensions learned separately. See the following section of “Partner selection task” for details.

The behavioral training of the partner selection task had two purposes. First, it was to ensure participants understood the novel decision-making task in which they had never been exposed. Second, it was to ensure participants did not use alternative heuristics for decisionmaking by providing test trials where the potential heuristics failed (See the Partner selection task for details). To do this, after the instructions were given, participants were asked to perform 24 trials of the partner selection task without time constraints. To prevent learning, we did not give any feedback to their decisions during the training of the partner selection task. They were only informed whether they made all correct selections or not at the end of a block which comprised of 24 trials. We generated three sets of 24 trials for the purpose of behavioral training. Those trials were not included in the fMRI experiments to make sure participants made novel decisions for the fMRI experiment. That is, during fMRI, participants did not compare the same two pairs about which they had made decisions during behavioral training on the partner selection task. The pairs that were presented during the behavioral training on the partner selection task were paired with different pairs if they were also shown during fMRI. Therefore, the winning pair in a trial of behavioral training will often be a losing pair during fMRI or vice versa (**Fig.S3b**). Therefore, remembering the winning pair from previous trials of the partner selection cannot be an effective solution for the task and could not produce the high performance level we observed in our sample. Participants who failed to make all correct inferences in 24 trials were further asked to read again the instructions and to explains the instruction to the experimenter to ensure that they understood the task. Participants were asked to complete the same set of 24 trials in a randomized order until they made all correct decisions. After reaching 100%accuracy, participants were asked to complete the same set of 24 trials in a randomized order with time constraints (5 s), like in the fMRI experiment. To ensure that participants were not simply relying on remembered correct answers of the selected set were able to generalize their knowledge to make novel inferences, we confirmed that all participants could also make correct decisions for a novel set of 24 trials of the partner selection task. Finally, using this procedure, we recruited participants who were reach an accuracy of 95% or more in both dimensions for more than two trial sets (48 trials total) to participate in the fMRI experiment.

#### Participant retention

107 participants were recruited and started the initial training. 15 participants participated only in the day 1 training. Among 92 participants who finished the day 2 training, 39 participants passed the threshold of behavioral accuracy (>80% in the Test2 phase). The retention rate was 42.39%. Among 39, 29 participants agreed to continue training after day 3 training. The relatively high dropout rate in part reflects difficulties retaining subjects for effortful and time-intensive multi-day studies, variable performance due in part to using course credit as incentives (many students had achieved full credits for their courses before the end of Day 2 training, and 10 participants who passed the threshold of behavioral accuracy decided not to continue the training), and true variability of task performance in our limited initial training timeframe. 27 of 29 participants reached more than 95% of behavioral accuracy during the Test2 phase during day 3 and after training. The bar graph in **Fig.S1 g** shows the number of additional sessions that each of 29 subjects participated in. 2 among 29 participants stopped the training before reaching 95% accuracy, and 2 who reached >95% stopped the experiment before scanning (marked with * in **Fig.S1g**). We included a high-performance threshold because we needed to ensure accurate representations of cognitive maps, should they exist, to be able to measure the effects of grid-like coding reliably. We scanned 25 subjects who agree to participate in fMRI part of the experiment.

### Partner selection task (fMRI experiments)

Participants were given the following instructions. The goal of the partner selection task was to find a better business partner for a given individual (F0) between two candidate partners (F1 and F2). F0, F1, and F2 were indicated as face stimuli which were selected from the 16 entrepreneurs that had been presented to participants during behavioral training. To find a better business partner for F0, participants need to compute the ‘growth potential’ (GP) of each potential collaboration. The GP indicates the level of benefits that F0 could expect from the potential collaboration with each of two partners, F1 and F2. Participants were subsequently asked to choose between F1 and F2 according to the GP of each of two pairs – GP1: GP for F0-F1 pair; GP2: GP for F0-F2 pair.

To estimate GP accurately, first, participants were asked to compute the ‘rank of the pair’ in each dimension which was determined by the rank of the entrepreneur who had a higher rank (where the lowest rank was one). For example, if F0 was at a higher rank than F1 in the competence dimension while F1 was at a higher rank than F0 in the popularity dimension, the rank of the F0-F1 pair would be the rank of F0 in competence dimension and rank of F1 in popularity dimension. Second, participants were asked to weight the ‘rank of the pair’ in both dimensions equally to compute the GP. For example, when the ranks of F0 and F1 in the competence and popularity dimensions are [*C*_*F*0_,*P*_*F*0_] and [*C*_*F*1_,*P*_*F*1_], respectively, the ‘rank of the pair’ would be *max* (*C*_*F*0_, *P*_*F*1_) in the competent dimension and *max*(*P*_*F*0_, *P*_*F*1_) in the popularity dimension. And the GP1 would be determined by multiplication of two ‘ranks of the pair’. That is, *max* (*C*_*F*0_, *C*_*F*1_) × *max* (*P*_*F*0_, *P*_*F*1_). Last, participants were asked to make a binary choice between F1 and F2 to find a better partner for F0 by selecting the pair whose GP was greater than the other pair: choose F1 if GP1 *>* GP2; choose F2 if GP2 *>* GP1. During the partner selection task, participants did not receive any feedback for their decisions. Participants were notified that they would receive extra rewards in proportion to their overall performance, if their overall accuracy was greater than 90% at the end of the experiments (after the second-day scan).

As **Fig.1e** illustrates, in each trial of the partner selection task, participants were presented with two pairs of face stimuli sequentially in the order of F0, F1, F0, and F2. The stimuli, F0 were presented for 2.5 s and the stimuli, F1 and F2 were presented for 5s. In each trial, the inter-stimulus fixation was presented between F0 and F1 and between F0 and F2 for the same duration (a white cross, 2~5 s jittered). F0 was presented before each partner to anchor hypothesized direct vectors to the partner. The F1 stimuli were followed by a 3 s an inter-stimuli fixation (a purple cross). Participants were asked to make a binary decision to indicate a better business partner for F0 between F1 and F2 during the F2 presentation by pressing a button. The inter-trial fixation was presented at the end of each trial (green cross, 1~4 s jittered). During the fMRI experiment, the color of the fixation cross on the middle of the screen informed participants the progress of the trial. No feedback was given to the participants. Trials that participants did not respond to within the allotted time were shown later after a random number of trials to make sure that all subjects made responses to every trial.

### Novel inferences for the partner selection task

During the partner selection task, participants were asked to make decisions that they had not made before. Though participants were trained extensively to learn the rank differences in each one-dimensional social hierarchy, participants still needed to make inferences of the decision value for novel decisions in the partner selection task. During behavioral training, participants only compared the ranks of two individuals in one social hierarchy dimension at a time. This is also true for the Test2 blocks of the behavioral training in which the two dimensions were intermixed across trials. Importantly, there is no *a priori* reason that participants need to construct a combined 2-D representation of the social hierarchy during behavioral training. Instead, participants could still reach a high accuracy level by constructing two separate 1-D representations for the competence and popularity dimensions that were learned with feedback on different days (**Fig.S1c**) or by updating separate competence and popularity values under certain assumptions (Kumaran *et al.*, 2016). However, during fMRI, participants need to make inferences to compute a pair’s ‘growth potential (GP)’ and make a novel comparison against another pair to make a correct decision in the partner selection task. Therefore, though the same pair of individuals may be presented as in training, participants need to compute different decision values for the partner selection task compared to the behavioral training task. If participants’ brains use two 1-D representations, the brains should not encode the vector angles of inferred trajectories, which are defined on a 2-D space, while making decisions in the partner selection task. **Fig.S3a** shows a representative example in which the decision value for behavioral training of the same pairs cannot be generalized for the partner selection task.

Notably, here we use the term ‘inference’ in a manner consistent with its usage in other studies on transitive inference (Kumaran, Melo and Duzel, 2012; Kumaran *et al*., 2016) and modelbased inference using a pre-sensory conditioning procedure (Jones and Mishkin, 1972; Barron *et al*., 2020; Wang, Schoenbaum and Kahnt, 2020). In those studies, participants are trained on predictive pairs (e.g. AàB and BàC), but are frequently tested multiple times on inference trials without feedback during the critical test (e.g. AàC). Importantly, unlike in some associative inference paradigms, simply presenting the pairs for a choice without feedback does not provide any additional information concerning which face is of higher rank, and, more importantly, which pair has a higher growth potential.

### fMRI experimental procedure

Subjects participated in two days of scanning sessions more than a week apart (**Fig.1c**). The average time gap between the two sessions was 8.95 ± 0.81 days (±SE) (in the range of 7 to 19 days). On each day of the fMRI experiment session, participants performed three blocks of the partner selection task. A block consisted of 48 trials. Participants, therefore, needed to compute the GP of 576 pairs in total (48 trials ×2 pairs per trial ×3 blocks × 2 days) to make accurate decisions for the partner selection task. The pairs presented during the first-day scan were represented for the second-day scan in the reverse order. That is, the F0 and F1 pairs in the first-day scan were presented as the F0 and F2 pairs in the second-day scan and vice versa. This allowed us to test whether participants made correct decisions consistently as well as whether their brain consistently represented the direction of inferred trajectories (θ) over the abstract 2-D social hierarchy space across sessions acquired more than a week apart. While we randomized the trial order in each block, we also changed the order of blocks between the first day scan and the second day scans. That is, the trials tested in the blocks 1, 2, and 3 on the first day scan were tested in the blocks 5, 6, and 4, respectively on the second-day scan, which allowed us to generate unique comparisons for cross-validation across blocks (see the fMRI analysis).

### The frequency of the pairs presented for the partner selection task

16 individuals can be paired with 15 other individuals except for oneself. Therefore, there are 240 pairs that can possibly be generated from 16 individuals (since the vector angles of the inferred trajectory are different, we treated the AB pair and the BA pair differently) (**Fig.S2a**). Among 240 pairs, we carefully chose specific pairs for the partner selection task because we hypothesized participants may be less likely to use grid-like codes to compute the GP for some pairs where they could rely on a heuristic instead. For example, participants do not need to compute GP of two pairs if one of them includes an individual who has extreme ranks in both dimensions such as face 1 and 16 (**Fig.2h** shows the position of these faces; [P,C] = [1,1] and [4,4] respectively where 4 is the highest rank). In addition, if F0 is at the highest rank in one of two dimensions, participants only need to compare the rank of F1 and F2 in one dimension where F0’s rank was relatively lower. Considering the above, we sampled face 1 and 16 less frequently and were more likely to choose F0 from faces whose rank was not the highest in either dimension of the social hierarchy. **Fig.S2d** shows the 99 unique pairs presented for the partner selection task for the first time in the fMRI experiment and the frequency of the presentation of each of those pairs during the partner selection task. **Fig.S2e** shows the frequency of the inferred trajectories for the fMRI experiment categorized into 12 equal bins of 30° according to their direction (θ), which indicates that the 2-D social hierarchy space was well sampled.

### Post-scanning debriefing

After all the scans finished (on the second-day scan), we asked participants to describe the mental images that they had used to represent the interrelated 16 face stimuli and their decision strategy used to perform the partner selection task. All participants mentioned that when they see a face stimulus, they did not recall the rank in one dimension at a time, but they were able to recall the ranks in both dimensions simultaneously. Among 21 participants whose data were analyzed, only two participants reported the 4×4 social hierarchy structure in a 2-D space they used to represent the relationship. Other participants did not report being aware that this task can be solved spatially using an explicit 2-D representation of the social hierarchy.

### Post-scanning placement task

After participants reported their strategy, we asked them to perform a placement task. We asked participants to drag-and-drop each of the 16 faces of entrepreneurs to place on a 2-D space defined by the two social hierarchy dimensions according to their ranks. Note that all other behavior and imaging data were collected before this placement task. Importantly, participants were never explicitly asked to think about the relationship between individuals spatially before this placement task. Though most of the participants reported that they did not use a spatial representation to perform the task during post-scanning debriefing, all participants were successfully able to place the faces according to their social hierarchy ranks and reconstruct the social hierarchies in a 4×4 grid.

For visualization purposes, we rescaled the map of the faces placed by each participant by normalizing the longest pairwise distance between faces on the X coordinate and that on the Y coordinate to one (**Fig.2i**). Note that after rescaling, the location of faces can appear more precise at the boundary compared to the faces in the middle of the map. However, we confirmed that all participants reported an equidistant interval between any pair of neighboring entrepreneurs who differed by one rank in either the competence or popularity dimension.

### Behavioral analyses

We examined whether participants made a model-based decision using the learned structure of the social hierarchy instead of a model-free decision using heuristics as an alternative decision strategy. Specifically, we examined whether the decision of participants was better explained by the difference between GPs of two pairs (GP1-GP2), which requires participants to assemble the ranks of individuals in both dimensions compared with the difference in ranks or values that participants assigned to each individual, which does not require the internal representation of the social hierarchy to guide a decision.

As the first alternative way to make decision, we tested whether participants used an overall rank assigned to each individual in a combined dimension rather than ranks of each of two dimensions and chose the one who had greater overall rank between F1 and F2. That is, this model predicted that participants choose F1 if [*p*_*F*1_ × *G*_*F*1_] > [*P*_*F*2_ × *C*_*F*2_], and otherwise F2. This model differs from the GP-based decision in that the decision does not consider the rank of F0. As the second alternative model, we tested whether participants only use the ranks of F1 and F2 in one of two dimensions in which the rank of F0 was relatively deficient. For example, the decision of participants might only depend on the ranks of F1 and F2 in the popularity dimension when F0 was at a lower rank in the popularity dimension than their rank in the competence dimension. Formally, this model predicts that participants choose F1 if *C*_*F*1_ > *C*_*F*2_ given that *P*_*F*0_ > *C*_*F*0_ or if *P*_*F*1_ > *P*_*F*2_ given that *C*_*F*0_ > *P*_*F*0_, otherwise they choose F2. If participants use this heuristic, therefore, participants could have made sequential value comparisons, rather than using the inferred trajectories.

To test these alternative hypotheses, we included trials in which participants would fail to choose the correct partner if they adopted one of the alternatives. If a participant used the first heuristic, they would fail to be correct in 35 trials (24.3% of total trials). If a participant used the second heuristic, they would fail to be correct in 109 trials (75.7% of total trials). This allowed us to assess whether participants used the ranks of individuals in both dimensions to compute the expected GP, rather than using their ranks in only one dimension (**Fig.S3c**).

Next, we analyzed the reaction times (RT) of partner selection decisions. Notably, the accuracy of subject decisions was at ceiling and there was limited variability across trails or between subjects (mean accuracy was 98.68±0.27%; **Fig.S3c**), precluding analysis of variables explaining accuracy. We performed a multiple linear regression to explain the RT of decisions instead. The RT was measured from the F2 onset to the response. The following 11 regressors were inputted into the multiple linear regression: the Euclidean distance between F0F1 and F0F2 pairs, the absolute difference between them (*E*_*F*0*F*1_, *E*_*F*0*F*2_, and Δ*E* = |*F*_*F*0*F*1_ – *E*_*F*0*F*2_|), the GP of F0F1 and F0F2 pairs, and their absolute difference (*GP*_*F*0*F*1_, *GP*_*F*0*F*2_, and Δ*GP* = |*GP*_*F*0*F*1_ – *GP*_*F*0*F*2_|), the cosine angles of the trajectories *(cosθ*_*F*0*F*1_, *cosθ*_*F*0*F*2_, and *cosθ*_*F*1*F*2_), and whether the F0F1 and F0F2 vector was aligned to the EC grid orientation or not *(On_*F*0*F*1_* and *On*_*F*0*F*2_; where it was 1 for the on-grid pairs and θ for the off-grid pairs).

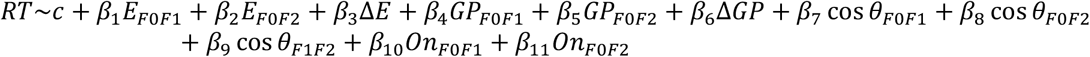

Due to the moderate correlation between the distances of trajectories (*E*) and the decision values *(GPs)* (r=0.47; **Fig.S4a**), we also performed the multiple linear regression while including *E_*F*0*F*1_* and *E*_*F*0*F*2_ after partialling out their covariance with GP_*F*0*F*1_ and *GP*_*F*0*F*2_. Group level effects of each of the distance measures were tested with a one-sample t-test to treat subjects as a random variable.

### MRI data acquisition

We acquired T2-weighted functional images on a Siemens Skyra 3 Tesla scanner. We used gradient-echo-planar imaging (EPI) pulse sequence with a slice angle of 30° relative to the anterior-posterior commissure line, minimizing the signal loss in the orbitofrontal cortex region (Weiskopf *et al*., 2006). We acquired 46 slices, 3mm thick with the following parameters: repetition time (TR) = 1300 ms, echo time (TE) = 24 ms, flip angle = 67°, field of view (FoV) = 192mm, voxel size = 3 x 3 x 3 mm^3^. Contiguous slices were acquired in interleaved order. To correct for deformations, we also acquired a field map with dual echo-time images covering the whole brain, with the following parameters: TR = 630 ms, TE1 = 10 ms, TE2 = 12.46 ms, flip angle = 40°, FoV = 192mm, voxel size = 3 x 3 x 3 mm^3^. We acquired a T1-weighted structural image using a magnetization-prepared rapid gradient echo sequence (MPRAGE) with the following parameters: TR = 1810 ms, TE = 2.98 ms, flip angle = 7°, FoV = 256mm, voxel size = 1 x 1 x 1 mm^3^.

### Pre-processing

The preprocessing of functional imaging data was performed using SPM12 (Wellcome Trust Centre for Neuroimaging). Images were corrected for slice timing, realigned to the first volume, and realigned to correct for motion using a six-parameter rigid body transformation. Inhomogeneities created using the phase of nonEPI gradient echo images at 2 echo times were coregistered with structural maps. Images were then spatially normalized by warping subject-specific images to the reference brain in MNI (Montreal Neurological Institute) coordinate space (2mm isotropic voxels). For the univariate analysis, images were smoothed using an 8-mm fullwidth at half maximum Gaussian kernel (Mikl *et al*., 2008).

### fMRI representational similarity analysis (RSA)

We performed a multivariate analysis using RSA (Kriegeskorte, 2008) to test whether the *a priori* regions of interest (ROIs) in the brain reflected the structure of the social hierarchy (**Fig.2a**). The ROIs were defined anatomically in the bilateral hippocampus (HC) (Yushkevich *et al*., 2015) and bilateral entorhinal cortex (EC) (Amunts *et al*., 2005; Zilles and Amunts, 2010) (**Fig.2b**). We also included additional ROIs in the bilateral primary motor cortex (M1) (Glasser *et al*., 2016) as control regions. Note that ROIs were defined independently from the current task.

By using RSA, we tested whether the brain integrates piecemeal learned relative status between entrepreneurs into a social hierarchy, and more importantly, whether the brain combines two social hierarchy dimensions, which were learned on different days, into a single 2-D representation. To address this question, we examined whether the level of pattern dissimilarity between the neural activity evoked by each entrepreneur was explained by a function of pairwise Euclidean distances between entrepreneurs on the 2-D social space. This analysis allowed us to test whether the brain had increasingly similar patterns of neural activity for entrepreneurs who were increasingly close to each other in the true 2-D social space.

To test our hypotheses, we first estimated *β* coefficients when each of the individual faces was shown at the time of F0, F1 or F2 using a GLM with a 2s box-car function in each of our anatomically defined *a priori* ROIs. We then averaged the unsmoothed *β* maps when the same face among 14 faces was presented across F0, F1 and F2 presentations, allowing us to estimate the patterns of neural activity in each ROI. We excluded two faces (1 and 16) who were sampled less frequently for the RSA analysis in order to match observations per face (see the pairs for the partner selection task section in methods). The reliability of the data was improved by applying multivariate noise normalization (Walther *et al*., 2016). Second, we quantified the representational similarity between the neural activity patterns elicited by face stimuli across three independent fMRI blocks acquired in the same day using the Mahalanobis distance. This pairwise dissimilarity further constructed a 14×14 representational dissimilarity matrix (RDM). These analysis steps were repeated per ROI. The separately estimated RDMs for each of the two-day scans were averaged across days. Third, we constructed a model RDM that contained the pairwise Euclidean distances between entrepreneurs on the 2-D social hierarchy space (**Fig.2a**). Last, we compared the neural RDMs estimated in each of the ROIs to our model RDMs using a rank correlation using Kendall’s *τ*_A_ (Kendall, 1938). Since we excluded two faces who had extreme ranks in both dimensions, the pattern dissimilarity associated with the pairs in which the Euclidean distance between them was 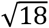 was only determined by one pair (face 4 and 12; [P,C] = [1,4] and [4,1] respectively). We therefore excluded the pattern dissimilarity estimated between them to ensure that the effects of Euclidean distance could not possibly be driven by the patterns to particular faces. Therefore, we examined to what extent the dissimilarity between activity patterns is explained by the Euclidean distance in the range of 1 to 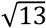. Statistical significance was calculated by constructing a null distribution by shuffling our data 1000 times at the subject level. Each subject’s *τ*_A_ was compared to the mean of this permutation distribution and submitted to a one-sample Wilcoxon signed-rank test across participants. We reported the results with the family-wise error rate (FWE) to correct multiple comparisons by the number of ROIs using the Holm-Bonferroni method (Holm, 1979).

To test more rigorously the effect of Euclidean distance, which can be factorized into a 1D distance of rank differences in the competence dimension and another 1-D distance of rank differences in the popularity dimension, we estimated the neural representation of the relationship between entrepreneurs in each of the two dimensions separately. We estimated the rank correlation (Kendall’s *τ*_A_) between the neural RDM which included neural activity patterns estimated from the same ROIs and the model RDMs which included the rank differences between entrepreneurs in each dimension. We further compared those rank correlations (*τ*_A_) that represent to what extent a ROI contained the representation of 16 entrepreneurs’ relationship in one of two social hierarchy dimensions, to test whether the neural representation of the relative ranks among entrepreneurs in one dimension was not different from that of the other dimension with the nonparametric two-sided Wilcoxon signed-rank test.

First, we performed RSA of 14 faces in the social hierarchy, while matching the number of samples per face, to test the effects of Euclidean distances on the neural activity patterns. To do that, we down-sampled by randomly choosing 12 trial responses associated with each of 14 faces acquired in each day. While down-sampling allows us to circumvent the potential concerns of unequal sampling, it does suffer from an overall small sample size. To complement this result, second, we performed the same RSA of 14 faces while including all trials’ neural responses. Third, we performed a control RSA in which the neural activity only acquired at the time of F0 presentation was included. Since some faces were selected less frequently as F0 during partner selection (See the pairs for the partner selection task in methods), we only include 8 individuals (**Fig.S5c**) for the control RSA. While this control analysis allows us to examine activity patterns while the brain activity was minimally modulated by other task-relevant cognitive processes, notably, the measurements of pattern similarity for F1 and F2 are likewise valid. Showing that the mean signals are hexadirectionally modulated with respect to F0 when presenting F1 and F2 does not imply the pattern of the activity across voxels cannot also reflect the relative position in the social hierarchy. For these reasons we include all three events (F0, F1, and F2) in the RSA analysis and present the new analysis of F0 as an additional result.

### Searchlight-based RSA

In addition to the ROI-based RSA, we performed whole-brain searchlight RSA to examine brain areas in which the dissimilarity between neural activity patterns associated with each of 14 faces is explained by the pairwise Euclidean distance between faces in the 2-D social hierarchy. We estimated the neural activity patterns elicited while each of the 14 faces was presented at the time of F0, F1, or F2 from each of the searchlights. Consistent with the ROI-based RSA, first, we performed the searchlight-based RSA while matching the number of samples of each face presentation. To complement the results of RSA based on the down-sampled responses, second, we also performed the RSA while including neural responses elicited from every trial. The dissimilarity matrices were quantified with the Euclidean distance between neural activity patterns estimated from different sessions acquired in the same day and averaged across days. Using the pairwise Euclidean distances as a predictor (**Fig.S2a**), we estimate the neural representational dissimilarity across searchlights with Kendall’s *τ*_A_ rank correlation. At the subject level, the effects of the Euclidean distances were further compared to a null distribution which was estimated from 1000 permutations while the positions of 14 faces were allocated randomly in the 2-D space. Specifically, each participant’s *τ*_A_ was compared to the mean of the permuted distribution and then was mapped back on the central voxel of each searchlight, which created a continuous map of the levels of representation of 2-D social hierarchy in the whole brain per subject. This map was further smoothed using an 8-mm full-width at half maximum (FWHM) Gaussian kernel and Fisher’s Z transformed. We further performed one-sample t-tests for group-level analysis. We reported the results corrected for multiple comparisons across the whole brain using TFCE (Smith and Nichols, 2009) with 1000 iterations of simulation (p_TFCE_<0.05). In addition, because we hypothesized that that cognitive map representations would be found in HC and EC based on prior studies (O’Keefe and Nadel, 1978; Eichenbaum and Cohen, 2014; Park *et al*., 2020; Whittington *et al*., 2020), we report results in the HC and EC corrected in the anatomically defined ROI, (Amunts *et al*., 2005; Zilles and Amunts, 2010; Yushkevich *et al*., 2015) which is independent of the current study (p_TFCE_<0.05).

### Multidimensional scaling

To visualize the representation of the social hierarchy structure built in HC and EC, we performed multidimensional scaling (MDS) analyses using the ‘midscale’ function in Matlab. The MDS allowed us to find the best way to arrange 14 individuals spatially in a 2-D space so that the distances between individuals in a 2-D space reflect their similarities between neural activity patterns. The distances between two individuals in the social hierarchy were measured by Mahalanobis distances between the neural activity extracted from the same HC and EC ROIs acquired in the same session. The dissimilarity matrix of each participant was averaged across sessions, across hemispheres in the HC and EC ROI separately, and averaged across participants. We used the first two eigenvalues to reconstruct the representation in a 2-D space.

With MDS, the locations of less sampled entities are more likely biased by noise compared to those of the entities sampled with higher frequency. Therefore, as can be seen below, the measures for face 1 and 16 in particular are expected to be less reliable than others. We deliberately sampled less frequently these faces to mitigate against subjects relying on heuristics during inferences, since Face 1 always lost on both dimensions and Face 16 always won on both dimensions. We therefore included 14 individuals from amongst 16 for the MDS analysis, excluding two faces who had extreme ranks in both dimensions. Moreover, we performed a MDS based on the activity patterns of the HC and EC while matching the number of sample responses associated with each of 14 individuals. Importantly, we apply MDS only for visualization of the group’s social hierarchy representation in the HC and EC, not for population inferences, where we instead rely on rank correlations of RSA. To complement the MDS based on equivalent sampling which suffers from an overall smaller sample size, we also performed the additional MDS while including all responses associated with 14 individuals. We only report the Spearman correlation coefficients, along with the above caveat about sampling frequencies. The numbers of presentations of each of 16 faces during a single scan (three blocks of one day) were shown in brackets as follows:

|  |  |  |  |
| --- | --- | --- | --- |
| Face 13 (42) | Face 14 (21) | Face 15 (16) | Face 16 (8) |
| Face 9 (23) | Face 10 (58) | Face 11 (58) | Face 12 (19) |
| Face 5 (69) | Face 6 (45) | Face 7 (57) | Face 8 (18) |
| Face 1 (12) | Face 2 (69) | Face 3 (18) | Face 4 (43) |

Last, we tested to what extent the pairwise Euclidean distance and pairwise cosine angle between 14 individuals in the MDS graph correlated with the true pairwise distance in the 4×4 social hierarchy structure. While an MDS can be acquired from the activity patterns of each participant, the MDS at the group level cannot be estimated by averaging multiple MDS due to differences in the scale and the orientation across individual MDS’s. Thus, we can only reliably acquire one MDS from the mean pattern dissimilarity of the group. For formally test to what extent the group MDS accounts for the true social hierarchy structure, we compared the level of correlation (Spearman’s *ρ*) between the pairwise distances between all possible pairs of 14 faces in the MDS and the pairwise distances estimated from the true social hierarchy and compared it to the distribution of the correlation coefficients that we computed from 1000 permutations while randomly locating those 14 faces in the 2-D hierarchy. In addition to the pairwise Euclidean distances, we also estimated how well the MDS captures the pairwise cosine angles of all possible pairs of the true social hierarchy. Using the same method, we further compared the correlation coefficient between pairwise cosine angles to the null distribution of the correlation coefficients (1000 random permutations). In addition to Spearman’s *ρ*, we report p-value based on the permutation.

### fMRI whole-brain analyses

In previous findings of rodent electrophysiology (Fyhn *et al*., 2004; Hafting *et al*., 2005; Doeller, Barry and Burgess, 2010), grid cells in the entorhinal cortex (EC) showed greater activity for navigating in the direction aligned with the mean grid orientation *(ϕ)* across grid cells compared to them for navigating in the direction misaligned with *ϕ,* which generated a specific pattern of sixfold symmetry (**Fig.1b**). Recent studies have shown that a hexagonally symmetric grid-like code can be measured in the blood oxygen-level-dependent (BOLD) signal with fMRI in humans (Doeller, Barry and Burgess, 2010; Constantinescu, O’Reilly and Behrens, 2016; Julian *et al*., 2018; Nau *et al*., 2018; Bao *et al*., 2019). We leveraged these established fMRI markers to test whether a gridlike code is used for inferred trajectories between entities in an abstract cognitive map to guide decision-making. Specifically, we searched for neural evidence for hexa-directional modulation of inferred decision trajectories through the reconstructed cognitive map of a 2-D social hierarchy that we hypothesized were used to compute the growth potential (GP) to guide decisions in the partner selection task: The 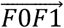 vector at the time of F1 presentation and the 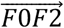 vector at the time of F2 presentation (**Fig.1e,f**).

To address this, we implemented a series of general linear models (GLM) to analyze fMRI data. First, we performed a whole-brain analysis to identify brain areas in which activity showed hexagonal symmetrical patterns according to the direction of the inferred trajectories (θ) (GLM1). Importantly, GLM1 was performed without any assumption of the subject-specific mean grid orientation *(ϕ).* Based on the results of GLM1, we define regions of interest (ROIs) in the brain in unbiased way for GLM2. Second, in GLM2, we tested the effects of hexadirectional modulations according to *θ* aligned with *ϕ* using a leave-one-and-out cross-validation (CV) procedure. Specifically, in GLM2, we estimated the putative grid orientation *(ϕ)* from one dataset (either two of 3 runs for within-day analyses or 1 of 2 days for between-day analyses) using an ROI defined independently based on GLM1 and applied the estimated *ϕ* to the other remaining dataset to test for the hexadirectional grid-like representation across the whole brain. Third, with GLM3, we identified brain areas encoding the variables guiding the partner selection decision including the growth potential (GP) and IGP1-GP2I, which is independent from both *θ* and *ϕ.* In addition, with GLM3, we further examined the relationship between neural encodings of the decision variables (GP) and individual differences in the effects of hexadirectional modulation of the inferred trajectories while controlling the overall accuracy of decision making. Last, with GLM4, we tested for complementary evidence of greater GP effects for the aligned compared to misaligned trajectories, consistent with a role of hexadirectional modulations in EC for guiding map-based decision making.

#### Defining a region of interest in an unbiased manner (GLM1)

With GLM1 we identified brain areas where activity showed hexagonally symmetric patterns according to the direction of the inferred trajectories (θ) but did not depend on the grid orientation *(ϕ).* The purpose of GLM1 was to functionally define ROIs in a way that was unbiased from the putative grid orientation *(ϕ)* for future tests (see GLM2). GLM1 contained separate onset regressors for the onsets of F0 and the partner stimuli presentations (combining F1 and F2 presentations). The BOLD responses of those regressors were modeled with the 2.5s boxcar function. The regressors of F1 and F2 presentations were modulated with two parametric regressors which include the sine and cosine of the direction of inferred trajectory (*θ*) with a sixfold periodicity, that is, *sin(6θ)* and *cos(6θ).* These regressors produced parameter estimates, *β_sin_* and *β_cos_* with amplitude 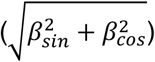 for brain regions that are sensitive to hexagonal symmetry. When a partner face stimulus (Fx) followed F0, the direction of inferred trajectory *θ_F0Fx_* was defined as follows:

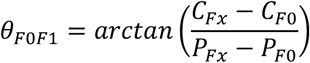

where *C_Fx_* and *P_Fx_* indicate the level of competence and popularity dimensions of the individual, Fx respectively. We included the onset of button presses with a stick function as additional regressors of no interest to account for the potential motor related effects. The 6 motion parameters obtained during realignment were also entered into GLM1 as regressors of no interest. The orthogonalize function was turned off. All these regressors were convolved with the canonical hemodynamic response function (HRF). Using an F-test, we identified the brain areas in which BOLD signals were significantly modulated by a weighted sum of two regressors, *βsin × sin(6θ)* + *βcos × cos(6θ).* This method we adopted here was established in a previous study (Constantinescu, O’Reilly and Behrens, 2016). All these regressors in these GLMs were convolved with the canonical hemodynamic response function (HRF). Individual whole brain contrast maps were z-scored and inputted to a one-sample t-test to test for group level effects using the z-statistic based on an asymptotic approximation (Jenkinson and Woolrich, 2004).

Considering that the F-statistics could be overestimated if the variance of the first level statistic was under-estimated, we only used GLM1 to create ROIs in an unbiased manner, but not for statistical inference. We defined ROIs for future tests in the entorhinal cortex (EC) (peak MNI coordinates at [x,y,z]=[26,-2,-36], z=2.62) and medial prefrontal cortex (mPFC) (peak MNI coordinates at [x,y,z]=[2,66,-4], z=5.58) (right panel in **Fig.S2a**). We defined the ROIS in the EC and mPFC based on the previous finding showing grid cells in these areas during virtual navigation in humans using pre-surgical single unit recordings (Jacobs *et al*., 2013) and previous grid-like fMRI effects in these regions (Doeller, Barry and Burgess, 2010; Constantinescu, O’Reilly and Behrens, 2016). Based on the F-test results, we defined ROIs in the EC and mPFC within their anatomical masks defined in EC (Amunts *et al*., 2005; Zilles and Amunts, 2010) and mPFC (Neubert *et al*., 2015), while including the voxels showing the effects of six-fold symmetry at the threshold z > 2.3 which corresponds to p = 0.01.

#### Testing hexadirectional modulation aligned to the grid orientation (GLM2)

We performed the GLM2 to identify brain areas whose activity was modulated by the direction of inferred trajectories (θ) with six-fold symmetry aligned with the grid orientation *(ϕ).* To disentangle the effects of grid orientation *(ϕ)* from the direction of inferred trajectories (θ), we used a crossvalidation (CV) procedure. This allowed us to identify hexadirectional modulations with consistent grid orientation *(ϕ)* across sessions.

The CV was performed as follows. We estimated *ϕ* from a separate dataset of each participant (called estimate set) and applied the participant-specific parameter *ϕ* to the other dataset (called test set) to test the effects of θ. For instance, the effect of θ in block 1 was tested while applying *ϕ* estimated from blocks 2 and 3 acquired on the same day scan. Likewise, the *ϕ* for block 2 was estimated from blocks 1 and 3; and the *ϕ* for block 3 was estimated from blocks 1 and 2 (bottom left panel in **Fig.S2c**). The CV was possible because whereas the direction of inferred trajectories 0 was consistent for the same trajectory across participants but varied across trajectories, the grid orientation *ϕ* was relatively consistent within a participant but varies across participants (see the top panel in **Fig.S2c**). We estimated the mean grid orientation *ϕ* in the functionally defined ROIs in the EC defined based on GLM1 (**Fig.S2b**), by adopting the same method proposed in previous studies (Doeller, Barry and Burgess, 2010; Constantinescu, O’Reilly and Behrens, 2016; Nau *et al*., 2018). Specifically, we estimated *ϕ* in each voxel of the ROIs using the *β* coefficients for the sine and cosine regressors *(β_sin_* and *β_cos_*) in GLM1 as follows:

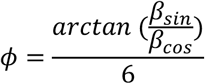

The mean grid orientation *(ϕ)* across each voxel in each of the ROIs was estimated using the circular mean (Berens, 2009), which was divided by six to compute *ϕ* in a range between 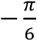 to 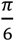, thus taking the six-fold periodicity into account. We expected that the estimated grid orientations are not randomly distributed across voxels but show a non-uniformity of circular distances across voxels. We tested if there was a non-uniform clustering of putative grid orientations estimated from different sessions across voxels in each participant using Rayleigh’s tests.

In GLM2, we modeled the BOLD signals of onsets of the partner stimuli (combining F1 and F2 presentations) with a 2.5 boxcar function. These events were modulated by a parametric regressor, cos(6[θ – *ϕ*]), where *θ* was the hypothesized direction of inferred trajectories between individuals in each pair and *ϕ* was the mean grid orientation estimated from the separate estimate dataset. Additionally, the onsets of F0 and button presses were modeled as separate regressors with a stick function. The 6 motion regressors were also entered as regressors of no-interest. The orthogonalize function was turned off. All regressors in these GLMs were convolved with the canonical HRF. Last, the group-level effects were tested with a one-sample t-test.

In addition to the grid orientation consistency across sessions acquired within the same day, we further tested for the grid orientation consistency across sessions acquired on different days. Again, to test the effect of hexadirectional modulations in a statistically unbiased manner, we used a CV procedure. Specifically, the effect of the was tested on the fMRI sessions acquired in one of two days scans while applying the grid orientation, *ϕ* estimated in the EC activity from a session acquired in the other day scan (**Extended data Figure 3c**). This additional analysis allowed us to test whether the brain activity had the hexadirectional modulations in one day consistently showed the same effects to indicate the inferred trajectories in a different day. That is, we tested for a stable representation of the learned social hierarchy in the brain more than a week later.

#### Identifying neural encoding of decision-related variables (GLM3)

To make accurate decisions in each trial of the partner selection task, participants were required to compute GP1 for the F1 presentation and GP2 for the F2 presentation. Specifically, the GP of each pair corresponds to the size of the rectangle that is drawn by the rank positions of two entrepreneurs in the abstract 2-D social hierarchy space (as **Fig.1f** shows). Subsequently, the decision should be guided by the difference between the two GPs. To capture this value comparison term, we use the absolute difference in GPs (Δ*GP* = |*GP*_*F*0*F*1_ – *GP*_*F*0*F*2_|).

We performed the GLM3 to examine the neural encodings of decision variables (GP and *ΔGP)* which guide the partner selection decision. We modeled the BOLD signals of onsets of the partner stimuli (combining F1 and F2 presentations) with a 2.5 boxcar function which were subsequently modulated by a parametric regressor, GP. We also modeled the BOLD signals at the time of decision-making with a stick function which was modulated by a parametric regressor, Δ*GP*. Additionally, the onset of F0 and the 6 motion regressors were also modeled with separate regressors in the same way of the GLM1. The orthogonalize function was turned off. All regressors in these GLMs were convolved with the canonical HRF.

The GP was dependent on the relative ranks of two entrepreneurs in both dimensions, but independent from the directions of the inferred trajectories (**Fig.S10a**, mean r±SE = 0.04±0.03). To rigorously test whether these two regressors explained independent variance in neural activity, as a control analysis, we performed an additional GLM in which the BOLD signals at F1 and F2 presentations were modulated by two parametric regressors, *cos(6[θ-ϕ])* and GP, in the same model. The effects of each of those two regressors were further compared with the effects of *cos(6[θ-ϕ]*) in GLM2 and the GP effects in GLM3 to examine whether the effects of two regressors remained while controlling for the effect of the other regressor.

#### Interaction effects between alignments to the grid orientation and GP computations (GLM4)

In GLM 4 we tested the hypothesis that the GP effects depend on the alignments of the inferred trajectories to the EC grid orientation *(ϕ)* by comparing the GP effects between aligned trajectories and misaligned trajectories. Note that there was not a significant difference between GP regressor values for aligned and misaligned trajectories (**Fig.S10b,c**). In GLM4, we modeled the BOLD signals at the onset of the partner stimuli (combining both F1 and F2 presentations) separately for the trajectories aligned with the grid orientation *(ϕ)* from those misaligned trajectories with a 2.5s boxcar function. Both regressors were parametrically modulated by GP of the pairs. Consistently, we applied the same putative grid orientation, *ϕ* estimated in the EC ROI using a CV across sessions acquired within the same day, as before. Additionally, the onsets of F0 presentation and button presses were modeled with separate regressors with a stick function. The 6 motion parameters obtained during realignment were entered into the model as regressors of no interest. The orthogonalize function was turned off. All these regressors were convolved with the canonical HRF. We performed a contrast analysis between the GP effects in the aligned trajectories and those in the misaligned trajectories. The individual contrast maps were entered into a one-sample t-test to test group-level effects.

#### Group-level inference

For all analyses using a GLM (GLM1,2,3, and 4), group-level effects were tested with a one-sample t-test. First, we performed group-level inference on *a priori* task-independent EC (Amunts *et al*., 2005; Zilles and Amunts, 2010) and HC (Yushkevich *et al*., 2015) ROIs which were defined anatomically. Second, we performed exploratory whole-brain analyses. To correct for multiple comparisons, we used permutation-based threshold-free cluster enhancement (TFCE) (Smith and Nichols, 2009). The medial EC has famously been found to contain grid cells, and is thought to be their primary source (Hafting *et al.*, 2005). A signature of these grid cells in EC has been found in recent human fMRI studies, and thus a central hypothesis of our study was that EC would show grid-like coding. Specifically, these recent studies show EC grid-like coding both for spatially navigating virtual environments (Doeller, Barry and Burgess, 2010) and also for tracking the abstract task state defined by the continuously morphing neck and leg lengths of a bird stimulus (Constantinescu, O’Reilly and Behrens, 2016). The EC activity in these studies was modulated by the direction of the vector defined on a 2-D space with hexagonal patterns. The EC is, therefore, the primary ROI to test our main hypothesis in the current study (note that all vectors are aligned to the EC grid orientation). In addition, the anatomical geometry of the EC is not ideal for cluster-based inference such as TFCE. For those reasons, we used the anatomically defined ROI-based TFCE correction for statistical inference of the grid effect in the EC (p_TFCE_<0.05 corrected in the ROI). We report activations (**Fig.2,3,4**) that are cluster-corrected at the threshold p_TFCE_ < 0.05 over the whole brain for all brain regions, except for the EC, where we used a threshold of p_TFCE_ < 0.05 corrected within anatomically defined ROI, due to our strong a priori hypothesis of grid coding in EC.

### Region of interest analyses

#### Specificity for six-fold symmetry in ROIs

We conducted the regions of interest (ROI) analyses to test the hexadirectional patterns in alignment with the EC grid orientation *(ϕ)* within unbiased ROIs. Moreover, we visualized the sinusoidal modulations (**Fig.1c**) in the activity in these ROIs as a function of the direction of inferred trajectories (θ) with six-fold periodicity. In addition, we performed control analyses to test whether the periodic effects in the ROis were specific to six-fold but not to other control periodicities (four-, five-, seven- and eight-fold).

First, we tested the hexadirectional modulations on *a priori* ROis in the EC and mPFC which we defined based on the GLM1 (**Extended Data Figure 3b**) independently from the grid orientation *(ϕ).* We tested this effect also on the anatomically defined EC and mPFC ROis as well ((Amunts *et al*., 2005; Zilles and Amunts, 2010) for EC; (Neubert *et al*., 2015) for mPFC) (**Fig.S8a**) to confirm the effects did not depend on how we defined the ROis. Second, because we used faces as stimuli in our task, we tested the hexadirectional modulations in the ROis defined in the fusiform face areas (FFA). Note that the FFA was specifically engaged with the current task in response to the face stimuli. To define FFA ROis independently from *ϕ,* we included the brain area activated in response to the face stimuli compared to fixations. The inclusive mask was defined at the threshold, t_20_>3.6 which corresponds to p<0.001, uncorrected (peak MNI coordinate [x,y,z]=[16,-34,-6], t=10.94 for right FFA and [x,y,z]=[-18,-30,-10], t=9.81 for left FFA; **Fig.S8b**). Third, we defined ROIs in the brain areas where activity quantitatively encoded the GP, which included the bilateral TPJ and mPFC based on the GLM3 (**Fig.4a**; **Table S3a**). The inclusive mask was defined at the threshold, t_20_>3.6 which corresponds to p<0.001, uncorrected. With the TPJ and mPFC ROIs, we tested whether the neural encodings of GP *(β* GP) were modulated by the direction of the inferred trajectories (θ) with the six-fold periodicity aligned to the EC grid orientation *(ϕ).* Note that the functional definitions of the above ROIs were all independent from the subsequent statistical tests performed for inference.

To estimate the trajectory-by-trajectory neural activity, we modeled the BOLD signals at the time of onsets of every presentation of F1 and F2 stimuli as separate regressors with a 2.5s boxcar function. These event regressors were also modulated by a parametric regressor for the GP. The onsets of button presses were modeled with a separate regressor with a stick function. The 6 motion parameters were added as regressors of no-interests. All regressors were convolved with the HRF.

First, we extracted the BOLD signals activated at the time of every presentations of a partner stimulus (including both F1 and F2) in the EC, mPFC, and FFA ROIs. Seconds, we extracted the series of *β* signals encoding GP at the time of every partner stimulus presentation in the bilateral TPJ and mPFC ROIs. The series of the extracted BOLD signals and *β* GP were separately z-scored per ROI within each block. Based on the subject-specific EC grid orientation *(ϕ)* estimated with the CV across sessions acquired within a day, we categorized the z-scores into 12 equal bins of 30° according to the direction of the inferred trajectory (θ). The z-scores of each participant were then averaged within each of 12 bins ±SE. This procedure produced six mean activity of the aligned trajectories and six mean activity of the misaligned trajectories per ROI.

In each of the ROIs, we performed nonparametric one-sided t tests to test the effects of hexadirectional modulations which were measured as differences in z-scores (the aligned – misaligned trajectories’ bins). In addition, to test for the specificity of the six-fold periodicity, we performed the additional control analyses. Specifically, we performed two-sided paired t tests to compare the differences in z-scores computed using the model of the six-fold periodicity (effects of hexadirectional modulations) to the differences in z-scores computed using the other model assuming the different level of periodicity including four-, five-, seven- and eight-fold (effects of modulations in other alternative periodicities). Last, we performed leave-one-bin-out (LOBO) analyses which allowed us to confirm that the effect of hexadirectional modulations in *βGP* was not driven by the activity of any specific bin. Specifically, with the LOBO analysis, we performed one-sided t-tests to examine whether the effects of hexadirectional modulations (difference in mean z-scores between the aligned and misaligned trajectories) were still greater than zero when the effects were computed from only 11 bins while excluding each of 12 bins (**Fig.S10f**).

#### Testing hexadirectional modulation for novel pairs

As mentioned in behavioral training, participants experienced 2 or 3 training sets of the partner selection task without feedback before the fMRI session. While the trials presented during behavioral training were not presented again in the fMRI, some pairs (F0 and F1 or F2) that had been presented during behavioral training (always in the absence of feedback) were presented again with other pairs during fMRI. The number of these overlapping pairs differs across participants, depending on the number of training sets they experienced. Out of 99 unique pairs, we found 11 pairs were presented during both fMRI and behavioral training for 17 participants, and 16 pairs were presented during fMRI and behavioral training for 4 participants. Since no feedback was given to participants during behavioral training and fMRI, we reasoned that no learning process was involved during training on the partner selection task. However, it is possible that participants memorize the value of growth potential of some pairs when they were presented during behavioral training and recall the GP when the same pair was shown again in fMRI, even though they needed to compare it with a new pair to make a novel decision. If participants use their memorized decision values, it is also possible that participants may retrieve the GP computed in the previous trial to guide their decision in the current trial during fMRI. To test whether the gridlike code plays a role when making inferences about entirely novel pairs, we performed an additional ROI analysis in which we tested for a grid-like code only for the pairs that are presented for the first time during fMRI. In this control analysis, we include only the z-score transformed beta extracted from the EC and mPFC ROI from the 88 (83) remaining pairs that were shown for the first time to each participant during the first day of scanning (Day1 scan). We also tested the effects of hexadirectional modulations on *β* GP in the TPJ and mPFC ROIs aligned to the EC grid orientation *(ϕ)* only for those 88 (83) pairs when they were presented for the first time. **Extended Data Figure 4a** shows the 88 (83) pairs that we used for this analysis.

#### Between-subject relationship between hexadirectional modulations in EC and the GP effects

To test for a relationship between hexadirectional modulation and GP effects, we addressed the following question: Do those participants who had a greater hexadirectional modulation in EC and mPFC also show greater neural encoding of GP? The underlying rationale for this hypothesis is as follows. If the grid-like codes in the EC and mPFC are important for accurate representations of inferred trajectories, it might further serve for accurate computations of the decision values in decision-making guided by a cognitive map. To test this hypothesis, we defined the subject-specific “gridness” score as to what extent the EC and mPFC signals were modulated by the direction of inferred trajectories in six-fold periodicity. Therefore, the gridness score equals to the *β* cos(6[θ-*ϕ])* in EC and in mPFC, and then we tested whether individual differences in neural GP effects were explained by gridness scores. Specifically, using GLM3, we entered individual contrast maps of GP into a one-sample t-test to test the group-level effects and also entered the EC and mPFC gridness scores as (demeaned) covariates for the second-level regression analysis across participants. In addition to the gridness score, we also include the overall accuracy normalized in a range of 0 to 1 as an additional covariate to control for the choice accuracy. To test the between-subject effect in the brain areas engaged in the tasks requiring perspective taking, we tested this effect within the anatomically defined independent ROIs including bilateral TPJ (Mars *et al*., 2012) and mPFC (Neubert *et al*., 2015) (**Fig.4c**).

#### The GP effects modulated by the inferred trajectories aligned to the EC grid orientation

We examined the effects of grid alignments in EC for making inferences of direct trajectories between individuals in the abstract social space on the activity in the brain areas encoding GP. We defined the ROIs in the mPFC and bilateral TPJ in which their activity correlates with GP while presenting F1 or F2 stimuli (GLM3) within the inclusive mask defined at the threshold t_20_>3.6 which corresponds to p<0.001, uncorrected (**Fig.4a**; **Table S3a**). In the ROIs, we extracted the *β* GP of the aligned trajectories and the *β* GP of misaligned trajectories which indicated to what extent the BOLD signals encoded the GP when the inferred trajectories were aligned and that for the trajectories misaligned to the EC grid orientation *(ϕ).* Note that the GP was independently defined from the direction of inferred trajectories (θ) regardless of the EC grid orientation *(ϕ)* (**Fig.S10a,b,c**, mean r±SE = 0.04±0.03). The GP, therefore, also independent whether the inferred trajectory was aligned or misaligned which obviates the concerns of the ROI selection biases. For each ROI, we performed a one-sample t test to confirm their activity encode GP *(β* GP > 0) but also a paired t-test to test whether the *β* GP was greater for the trajectories aligned to the EC grid orientation compared to the *β* GP of misaligned trajectories (**Fig.4e**).

**Extended Data Figure 1.**
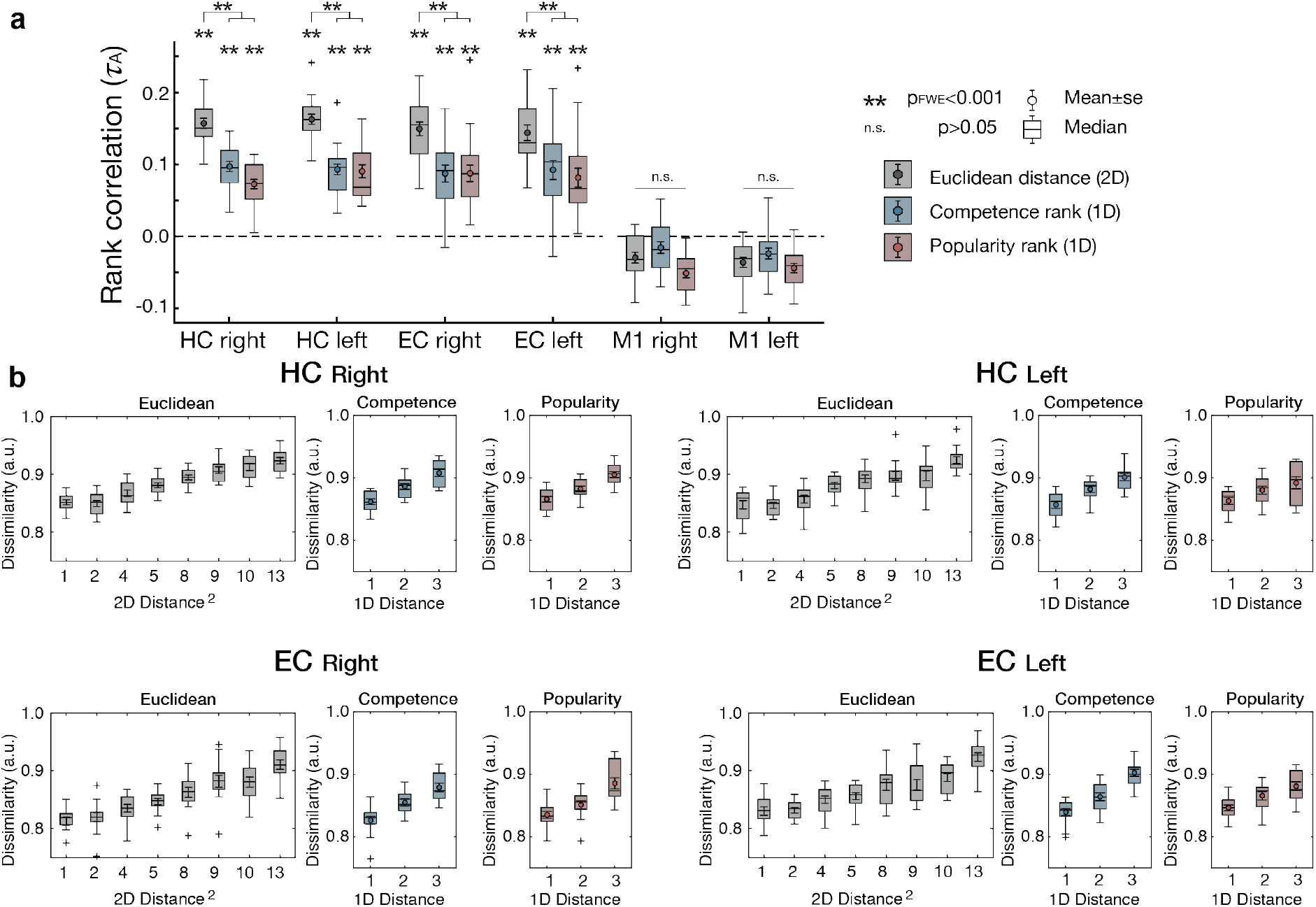
Representational similarity analysis (RSA) including all events of face stimuli presentation. **a.** RSA including neural responses to all the events associated with 14 individuals in the social hierarchy. Consistent with the results of the RSA based on downsampling shown in **Fig.2c**, we found effects of Euclidean distance on the pattern dissimilarity in the HC and EC but not in M1. The HC-EC system utilizes a 2-D relational cognitive map to represent the social hierarchy rather than representing 1-D map (*τ*_A_ of Euclidean distance (gray) > *τ*_A_ of one-dimensional rank distance in competence dimension (blue) and in popularity dimension (red)). **, p_FWE_<0.001 after correction for the number of bilateral ROIs (n=4) with the Bonferroni-Holm method; n.s., p>0.05, uncorrected. **b.** The dissimilarity between activity patterns estimated in bilateral HC and EC increases in proportion to the true pairwise Euclidean distance between individuals in the 2-D abstract social space (left, gray). The dissimilarity between activity patterns increases not only with the 1-D rank distance in the competence dimension (middle, blue) but also the 1-D rank distance in the popularity dimension (right, red). Methods and notation are identical to **Fig.2**.

**Extended Data Figure 2.**
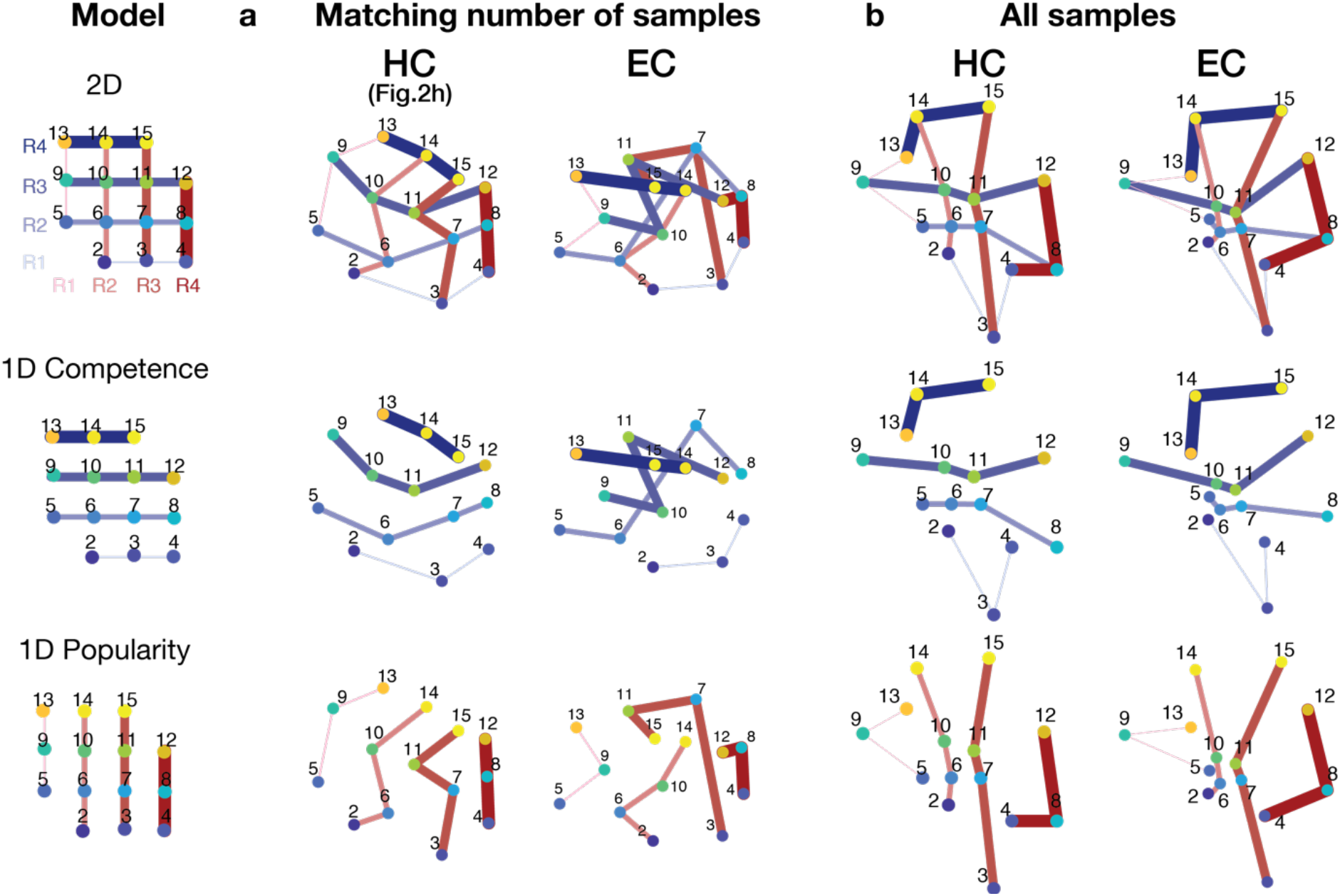
Multidimensional scaling (MDS) Visualization of the group representation of the social hierarchy in a 2-D space using MDS on the neural activity extracted from the HC and EC ROIs. There were considerably fewer presentations of individuals at the rank positions 1 and 16, making their estimates less reliable, and so these less frequently sampled individuals were excluded in computing the MDS. The 2-D representations (top) can be factorized into two 1-D hierarchies: competence (middle) and popularity (bottom). The lines indicate the individuals at the same rank in the social hierarchy. The thicker the line, the higher the rank in the given dimension. **a.** The MDS computed from the mean pattern dissimilarity across participants after matching the number of samples per face. Numbers indicate face position as shown in the true hierarchy in the Model on the left. Blue colors correspond to competence the dimension and red to the popularity dimension. The distances and angles between the estimated individual locations in the HC and EC MDS are significantly correlated with the pairwise Euclidean distances (Spearman’s *ρ*= 0.84 for HC and *ρ*= 0.63 for EC) and cosine angles *(ρ* = 0.93 for HC and *ρ* = 0.71 for EC) in the true 4×4 social hierarchy structure, compared to random configurations (both p<0.01 compared to 1000 random permutations). **b.** The MDS estimated from the mean neural activity patterns including every presentation of the 14 face stimuli. Associated with **Fig. 2g** and **Extended Data Figure 1**.

**Extended Data Figure 3.**
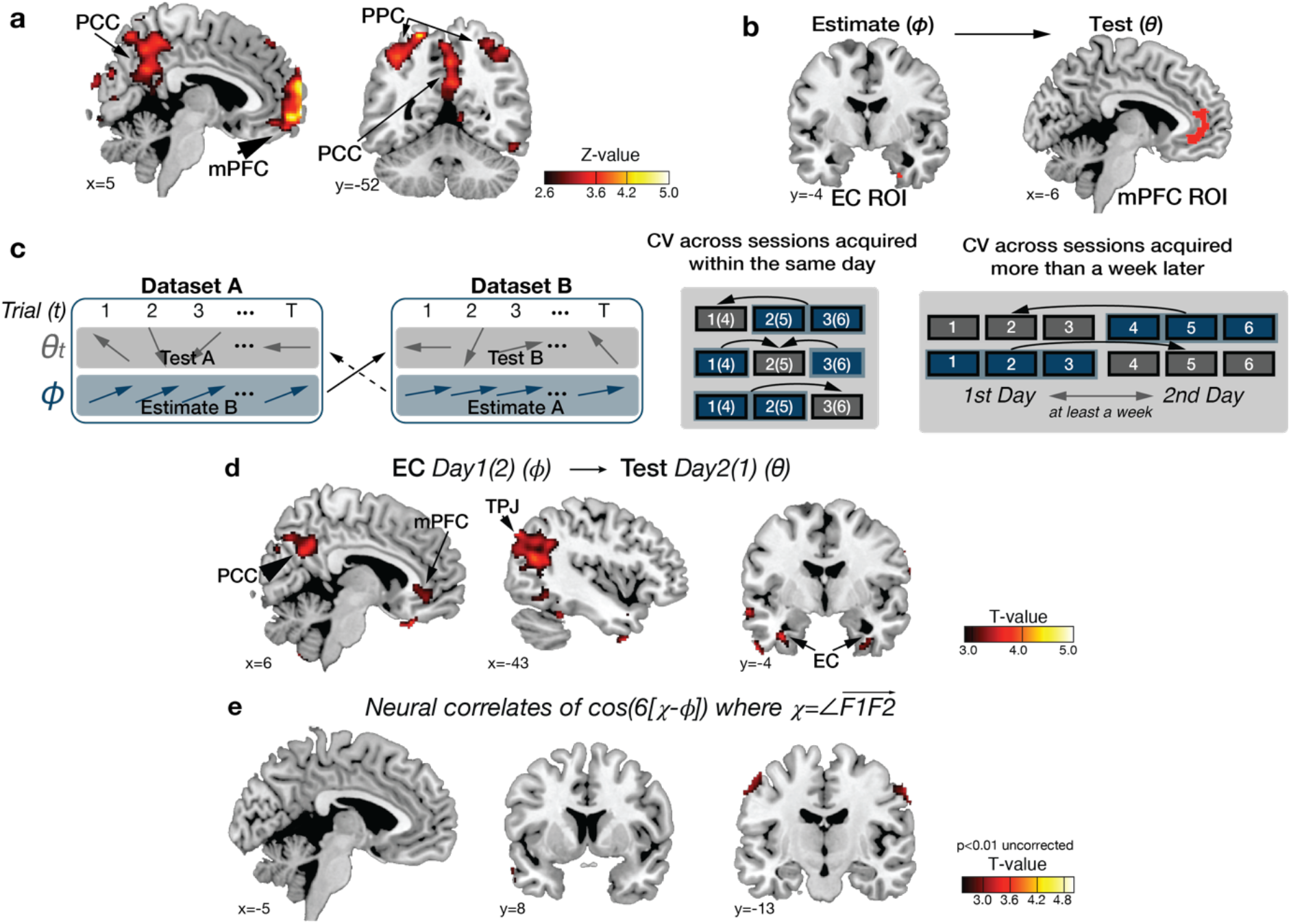
The analysis procedure to examine the effects of grid-like codes. **a.** Identifying brain regions sensitive to hexagonal symmetry across the whole brain without aligning to the entorhinal grid orientation, *ϕ.* To identify hexagonally symmetric signals, we adopted previously developed methods from a previous study (Constantinescu, O’Reilly and Behrens, 2016). We used a Z-transformed F-statistic to examine the BOLD activity modulated by any linear combination of *sin(6θ)* and *cos(6θ).* Hexagonal modulation was found in brain regions including in medial prefrontal cortex (mPFC; peak MNI coordinate, [x,y,z]=[2,66,-4], z=5.58), posterior cingulate cortex/precuneus (PCC; [2,-50, 36], z=3.19), bilateral posterior parietal cortex (PPC; [36,-46, 62], z=4.27 for right; [-38,-50, 50], z=3.81 for left), left lateral orbitofrontal cortex (OFC; [-42,44,8], z=3.79), and retrosplenial cortex (RSC; [x,y,z] = [2,-54,28], z = 3.45) at a threshold p_TFCE_<0.05 (whole brain TFCE correction), as well as the right entorhinal cortex (EC; [26, −10, −40], z=2.80) at a threshold, p_TFCF_<0.05 (corrected within *a priori* anatomically defined EC region of interest (ROI) (Amunts *et al*., 2005; Zilles and Amunts, 2010)). For visualization purposes, the maps are thresholded at z>2.6 (p<0.005 uncorrected). **b.** We did not use the results of F-test for statistical inference but to functionally define ROIs in EC and mPFC for future tests. The ROIs were defined within the anatomically defined masks in the EC (Amunts *et al*., 2005; Zilles and Amunts, 2010) and mPFC (Neubert *et al*., 2015) by including effects at a threshold of z>2.3, which corresponded txo p<0.01. Using these independently defined ROIs, we then tested if the grid orientation estimated in the ROI was consistent across separate fMRI sessions in an unbiased way. It is important to note that the ROIs were defined from results of a statistically independent analysis, which was not dependent on the grid orientation. **c.** Illustrations of the cross-validation (CV) procedure. By splitting a dataset for estimating the grid orientation, *ϕ,* from another dataset to test for hexagonal modulation for inferred trajectories, *θ,* we could test for brain regions where activity was modulated by cos(6[ε-ϕ]) in alignment with the grid orientation estimated from the independent dataset for each participant. The CV was possible because the grid orientation, *ϕ,* is thought to be relatively stable but different across participants, whereas the direction of inferred trajectories *θ* varies only according to the position of F0, F1, and F2 (left panel). We performed a CV procedure both from fMRI sessions acquired within the same day (middle panel), and also from fMRI sessions acquired more than a week apart (right panel). **d.** Consistency of grid orientation in EC across sessions acquired more than a week apart. Concretely, in alignment with the EC grid orientation estimated from sessions acquired on a different day, we found hexadirectional modulation in a network of brain regions, including the mPFC ([−2, 36, −8], t=5.04), PCC ([6,−58, 30], t=3.68), and TPJ ([−36, −64, 22], t=4.06) at our whole brain corrected threshold p_TFCE_<0.05, as well as in EC ([36, −10, −38], t=4,21) at p_TFCE_<0.05, small-volume-corrected in our *a priori* EC ROI (**Table S2b**). **e.** We estimated the angle of the F1 F2 vector *(χ)* and inputted the cos(6[*χ-ϕ*]) at the time of F2 presentation as an additional parametric regressor into GLM2. We did not find any brain areas encoding the angles of F1F2 vectors, except at a reduced threshold, in the bilateral somatosensory cortex ([x,y,z] = [-52,-14, 56], t=3.33 and [58,-8,50], t=3.17, p<0.005 uncorrected).

**Extended Data Figure 4.**
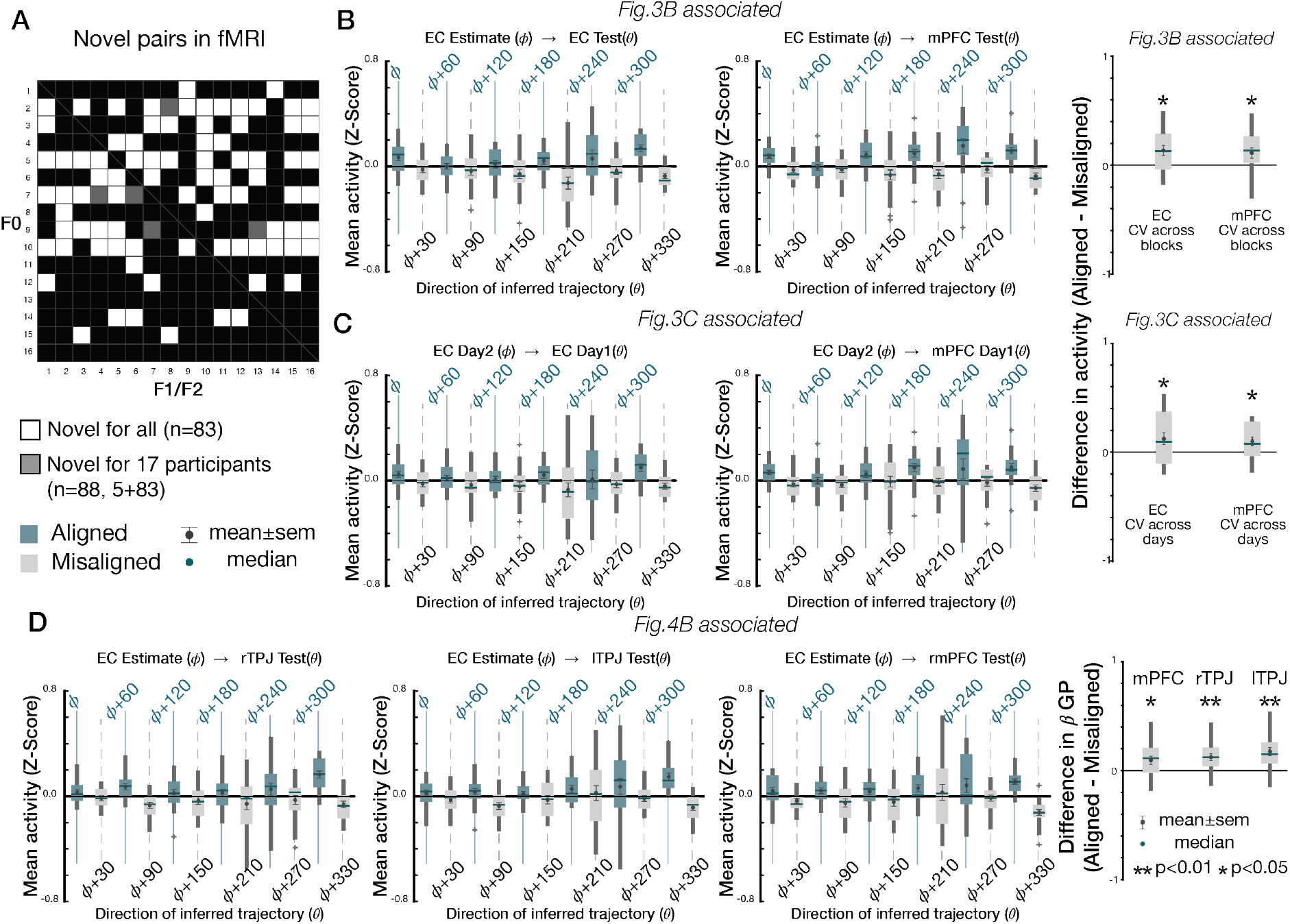
Hexagonal modulation for inferred trajectories only for the novel pairs when presented for the first-time. **a.** Among all the 88 (83) pairs (F0-F1 and F0-F2) presented during the partner selection task, those pairs that were not presented during behavioral training (always in the absence of feedback) but presented during fMRI for the first time are shown: 83 pairs in white were presented for the first-time for 4 participants; 88 pairs in white and gray were shown for the first-time for 17 participants. The grid effects were tested only for those pairs presented for the first time during the day1 scan. We extracted the mean activity and GP effects for each bin, restricted to when each pair was presented for the first time to participants. **b and c.** Associated with **Fig.3b and c**. The mean EC (left panel) and mPFC (middle panel) activity in 30° bins aligned to the EC grid orientation estimated from different blocks acquired in the same day’s scan (**b**) and the EC grid orientation acquired from a different day’s (day 2) scan (**c**) with six-fold symmetry. Right panel shows formal comparison of trajectories aligned and misaligned with both methods of computing the EC grid orientation. We found greater activity for the aligned pairs compared to the misaligned pairs to the EC grid orientation in EC and mPFC ROIs (one-sample t-test). **d.** Associated with **Fig.4b.** The GP effects in mPFC and bilateral TPJ are modulated by the grid alignment of the inferred trajectories aligned with the EC grid orientation. The GP effects are greater for the aligned pairs compared to misaligned pairs, even when they were presented for the first time (one-sample t-test). Box, lower and upper quartiles; line, median; whiskers, range of the data excluding outliers; +, the whisker’s range of outliers. **, p<0.01; *, p<0.05.

**Extended Data Figure 5.**
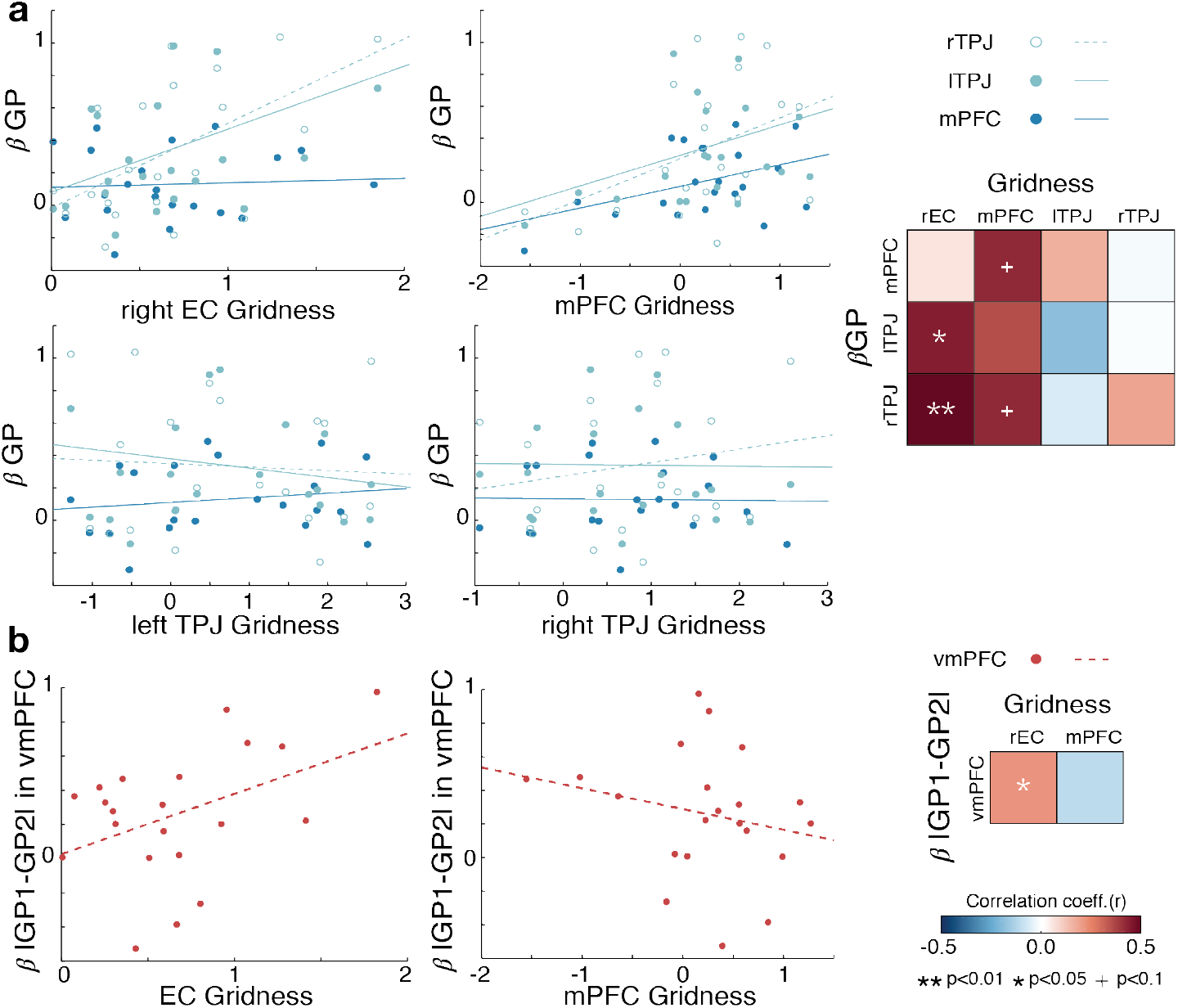
Individual differences in the relationship between the gridness and the effects of growth potential. **a.** In addition to the relationship between EC gridness *(β* cos(6[θ – Φ])) and GP effects (βGP) in bilateral TPJ that we present in **Fig. 4c** (upper left), we found a marginal positive correlation between the mPFC gridness and GP effects in mPFC and right TPJ (p<0.1) (upper middle). The gridness estimated in TPJ, however, does not correlate with their GP effects nor with the mPFC GP effect (p>0.1) (bottom left and middle). **, p<0.01; +, p<0.01. To further examine the relationship between the EC and mPFC gridness, we formally test which one better explains the GP effects in TPJ and mPFC. To address this question, we inputted the z-scored gridness of EC and mPFC into the same GLM to predict the GP effects in TPJ and mPFC. We found that the GP effect in bilateral TPJ was better explained by the EC gridness than the mPFC gridness (regression coefficient β_EC_=0.24** > β_mPFC_=0.17* for right TPJ; β_EC_=0.16* > β_mPFC_=0.13 for left TPJ; **, p<0.01 and *, p<0.05), and the GP effect in mPFC was better explained by the mPFC gridness than the EC gridness (β_mPFC_=0.09 (p=0.066) > β_EC_=0.01). Right: Colormap in matrix depicts regression coefficients for each regions’ gridness effect used to explain each regions’ GP effect. **b.** Left: Positive correlation between effects of differences in GP (IGP1-GP2I) in vmPFC during partner selection decisions and the EC gridness (r=0.43, p=0.05) but not mPFC gridness (r=-0.22, p=0.33). Right: Colormap shows regression coefficients for rEC and vmPFC gridness effects used to predict the vmPFC value difference effect. **, p<0.01; *, p<0.05; +, p<0.1.

