## Supplementary Information for "Inferences on a Multidimensional Social Hierarchy Use a Grid-like Code"

This PDF file includes:

Figures S1 to S11
Tables S1 to S7

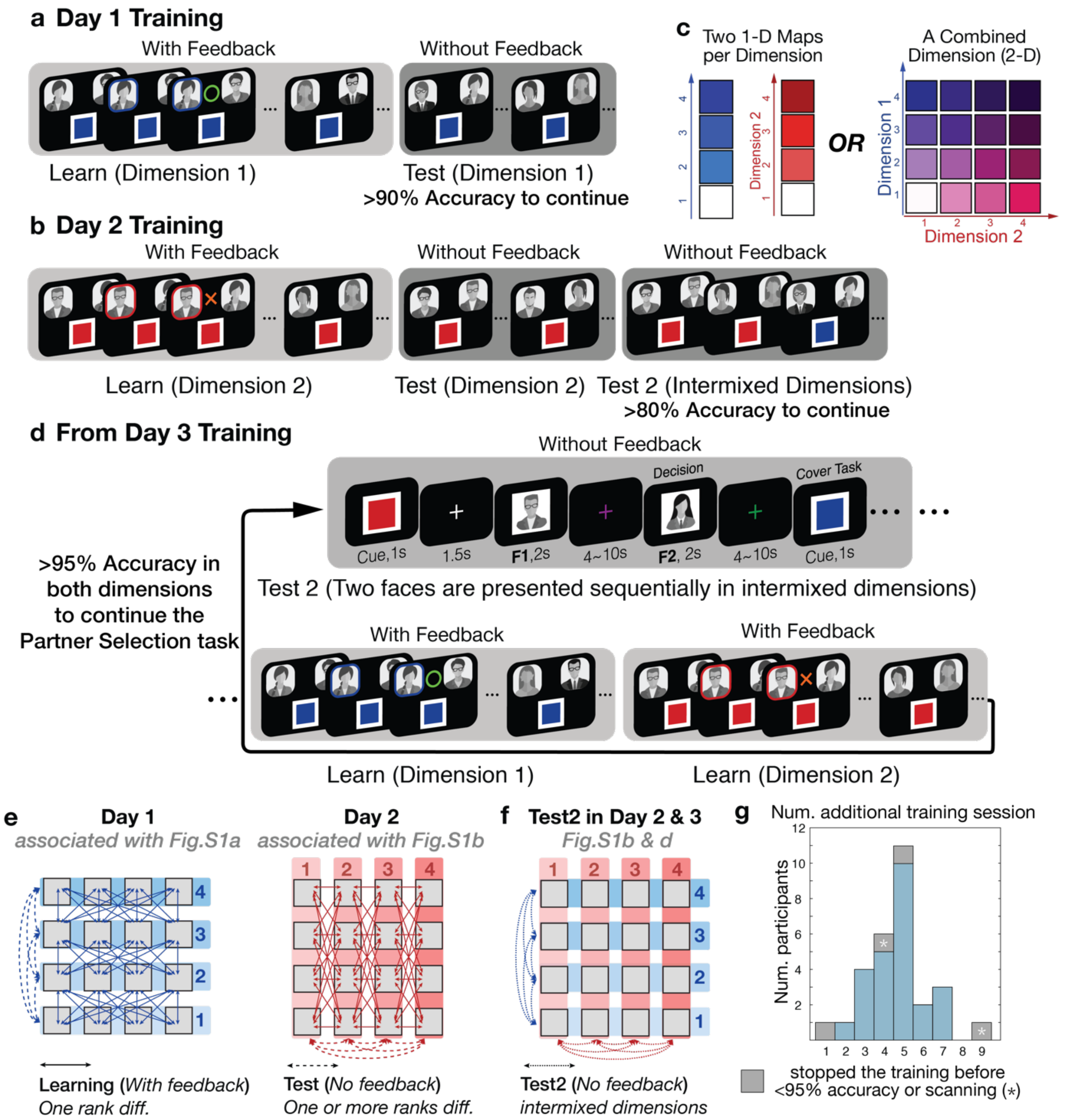


**Figure S1. Behavioral training before the fMRI partner selection task. a.** During the Learn phase on day 1, participants learned the relative rank of 16 individuals in one of two social hierarchy dimensions (indicated by cue color) based on feedback from binary comparisons. Participants were asked to choose the higher rank individual between two who differed by one level only in the given social hierarchy dimension. During the test phase on day 1, participants were asked to infer the relationship of people who were never paired during the Learn phase through transitive inferences. No feedback was given during the test phase. **b.** During the Learn phase on day 2, participants learned the relative status of the same 16 individuals in the unlearned dimension by comparing people who differed by one level only in the corresponding dimension. During the Test phase on day 2, participants were asked to infer the relative status of unlearned pairs through transitive inference. As before, no feedback was given during the test phase. At the end of day 2 training, participants' knowledge about both 1-D social hierarchies was tested (test

2). During the Test 2 phase, participants were asked to infer the relative status of two individuals while both dimensions were intermixed across trials. **c.** After day 1 training, participants could have built a hierarchical structure in one dimension. After training on day 2, participants could build two structures of social hierarchy per each dimension (left panel), or they could in principle have built a combined hierarchical structure in two dimensions (Right panel). **d.** The Test 2 phase and the two Learn phases per dimension were repeated until participants could make inferences correctly during the Test 2 phase with an accuracy higher than 90% in both dimensions to continue to the partner selection task. From day 3 training, during the Test 2 phase, stimuli were presented sequentially (top). While presenting F2, participants were asked to make an inference of the relative status of two individuals and indicate their decision. For the Test 2 phase, both dimensions were intermixed across trials and feedback was not given for participants' decisions. **e.** The squares indicate the position of individuals in the 4x4 social hierarchy. The arrows between individuals indicates the pairs that were presented to participants in learning block trials in which participants got feedback on their decision. As shown here, the paired individuals' ranks differ by 1 level in either the competence (left) or popularity (right) dimensions. Participants learned the relative status of those pairs in the left panel in the competence dimension (**Fig.S1a**) and the relative status of those pairs in the right panel in the popularity dimension (**Fig.S1b**) on different days. A learning block was followed by a test block. During the test block, all possible pairs except for oneself and those who are at the same rank in the given dimension were presented to participants. The dotted lines indicate the pairs presented during the test blocks that include pairs whose ranks differed by 1,2, and 3 ranks. **f.** After two days of training, Test2 blocks followed at the end of Day2 (the right panel in **Fig.S1b**) and at the beginning of Day3 (the top panel in **Fig.S1d**). The dotted lines indicate the pairs presented in Test2 block. The pairs presented in Test2 block comprise the pairs presented during the test blocks in both dimensions. **g.** The bar graph shows the number of additional sessions that each of 29 subjects participated in. 2 among 29 participants stopped the training before reaching 95% accuracy, and 2 who reached >95% stopped the experiment before scanning (marked with *).

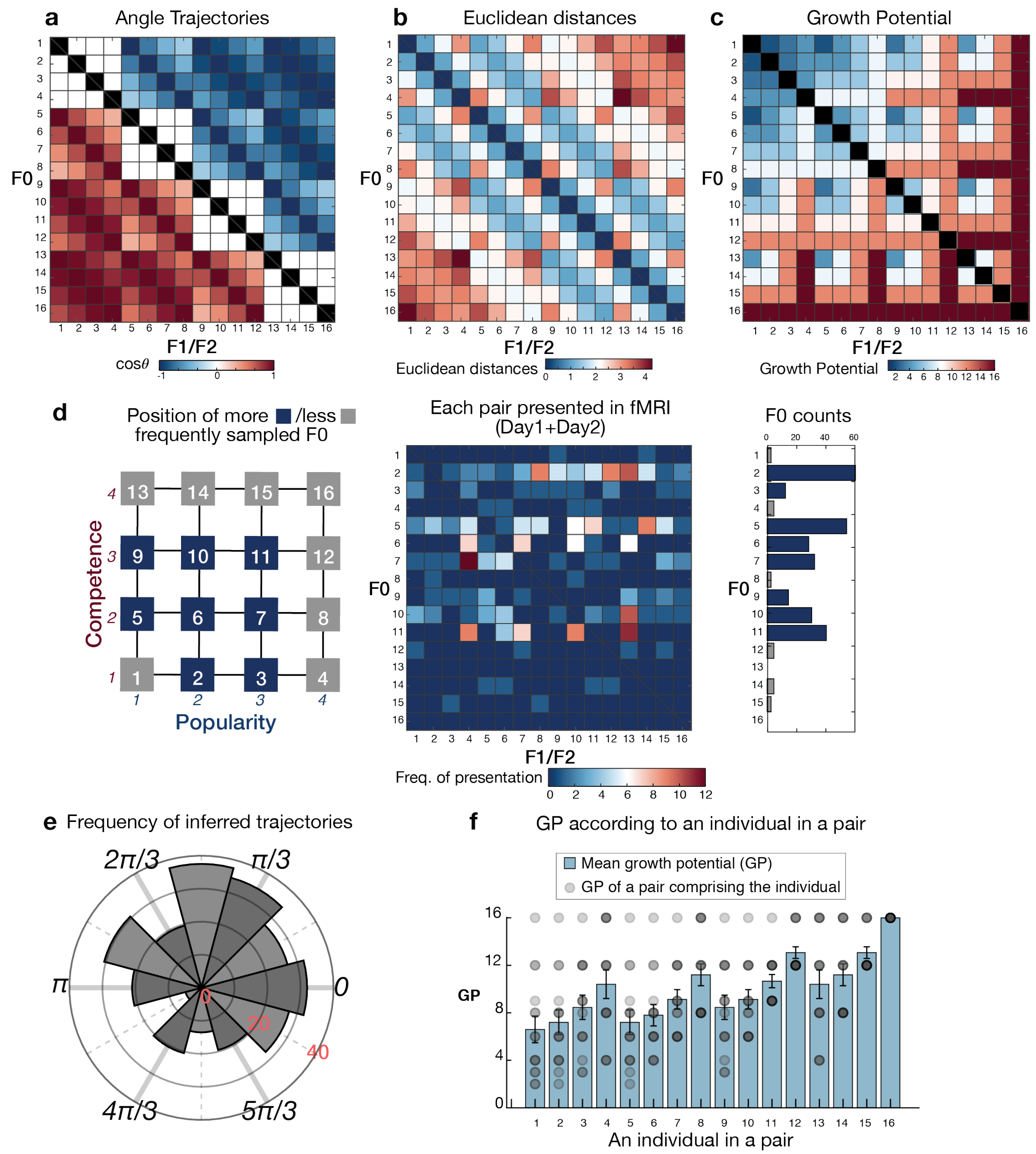


**Figure S2. The angles, distances, and GP of pairs during the partner selection task. a.** The 16 ×16 matrix which includes all possible pairs of F0 and F1/F2 shows the values of the cosine angle ($\cos\theta$) of the trajectories of all possible 240 pairs (except for the pairs in black where F0 and F1/F2 are the same). **b.** Euclidean distance of the trajectories of all possible pairs. **c.** Growth potential (GP) of the trajectories of all possible pairs. The pairwise Euclidean distance and the GP differ from each other, suggesting participants cannot use the Euclidean distance but should compute the GP of each pair to be accurate in the partner selection task. **d.** Among the 240 possible pairs shown in **Fig.S2a**, we carefully chose specific pairs for the partner selection task. The less sampled F0 positions are shown in gray (left panel). For the pairs where F0 is at the highest rank in one social hierarchy dimension, participants could make partner selection decisions by comparing the ranks in one dimension only. Similarly, position 1 and 16 always lose and win, respectively. Because we hypothesized the grid code may only be utilized when subjects have to integrate both dimensions and simulate a relationship in 2D, rather than rely on simple choice heuristics, we only minimally sampled those F0 positions. The middle panel shows the number of presentations of each pair during the fMRI partner selection task, and the right panel shows the number of pairs according to their F0 face. **e.** The frequency of decision trajectories in the experiment, categorized into 12 equal bins of 30° according to the direction of inferred trajectories, $\theta$. **f.** The dots indicate the possible GP an individual can have according to whom they are paired with. The bars indicate the mean GP±SEM of possible GP values for that individual. It shows that the GP cannot be predicted by only one individual in a pair, but instead participants need to take both individuals in a pair into account to compute the GP. Moreover, the mean GP cannot be well accounted for by a linear function of the rank in one dimension but by the ranks in both dimensions. This implies that participants cannot utilize a separate 1-D cognitive map of the GP to make accurate decisions, but need to use the ranks of both individuals in a pair in both social hierarchy dimensions.

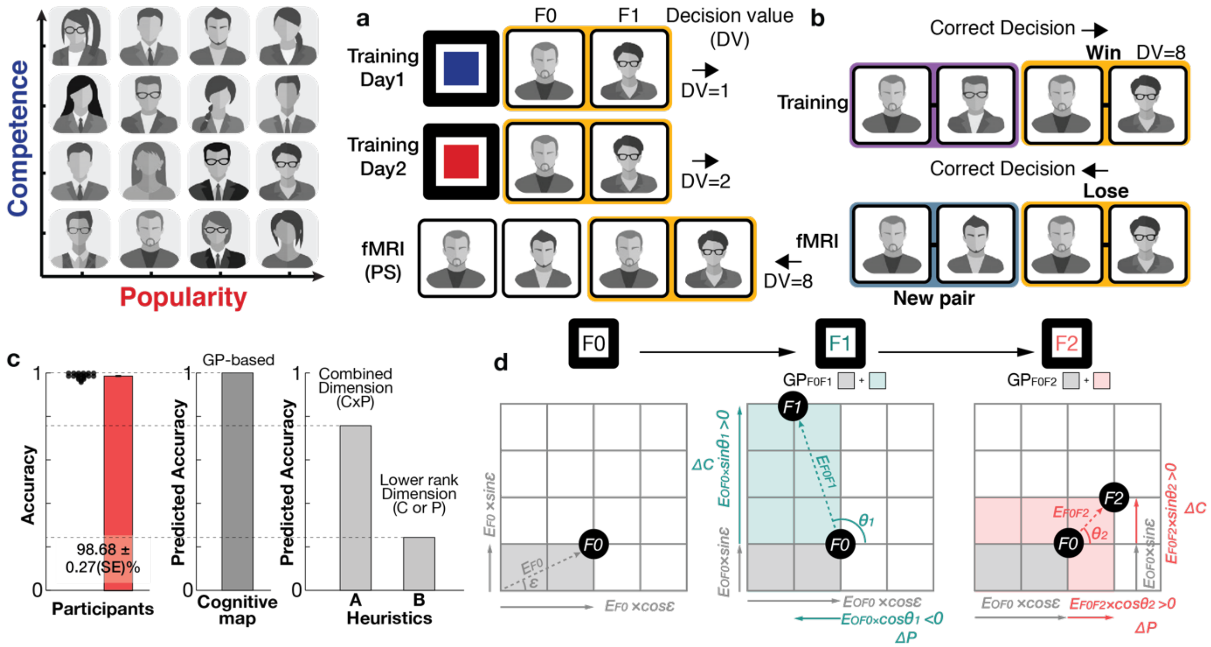


**Figure S3. Decision values in behavioral training cannot be generalized for partner selection decisions. a.** An example showing that when the same F0F1 pair is presented, the decision value (DV) in a trial during behavioral training (DV=1 for day 1 training; DV=2 for day 2 training) cannot be generalized for the DV for the partner selection decision (DV=8). **b.** An example showing that remembering the winning pair in a trial of the partner selection task during behavioral training (F0F1 pair here) does not help participants to make a correct decision in the partner selection task in fMRI since they are compared with other pairs that participants had never compared them with. These examples (**a** and **b**) illustrate that on most trials, participants needed to make inferences to compute the decision value (GP) in order to be accurate in the partner selection task, though they had been trained extensively to learn the ranks differences in each one-dimensional social hierarchy during behavioral training. **c.** Participants were able to choose the better partners during the partner selection task (mean accuracy ± sem = 98.68 ± 0.27%; left panel; a black dot indicates the accuracy of each participant). We examined whether the decisions were better explained by the difference between GPs of two pairs (GP1-GP2), which in principle may require participants to use the cognitive map (middle panel), or other alternative heuristics (right panel). As the first alternative way to make a decision, we tested whether participants used a rank assigned to each individual in a combined dimension (such as, rank in competence dimension (C) × rank in popularity dimension (P)), rather than using ranks of each of the two dimensions, and chose the one who had greater overall rank between F1 and F2 (Heuristic A). As the second alternative model, we tested whether participants only use the ranks of F1 and F2 in the dimension in which the rank of F0 was relatively deficient (Heuristic B). We found that both heuristics cannot explain the high accuracy that participants achieved during partner selection, suggesting that participants use the cognitive map to compute GP to guide correct decisions. **d.** The computation of growth potential (GP) based on the Euclidean distance and the angle of the inferred trajectories using the polar coordinate system. At the time of F0 presentation (left panel), the ranks of F0 in the competence and popularity dimensions are $E_{F0}\times\sin(\varepsilon)$ and $E_{F0}\times\cos(\varepsilon)$, respectively, where $E_{F0}$ indicates the Euclidean distance of $\overrightarrow{F0}$ vector from the origin, [0,0] and their angle is $\varepsilon$. At the time of F1 presentation, participants can compute the $GP_{F0F1}$ based on the non-negative relative rank difference between F1 and F0. The rank difference in the popularity dimension is computed as $\Delta P = E_{F0F1}\times\cos(\theta)$, while the rank difference in the competence dimension is computed as $\Delta C = E_{F0F1} \times \sin(\theta)$. Therefore, $GP_{F0F1}$ is computed as $(C0+\Delta C) \times (P0+\Delta P)$, while $\Delta C$ and $\Delta P \geq 0$. Last, $GP_{F0F2}$ is also computed in the same way when F2 is presented.

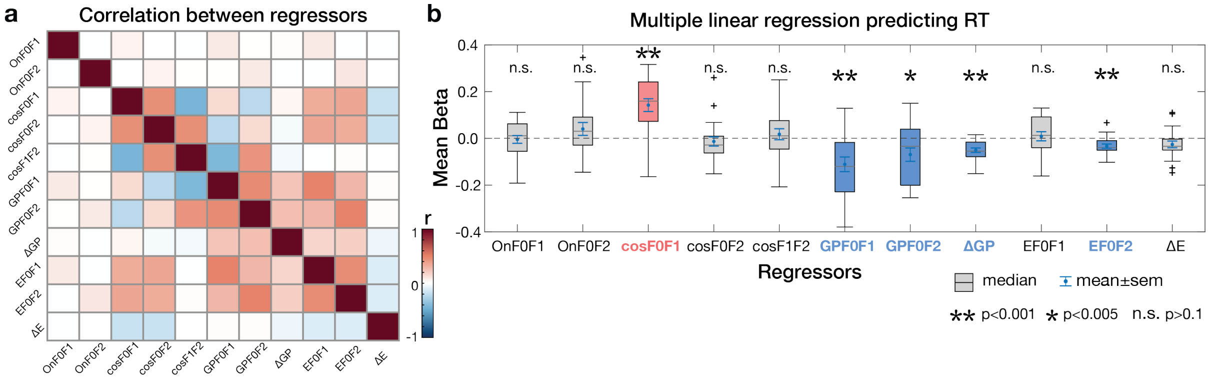


**Figure S4. Results of a multiple linear regression on reaction times (RTs) of partner selection decisions. a.** The correlation matrix shows the correlation (r) between 11 regressors. The 11 regressors include the Euclidean distance between F0F1 and F0F2 pairs, the absolute difference between them ($E_{F0F1}$, $E_{F0F2}$, and $\Delta E = |E_{F0F1} - E_{F0F2}|$), the GP of F0F1 and F0F2 pairs, and their absolute difference ($GP_{F0F1}$, $GP_{F0F2}$, and $\Delta GP = |GP_{F0F1} - GP_{F0F2}|$), the cosine angles of the trajectories ($cos\,\theta_{F0F1}$, $cos\,\theta_{F0F2}$, and $cos\,\theta_{F1F2}$) and whether the $\overrightarrow{F0F1}$ and $\overrightarrow{F0F2}$ vector was aligned to the EC grid orientation or not ($On_{F0F1}$ and $On_{F0F2}$). In this analysis, we compute the angle aligned to the diagonal since ranks in both dimensions are equally important in the partner selection task. Therefore, cos($\theta$-$\pi$/4) was inputted into the regressors. We found that the results changed only minimally when the cos angles were computed from the x-axis, cos($\theta$). **b.** The mean beta (regression coefficients). We found significant negative effects of the $E_{F0F2}$, $GP_{F0F1}$, $GP_{F0F2}$, $\Delta GP$ on RT (indicating faster RT) and a positive significant effect of $cos\,\theta_{F0F1}$on RT (indicating slower RT). The effect sizes of this regression are reported in **Table S5**. **p<0.001, *p<0.005, n.s. p>0.1. The distances of trajectory ($E$) have a moderate correlation level with the decision values ($GPs$) (r=0.47). When we include $E_{F0F1}$ and $E_{F0F2}$ after partialling out their covariance with $GP_{F0F1}$and $GP_{F0F2}$, the $GP$ effects ($GP_{F0F1}$and $GP_{F0F2}$) get a little stronger (t=-4.67 and t=-4.85 respectively p<0.005) while no notable changes were found for effects of the other regressors.

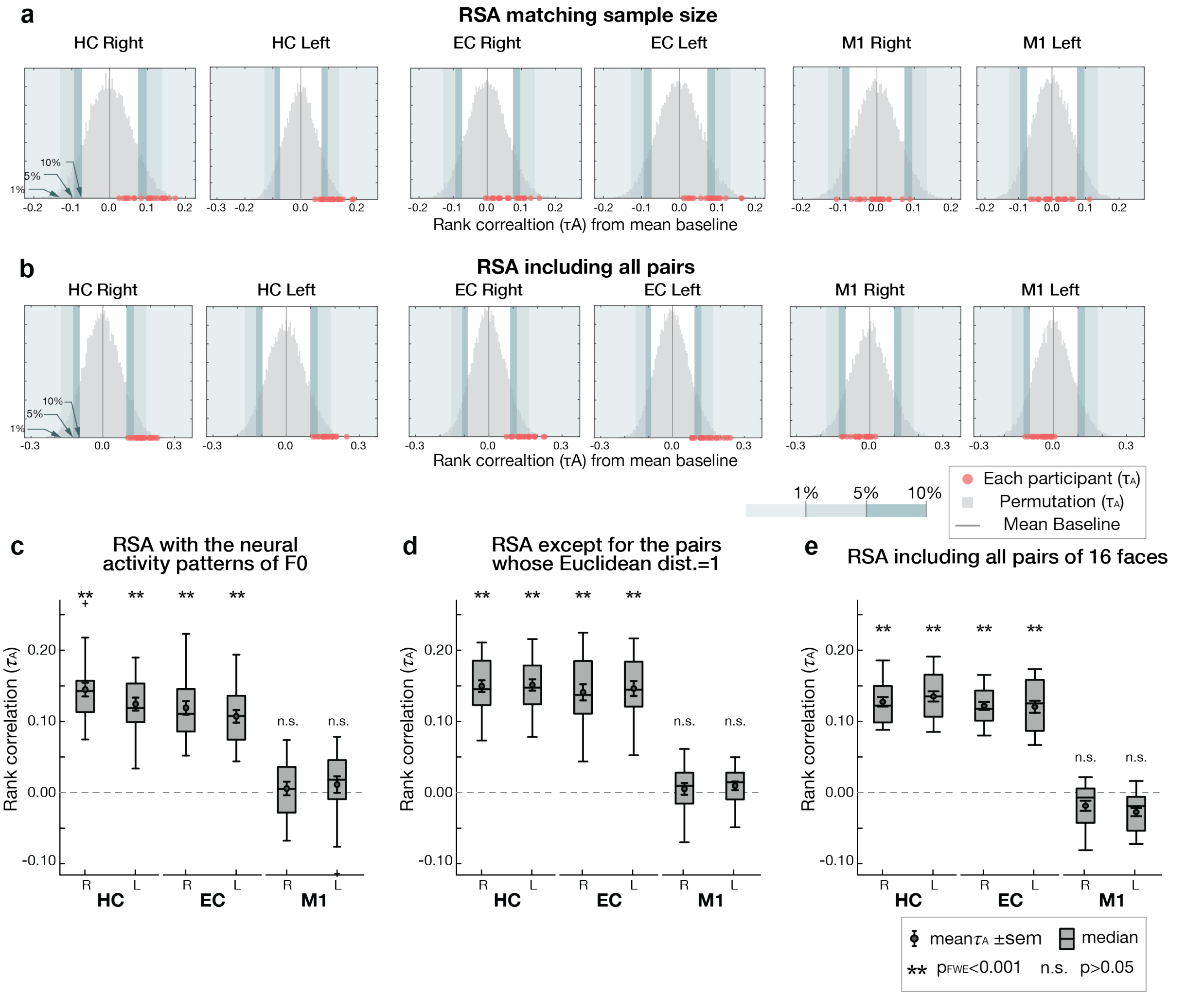


**Figure S5. Representational similarity analysis (RSA). a and b.** The rank correlation (Kendall's $\tau A$) of each of the participants corrected from the baseline (n=21) are shown as red dots on the x-axis. $\tau A$ indicates to what extent the pairwise Euclidean distance in the 2-D social hierarchy explains the neural activity patterns of the ROIs. The gray histogram shows the baseline distribution of $\tau A$ which was acquired from 1000 permutations while randomly shuffling the positions of the 14 individuals in the 2-D space. Based on the distribution, the thresholds of p=0.01, p=0.05, and p=0.1 are marked for two-tailed tests. The $\tau A$ of most participants are within the range of 0~10% in the bilateral HC and EC, whereas most are within the range of 10~90% in bilateral M1. We performed RSA while down-sampling the observations to match the same sample size of each face presentation to be the same (**a**) and while including all observed samples (**b**). **c.** To examine activity patterns in our ROIs while the activity was minimally modulated by other task-relevant cognitive processes, we performed a control RSA in which the neural activity only acquired at the time of F0 presentation (but not F1 or F2 presentation) was included. Since some faces were selected less frequently as F0 during partner selection, we only include 8 individuals at the positions in the social hierarchy for the control RSA. The dissimilarity between activity patterns associated with F0 presentation estimated in bilateral HC and EC increases in proportion to the pairwise Euclidean distance between individuals in the 2-D abstract social space. The rank correlation (Kendall's $\tau_A$) shows robust effects of Euclidean distance on the pattern dissimilarity estimated in the HC and EC compared to the permuted baseline (1000 iterations), but not in our control region (M1) **d.** After excluding the pairs with Euclidean distance of 1, we still found robust effects of Euclidean distance on the increases in the pattern dissimilarity estimated in the HC and EC compared to the permuted baseline (1000 iterations), but not in a control region (M1). **e.** After including all the pairs between 16 individuals, we found robust effects of Euclidean distance on the increases in the pattern dissimilarity estimated in the HC and EC compared to the permuted baseline (1000 iterations) but not in M1. **, $p_{FWE}<0.001$ Bonferroni-Holm method; n.s., $p>0.05$, uncorrected. **c. d., and e.** These results are consistent with the finding shown in **Fig.2** and **Extended Data Figure 1** using faces at all events (F0, F1 and F2 presentations).

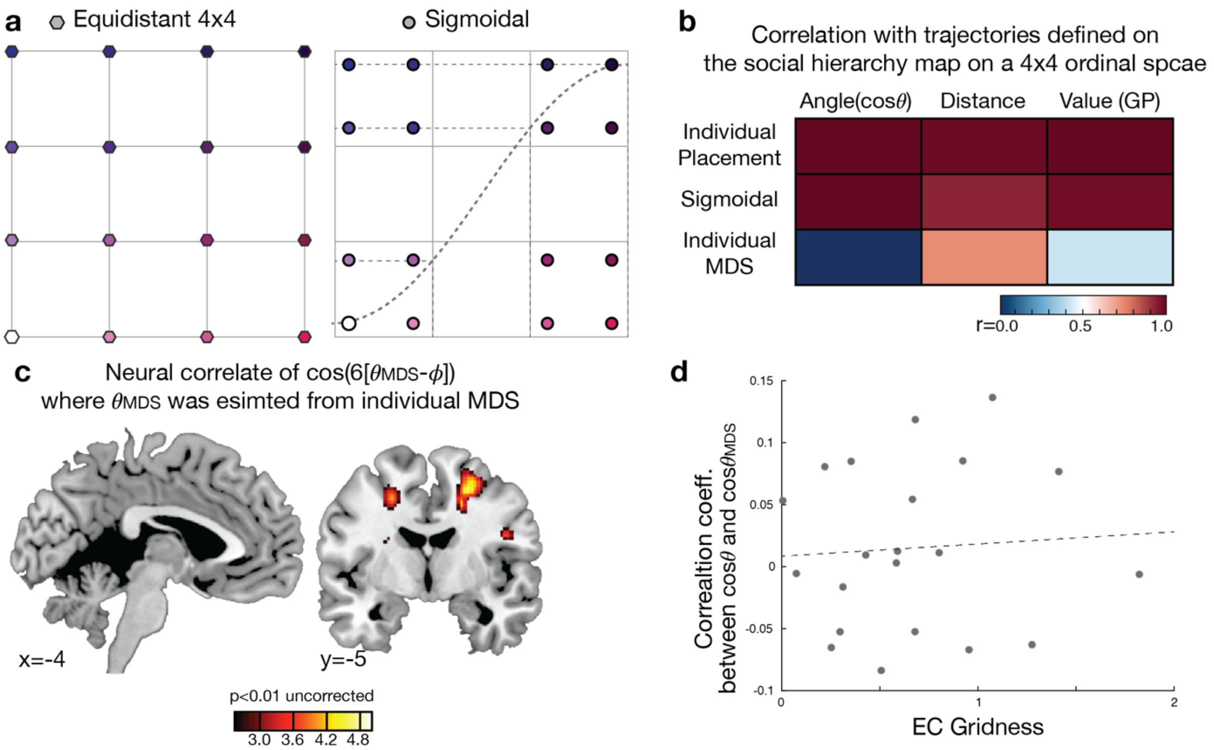


**Figure S6. Inferred trajectories defined on cognitive maps based on alternative geometries. a.** The positions of 16 individuals computed using non-linear monotonic dimensions using a sigmoid function to test the effects of deformation of the cognitive map. **b.** The level of correlation between the properties of inferred trajectories (angles, distances, and decision value [GP]) computed in a 4×4 ordinal space and those computed from other cognitive maps defined on alternative dimensions (placement, sigmoidal and MDS estimated from the hippocampus (HC) of individual participants). **c.** The effects of hexadirectional grid-like coding modulated by the angles of trajectories estimated from the coordinates of each individual MDS ($\theta_{MDS}$), rather than angles in the true hierarchy space, were tested. We found only the motor cortex showed significant effects at a reduced threshold ($p<0.005$, uncorrected) when estimating $\theta$ from the individual MDS. **d.** In addition to the within-subject analyses, we further performed a between-subject analysis to test whether those participants who had greater grid effects in the EC were more likely to have an MDS close to the true social hierarchy structure. The EC gridness indicates to what extent the EC activity of each participant is hexagonally modulated. The level of similarity between the MDS of each participant and the true structure was estimated from the correlation coefficient between cosine angle trajectories estimated from each of two spaces. We find that the EC gridness does not correlate with to what extent the MDS closely reflects the true social hierarchy structure ($r=0.07$).

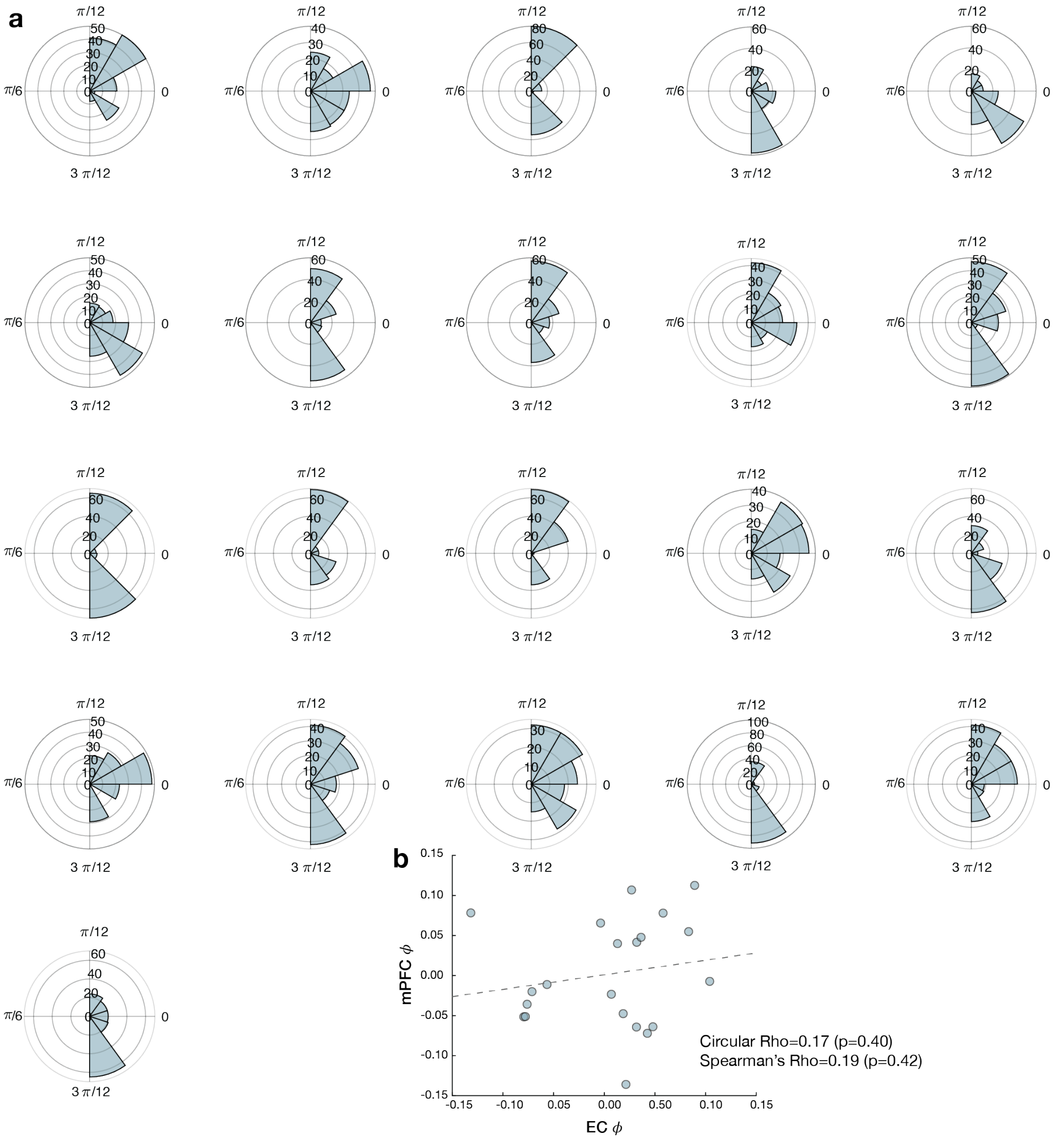


**Figure S7. Nonuniform distributions of EC grid orientation. a**. Clustering of putative EC grid orientations within each participant (n=21). Polar histograms show potential grid orientations (in the range 0 and $\pi/3$) of each participant estimated from the sessions acquired for the day1 scan for all voxels in the EC ROI. Grid orientations were significantly clustered in all participants ($p<0.01$, Rayleigh's tests for nonuniformity; mean z±SEM=50.98±4.14 for the first day scan and 58.66±4.23 for the second day scan). **b.** While the EC grid orientations (EC $\phi$) correlate with the mPFC grid orientations (mPFC $\phi$) minimally across participants ($p>0.05$), we found that 80.19±1.66% (SEM) of total pairs were classified as the same category (aligned or misaligned) when aligned to the EC and when aligned to the mPFC grid orientations.

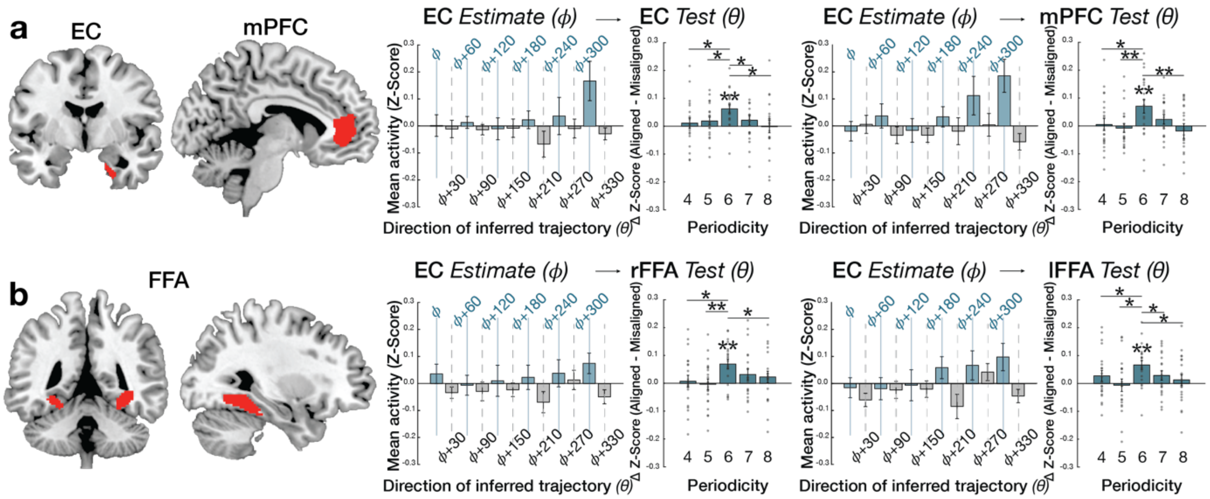


**Figure S8. Hexadirectional modulation on anatomically defined ROIs. a.** Confirmatory analyses of hexadirectional modulation on independent anatomically defined EC (Amunts *et al.*, 2005; Zilles and Amunts, 2010) and mPFC (Neubert *et al.*, 2015) ROIs in alignment with the EC grid angle. Consistent with our functionally defined ROI-based analyses (**Fig.3b,c)**, this effect in anatomically defined EC and mPFC was also specific to six-fold periodicity (one-sided t-test, $p<0.01$), as it was not seen not seen for four-, five-, seven-, or eightfold periodicities (all $p>0.05$). The effect at six-fold was significantly greater than those of the control periodicities (paired t-test). **b.** Hexadirectional modulation was also tested on independent, functionally-defined FFA ROIs. The FFA ROIs were identified by a contrast analysis between presentations of face stimuli and the fixation cross (within a mask defined at the threshold $p<0.001$, uncorrected). We found that this effect in bilateral FFA was also specific to a six-fold periodicity ($p<0.01$), as it was not present for other control periodicities (all $p>0.05$, uncorrected). **, $p<0.01$ and *, $p<0.05$. The effect at six-fold was significantly greater than those of the control periodicities for all but one comparison in rFFA (paired t-test).

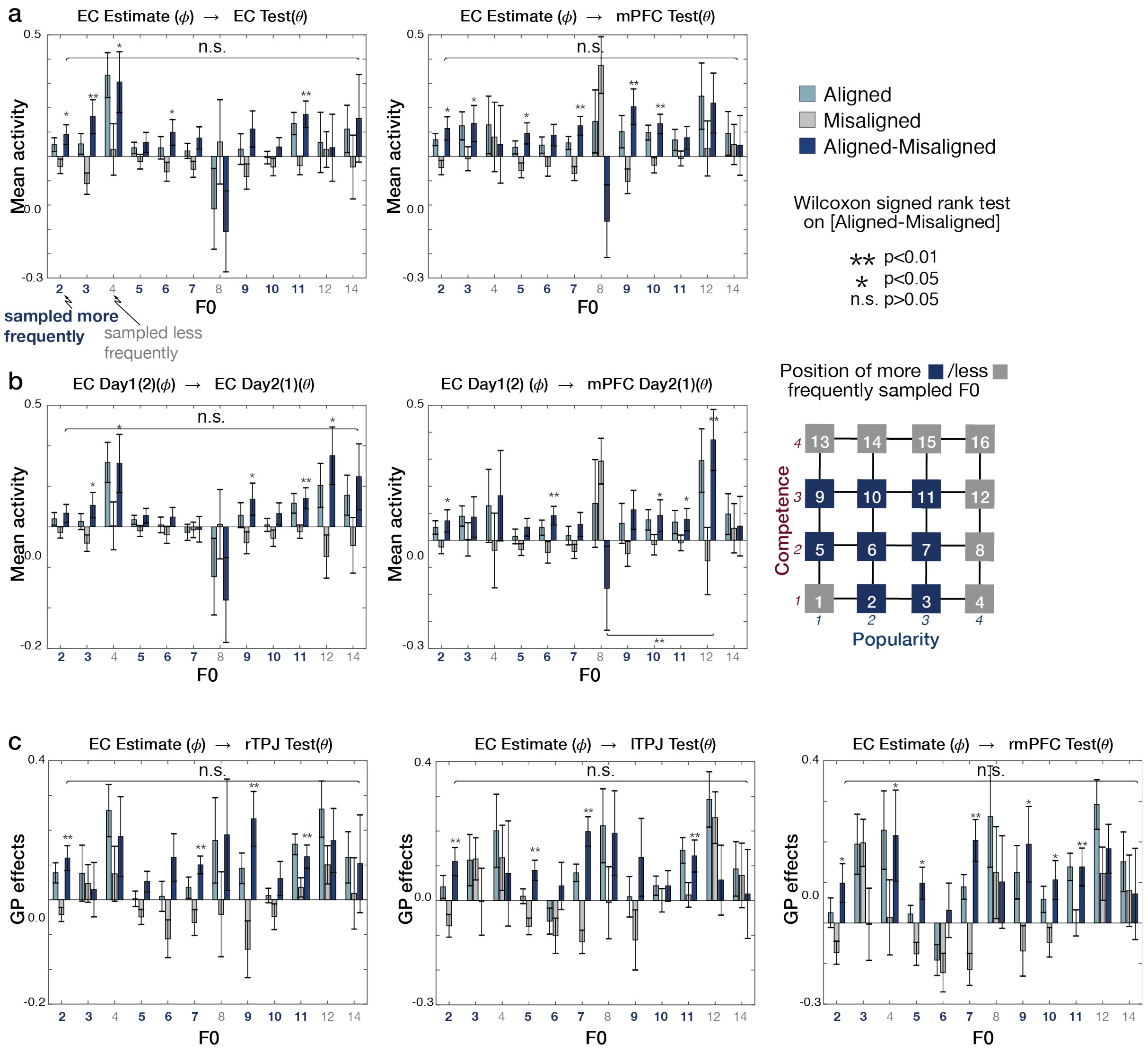


**Figure S9. Grid effects of the inferred trajectories were tested separately according to F0 of the pairs. a.** The mean activity in the EC and mPFC when the trajectories are aligned and misaligned to the EC grid orientation estimated from different blocks of the same day scan (Associated with **Fig.3b**). The blue bars indicate their differences (aligned - misaligned). **<0.01, *<0.05 in Wilcoxon signed rank test. This effect was not different across pairs according to the F0 position (p>0.05 in Kruskal-Wallis test). Note that the statistical analysis is not always reliable for some F0 positions since the sample sizes are significantly reduced for this analysis. **b.** The mean activity in the EC and mPFC when the trajectories are aligned and misaligned to the EC grid orientation estimated from the scan acquired from a different day (associated with **Fig.3c**). This effect was not different across pairs according to the F0 position in EC (p>0.05 in Kruskal-Wallis test). In mPFC, the effects did not differ across the F0 positions (p>0.05) except for the pair between face8 and face12 (p<0.01), although this difference is likely unreliable given their small number of samples. **c.** The GP effects ($\beta$GP) in the mPFC and bilateral TPJ when the trajectories are aligned and misaligned to the EC grid orientation (associated with **Fig.4b**). This effect was not different across pairs according to the F0 position (p>0.05 in Kruskal-Wallis test).

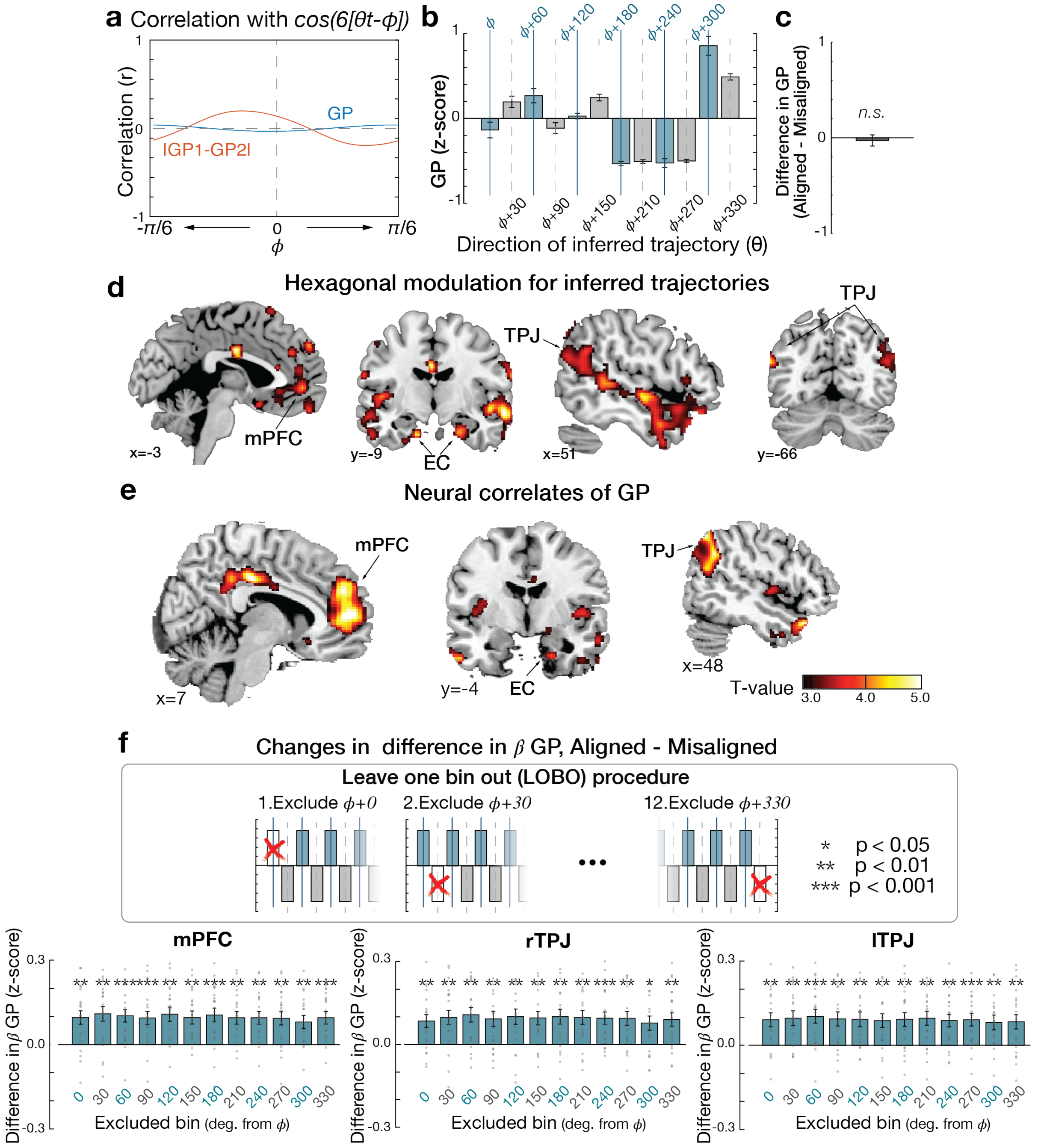


**Figure S10. Grid-like coding while controlling for the growth potential of inferred trajectories. a.** To make optimal decisions in the partner selection task, we hypothesized that participants could make inferences along direct trajectories between two individuals which guide the computation of GP and subsequently the two option's GP comparison, lGP1-GP2l. We confirmed that both GP and lGP1-GP2l were not explained by a function of the direction of the inferred trajectory, $cos(6[\theta - \phi])$ regardless of the grid orientation, $\phi$. This low correlation indicates that it is statistically possible to identify the neural encoding of GP while controlling for the neural modulation of the direction of inferred trajectories. **b.** The z-scored GP within each block categorized into 12 equally distributed bins of 30° according to the direction of the inferred trajectories ($\theta$), aligned to each participant's EC grid orientation ($\phi$) shows that the distribution of GP did not show the hexagonally symmetrical pattern. This pattern shows that the hexadirectional modulations of $\beta$ GP in TPJ and mPFC (**Fig.4b**) were not caused by difference in GP itself but were specific for the inferred trajectories in alignment with the EC grid orientation, **c.** We further tested whether there was a significant difference in GP between aligned and misaligned trajectories, which was not present (one-sided t test; t=-0.34, p=0.63). **d and e.** We found that the effects of hexagonal modulation $(cos(6[\theta_t - \phi]))$ (**Fig.3a**) and the GP (**Fig.4a**) remained when including both regressors in the same GLM, indicating each of the regressors explained independent variance in these regions. **f.** Leave-one-bin-out (LOBO) test. The average differences in z-scored GP effects (±SE) across participants between aligned and misaligned trajectories. The positive z-score differences (aligned > misaligned) suggesting hexadiretional modulations in the activity encoding GP in mPFC and bilateral TPJ were found when they were computed based on the activity of 11 bins while excluding the activity in one of 12 bins (all p<0.05). This result shows that the effects of hexadirectional modulation were not driven by the activity in any single direction but prevalent across directions. **, p<0.01; **, p<0.01.

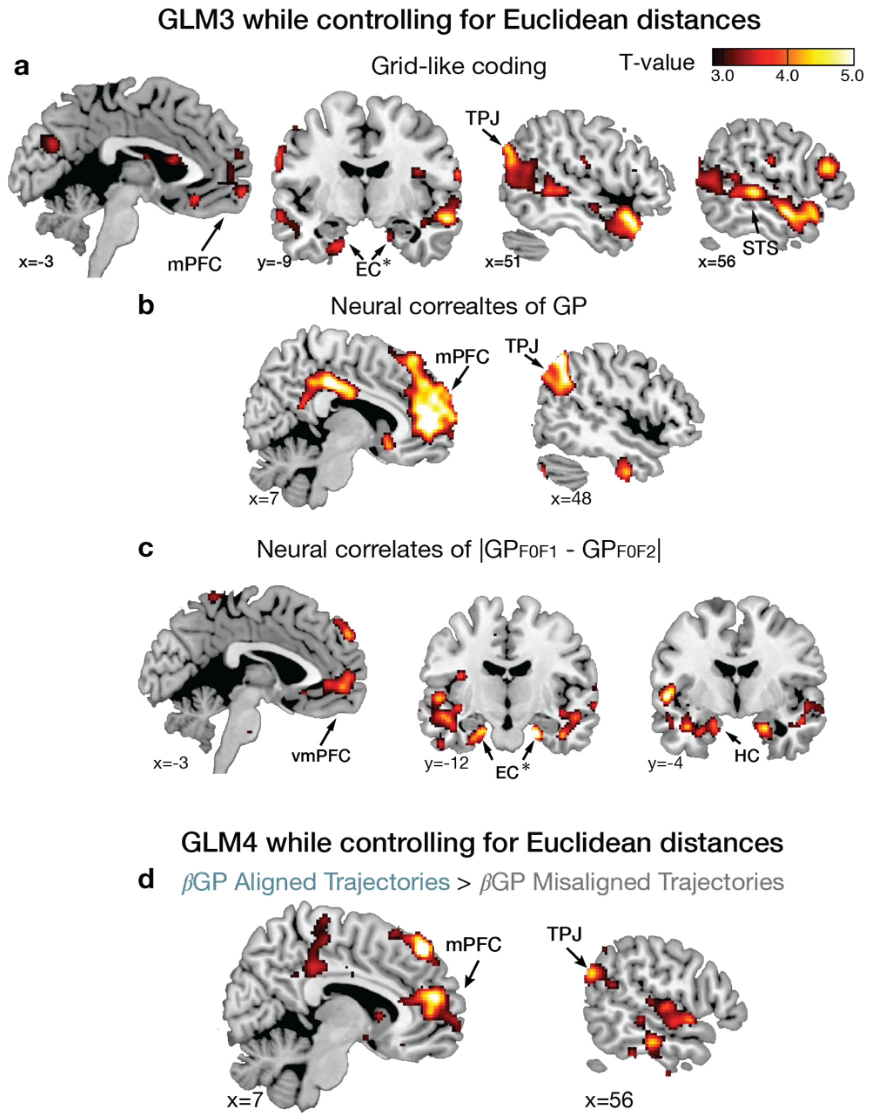


**Figure S11. Grid-like coding and GP effects while controlling for the Euclidean distances of inferred trajectories.** Associated with **Fig.4**. **a.** The hexadirectional grid-like effect ($cos(6[\theta - \phi])$) is shown, while controlling for the Euclidean distance of trajectories ($E_{F0F1}$ and. $E_{F0F2}$) (GLM3). The effects of grid-like coding were significant in mPFC, TPJ, and STS ($p_{FWE}$<0.05 whole brain TFCE correction) and EC ($p_{FWE}$<0.05 corrected in the anatomically defined EC ROI [marked with ✻]). **b.** Whole-brain map showing effects of the neural correlates of growth potential (GP), while controlling for the Euclidean distance of trajectories ($E_{F0F1}$ and. $E_{F0F2}$). The neural correlates of GP were significant in mPFC and TPJ ($p_{FWE}$<0.05 whole brain TFCE correction). **c.** Whole-brain map showing effects of value comparison ($|GP_{F0F1}$-$GP_{F0F2}|$). The neural correlates of value comparison were significant in vmPFC and HC ($p_{FWE}$<0.05 whole brain TFCE correction) and EC ($p_{FWE}$<0.05 corrected in the anatomically defined EC ROI), while controlling for the Euclidean distance of trajectories ($E_{F0F1}$ and. $E_{F0F2}$). **d.** Whole-brain map contrasting the effects of GP (β GP) for trajectories aligned with the EC grid orientation, $\phi$, compared to those misaligned trajectories (β GP aligned > β GP misaligned) (GLM4). The contrast effects of GP were significant in mPFC, and TPJ ($p_{FWE}$<0.05 whole brain TFCE correction), while controlling for the Euclidean distance of trajectories ($E_{F0F1}$ and. $E_{F0F2}$). For visualization purposes, the whole-brain maps are thresholded at p<0.005 uncorrected.

| Brain areas | Laterality | k | Z | Peak coordinate (MNI) | | |
|---|---|---|---|---|---|---|
| | | | | x | y | z |
| Medial prefrontal cortex (mPFC) | R/L | 768 | 5.58 | 2 | 66 | -4 |
| Posterior cingulate cortex (PCC) /Precuneus | R/L | 181 | 3.19 | 2 | -50 | 36 |
| Posterior parietal cortex (PPC) | R | 149 | 4.27 | 36 | -46 | 62 |
| Posterior parietal cortex (PPC) | L | 289 | 3.81 | -38 | -50 | 50 |
| Lateral orbitofrontal cortex (lOFC) | L | 65 | 3.79 | -42 | 44 | -8 |
| Retrosplenial cortex (RSC) | R/L | 49 | 3.45 | 2 | -54 | 28 |
| Entorhinal cortex (EC)✻ | R | 21 | 2.80 | 26 | -10 | -40 |

**Table S1. Whole brain analysis showing hexagonally symmetric signals.** All $p_{TFCE}<0.05$ within whole brain Threshold-Free Cluster Enhancement (TFCE) correction (Smith and Nichols, 2009) except with ✻ which indicates the correction within *a priori* regions of interest (ROI). The ROI was anatomically defined in the EC (Amunts *et al.*, 2005; Zilles and Amunts, 2010). Cluster size (k) was reported at Z>3.1 which corresponded to p<0.001 for all regions except for the *a priori* hypothesized effect in EC, in which we reported the cluster size at the threshold, Z>2.3 (p<0.01). R/L: the cluster extended across bilateral hemispheres. R, right hemisphere; L, left hemisphere.

**a. Hexadirectioanl modulations aligned to EC grid orientation**
(cross-validation (CV) between sessions acquired within the same day)

| Brain areas | Laterality | k | T | Peak coordinate (MNI) | | |
|---|---|---|---|---|---|---|
| | | | | x | y | z |
| Medial prefrontal cortex (mPFC) | R/L | 114 | 4.72 | -6 | 48 | -4 |
| Temporoparietal junction (TPJ) | R | 238 | 3.67 | 46 | -58 | 20 |
| Temporoparietal junction (TPJ) | L | 39 | 5.71 | -56 | -68 | 24 |
| Superior temporal sulcus (STS) | R | 794 | 4.05 | 50 | -40 | 4 |
| Superior temporal sulcus (STS) | L | 251 | 4.29 | -60 | -24 | -6 |
| Entorhinal cortex (EC) | R | 75 | 4.11 | 22 | -10 | -28 |

**b. Hexadirectional modulations aligned to EC grid orientation**
(CV between sessions acquired from a different day after more than a week)

| Brain areas | Laterality | k | T | Peak coordinate (MNI) | | |
|---|---|---|---|---|---|---|
| | | | | x | y | z |
| Medial prefrontal cortex (mPFC) | R/L | 278 | 5.04 | -2 | 36 | -8 |
| Posterior cingulate cortex (PCC) | R/L | 36 | 3.68 | 6 | -58 | 30 |
| Temporoparietal junction (TPJ) | L | 480 | 4.06 | -36 | -64 | 22 |
| Inferior temporal cortex (STS) | L | 106 | 4.20 | -52 | -64 | -4 |
| Fusiform gyrus (FFA) | R | 32 | 3.82 | 44 | -40 | -16 |
| Fusiform gyrus (FFA) | L | 106 | 4.00 | -48 | -54 | -6 |
| Entorhinal cortex (EC) ✻ | R | 30 | 4.21 | 36 | -10 | -38 |

**Table S2. Brain areas showing hexadirectioanl grid-like coding. a.** In association with **Fig.3a**. Whole brain analysis showing hexadirectioanl modulations according to the direction of inferred trajectories aligned with the EC grid orientation consistently across sessions acquired within the same day. **b.** In association with **Extended Data Figure 3d**. Whole brain analysis showing hexadirectioanl modulations according to the direction of inferred trajectories aligned with the EC grid orientation consistently across sessions acquired from a different day after more than a week. All $p_{TFCE}$<0.05, whole-brain cluster corrected using TFCE (Smith and Nichols, 2009) except with ✻ which indicates the correction within *a priori* regions of interest (ROI). The ROI was anatomically defined in the EC (Amunts *et al.*, 2005; Zilles and Amunts, 2010).

**a. Neural correlates of GP**

| Brain areas | Laterality | k | T | Peak coordinate (MNI) x | y | z |
|---|---|---|---|---|---|---|
| Medial prefrontal cortex (mPFC) | R/L | 1460 | 5.55 | 10 | 52 | 6 |
| Temporoparietal junction (TPJ) | R | 870 | 5.45 | 54 | -56 | 34 |
| Temporoparietal junction (TPJ) | L | 276 | 5.18 | -54 | -60 | 28 |
| Posterior cingulate cortex (PCC) | R/L | 246 | 5.76 | 0 | -24 | 36 |
| Entorhinal cortex (EC)✻ | R | 10 | 3.88 | 20 | -4 | -32 |

**b. Neural correlates of lGP1-GP2l**

| Brain areas | Laterality | k | T | Peak coordinate (MNI) x | y | z |
|---|---|---|---|---|---|---|
| Ventromedial prefrontal cortex (vmPFC) | R/L | 484 | 5.07 | -8 | 54 | -6 |
| Supramarginal cortex (SMC) | L | 1723 | 5.68 | -62 | -62 | 8 |
| Fusiform face area (FFA) | L | 14 | 4.10 | -32 | -30 | -24 |
| Hippocampus (HC)/ Entorhinal cortex (EC) | L | 79 | 4.53 | -28 | -4 | -26 |
| Entorhinal cortex (EC)✻ | R | 140 | 5.10 | 22 | -12 | -26 |
| Entorhinal cortex (EC)✻ | L | 14 | 4.00 | -26 | -16 | -32 |

**c. Contrasts of neural correlates of GP between aligned and misaligned trajectories**

| Brain areas | Laterality | k | T | Peak coordinate (MNI) x | y | z |
|---|---|---|---|---|---|---|
| Medial prefrontal cortex (mPFC) | R/L | 398 | 4.97 | 8 | 46 | 8 |
| Medial frontal gyrus (mFG) | R | 280 | 5.28 | 8 | 42 | 50 |
| Medial frontal gyrus (mFG) | L | 27 | 4.16 | -20 | 36 | 50 |
| Inferior orbitofrontal cortex (iOFC)/ Anterior insula | R | 339 | 5.83 | 30 | 12 | -22 |
| Temporoparietal junction (TPJ) | R | 78 | 4.92 | 56 | -66 | 28 |

**Table S3. Neural encoding of decision variables. a.** Whole brain analysis showing neural correlates of decision variables including growth potential (GP, in association with **Fig.4a**). **b.** Neural correlates of differences between GPs (lGP1-GP2l, in association with **Fig.4f**). **c.** The contrast map of neural correlates of GP between aligned and misaligned trajectories (Aligned > Misaligned, in association with **Fig.4d**). ✻ indicates correction within *a priori* regions of interest (ROIs). The ROI was anatomically defined in the EC (Amunts *et al.*, 2005; Zilles and Amunts, 2010). All other brain areas were significant at the threshold, $p_{TFCE}<0.05$, whole-brain cluster corrected using TFCE (Smith and Nichols, 2009). Cluster size (k) was reported at $p<0.001$.

| | ROIs | One-sample t test (First 6 30° bins) $\phi \pm 15° \sim [\phi + \frac{5\pi}{6}] \pm 15°$ | | Paired t test (Novel pairs only) (First 6 *vs.* last 6 30° bins) | |
|---|---|---|---|---|---|
| | | T | p | T | p |
| CV across blocks (Associated with **Fig.3b**) | EC xB | 2.35 | 0.03* | -0.96 | 0.35 |
| | mPFC xB | 3.84 | 0.00** | -1.48 | 0.16 |
| CV across days (Associated with **Fig.3c**) | EC xD | 2.35 | 0.03* | -0.72 | 0.48 |
| | mPFC xD | 2.72 | 0.01* | -0.33 | 0.75 |
| GP effects (Associated with **Fig.4b**) | mPFC GP | 5.07 | 0.00** | -0.20 | 0.85 |
| | rTPJ GP | 3.61 | 0.00** | -1.08 | 0.29 |
| | lTPJ GP | 4.98 | 0.00** | -1.06 | 0.30 |

**Table S4. Hexadirectional grid-like effects restricted to the low angle space.** The grid effects (aligned – misaligned) analysis including only the pairs in the first six 30° bins ($\phi \pm 15° \sim [\phi + 5\pi/6] \pm 15°$). The mean activity of aligned pairs and GP effects ($\beta$GP) of aligned pairs are greater than those of the misaligned pairs (One-sample t test; ** $p<0.01$, * $p<0.05$; left column in the Table). This finding shows that that the heaxadirectional grid-like effect was not only driven by angles larger than 180°. The more frequent samples of inferred trajectories could influence the relatively weaker effects in first six bins. We further test whether the effect sizes differ between the first and the last six bins when the pairs were presented for the first time. Specifically, we tested the grid effects (aligned – misaligned) including only the pairs that were shown for the first time to the participants (novel pairs only). We further compared the grid effects of the first six 30° bins (lower angle bins; $\phi \pm 15° \sim [\phi + 5\pi/6] \pm 15°$) to those of the last six 30° bins (higher angle bins; $[\phi + \pi] \pm 15° \sim [\phi + 11\pi/6] \pm 15°$). We find that neither the mean activity nor the GP effects ($\beta$GP) of aligned compared to misaligned pairs differ between the first and the last six bins when the pairs are shown first time to the participants (Paired t test, all $p>0.05$; right column).

| $On_{F0F1}$ | $On_{F0F2}$ | $\cos\theta_{F0F1}$ | $\cos\theta_{F0F2}$ | $\cos\theta_{F1F2}$ | |
|---|---|---|---|---|---|
| -0.295 | 1.496 | 5.335 ** | -0.693 | 0.773 | |
| $GP_{F0F1}$ | $GP_{F0F2}$ | $\Delta GP$ | $E_{F0F1}$ | $E_{F0F2}$ | $\Delta E$ |
| -3.636 ** | -2.486 * | -5.419 ** | 0.467 | -3.721 ** | -1.85 |

**Table S5. Multiple linear regression predicting reaction times.** Mean effect sizes (group T-values) of each regressor included in the multiple linear regression predicting reaction times (RT) in the partner selection task. Associated with **Fig.S4**. **$p<0.001$, *$p<0.005$, n.s. $p>0.1$.

| | | HC right | HC left | EC right | EC left |
|---|---|---|---|---|---|
| Z value<br>Wilcoxon signed rank test | $\tau$E vs. $\tau$D1 | 4.01 ** | 4.01 ** | 3.91 ** | 3.63 ** |
| | $\tau$E vs. $\tau$D2 | 4.01 ** | 4.01 ** | 3.28 ** | 3.42 ** |
| T values<br>Paired t test | $\beta$E vs. $\beta$D1 | 36.74 ** | 34.17 ** | 23.25 ** | 22.26 ** |
| | $\beta$E vs. $\beta$D2 | 50.08 ** | 37.48 ** | 21.27 ** | 25.18 ** |

**Table S6. The pattern similarity in the HC and EC activity is better accounted for by the 2-D Euclidean distance (E) than either 1-D distance alone.** First, we compared the rank correlation of E ($\tau$E, Kendall's $\tau_A$) with the rank correlation of the 1-D rank difference in the competence dimension ($\tau$D1) and the rank correlation of the 1-D rank difference in the popularity dimension ($\tau$D2) (top rows). The z-values of the two-tailed Wilcoxon signed-rank test are reported in the table (** $p_{FWE}<0.001$ in all 4 ROIs, Holm-Bonferroni correction). Second, we inputted the z-scored 2-D distance (E) and 1-D rank distance in one of two social hierarchy dimensions (D1 or D2) into the same general linear model (GLM) to pit them against one another to explain the pattern dissimilarity in the HC and EC (bottom rows). As regressors, we inputted E and D1 into one GLM, and E and D2 as regressors in another GLM. We further compared the regression coefficients $\beta$ with the paired t-test. Consistent with the Wilcoxon signed-rank test, we found that E explains the pattern dissimilarity in bilateral HC and bilateral EC significantly better than D1 or D2 alone (** $p_{FWE}<0.001$ in all 4 ROIs, Holm-Bonferroni correction). The t-values are reported in the table.

**a.**

| Brain areas | Laterality | k | T | Peak coordinate (MNI) | | |
|---|---|---|---|---|---|---|
| | | | | x | y | z |
| Supramarginal Gyrus | L | 271 | 5.74 | -42 | -46 | 26 |
| Caudate | L | 829 | 5.7 | -14 | 4 | 18 |
| Hippocampus | R | 1704 | 5.35 | 42 | -20 | -6 |
| Medial Prefrontal Cortex/ Rostral Anterior Cingulate Cortex | R/L | 658 | 4.66 | 0 | 30 | 10 |
| Orbitofrontal cortex (BA11) | R | | 3.54 | 26 | 48 | -10 |
| Hippocampus ❊ | L | 49 | 3.79 | -30 | -36 | -6 |
| Retrosplenial Cortex | R/L | 265 | 3.95 | 0 | -36 | 26 |
| Medial Prefrontal Cortex | L | 153 | 3.86 | -18 | 30 | -2 |
| Entorhinal Cortex ❊ | L | 24 | 3.81 | -18 | -10 | -26 |
| Entorhinal Cortex ❊ | R | 6 | 3.69 | 26 | -8 | -44 |

**b.**

| Brain areas | Laterality | k | T | Peak coordinate (MNI) | | |
|---|---|---|---|---|---|---|
| | | | | x | y | z |
| Hippocampus ❊ | R | 370 | 6.86 | 30 | -12 | -24 |
| Hippocampus ❊ | L | 164 | 4.44 | -26 | -10 | -28 |
| Posterior Cingulate Cortex/ Precuneus/ Retrosplenial cortex | R/L | 5301 | 6.33 | 22 | -68 | 38 |
| | | | 5.28 | 10 | -50 | 48 |
| Medial Prefrontal Cortex/ Rostral Anterior Cingulate Cortex/ Orbitofrontal cortex | L | 2063 | 5.82 | -14 | 38 | -6 |
| | | | 4.5 | 12 | 56 | 2 |
| Superior Frontal Sulcus | L | 32 | 4.65 | -24 | 0 | 44 |
| Inferior Parietal Lobule/ Temporoparietal Junction | R | 324 | 4.58 | 56 | -62 | 36 |
| Caudate | R | 75 | 4.44 | 14 | 22 | -2 |
| Entorhinal Cortex ❊ | R | 47 | 4.69 | 28 | -12 | -30 |
| Entorhinal Cortex ❊ | L | 53 | 4.32 | -28 | -14 | -32 |

**Table S7. Whole-brain searchlight representational similarity analysis (RSA). a.** Associated with **Fig.2f**. Brain areas in which its pattern dissimilarity was significantly explained by the pairwise Euclidean distances of 14 individuals on the 2-D social hierarchy (**Fig.2a**). The dissimilarity between activity patterns associated with each of 14 individuals from all events (F0, F1, and F2 presentations) was estimated from each searchlight while matching the number of observations of each individual (down-sampling). **b.** Associated with **Fig.2g**. Same as in (a.) except RSA is based on all observations acquired from all events (F0, F1, and F2 presentations). Results are reported at the threshold, $p_{FWE}$ <0.05 whole-brain TFCE correction except for the HC and EC (marked with ✻) which was tested at the threshold, $p_{FWE}<0.05$ TFCE corrected in the independently defined *a priori* ROI.
